# Integrating fish biology, population genomics and fishery policies for informed management of interjurisdictional multi-species fisheries

**DOI:** 10.1101/2024.07.12.603026

**Authors:** Mayuresh Gangal, Rohan Arthur, Venkatachalam Suri, Awadhesh Pandit, Abhinav Tyagi, Praveen Prakash, Mihir Sule, Iravatee Majgaonkar, Jagdish Krishnaswamy, Uma Ramakrishnan

**Affiliations:** Manipal Academy of Higher Education, Manipal, India; Oceans and Coasts programme, Nature Conservation Foundation, Mysore, India; National Centre for Biological Sciences, Bengaluru, India; Faculty of Biology, University of Gdansk, Poland; Ashoka Trust for Research in Ecology And the Environment, Bengaluru, India; School of Environment and Sustainability, Indian Institute for Human Settlements, Bengaluru, India

**Keywords:** IRIIS, Interjurisdictional fisheries management, Tropical multi-species fisheries, Genetic population structure, Fisheries genomics

## Abstract

Globally coastal fisheries are managed according to political jurisdictions of nations and their provinces. This practice often results in multiple states/provinces managing a single genetically distinct fish population autonomously and without coordination. In multi-species, decentralized and data poor fisheries such management lead to poor assessment and eventually overharvest of fish. To assess overexploitation risk for such interjurisdictional populations, we propose a risk indicator IRIIS which incorporates 1) species-specific vulnerability 2) distribution of genetically-distinct populations with respect to jurisdictional boundaries and 3) differences in management practices across jurisdictions. As a case study we evaluated overexploitation risk for individual populations of nine commonly caught fish species spanning across multiple jurisdictions along India’s west coast. Our results indicate that interjurisdictional differences in management alters innate biological susceptibility. IRIIS is also sensitive to population-level differences and policy changes. Our analyses emphasize the need for population-level assessments and interjurisdictional cooperation to achieve sustainable fisheries.

**Science and society statement:** Coastal fisheries worldwide are managed through politically defined maritime jurisdictions, while fish populations are structured by biological processes such as dispersal, ocean currents, and natural barriers. This fundamental mismatch between biological population boundaries and jurisdictional management boundaries often results in the same population being exploited under multiple, uncoordinated governance regimes. In the context of the global overfishing crisis, such governance fragmentation can lead to inaccurate stock assessments and heightened risk of local population collapse. This challenge is particularly acute in developing and tropical countries, where fisheries are multispecies, data-limited, and increasingly decentralized to support community-based governance. While decentralization promotes social justice, it can inadvertently intensify the misalignment between biological and administrative boundaries, complicating sustainable management.

In this manuscript, we introduce a robust framework — the Integrative Risk Indicator for Interjurisdictional Stocks (IRIIS) — which has a high applicability for multispecies, data-poor, and decentralized fisheries. Building on, and extending beyond, species-level Productivity Susceptibility Analysis (PSA), IRIIS integrates population genomics to delineate biologically meaningful population boundaries and combines these with fisheries policies and regulations across jurisdictions to quantify management-associated risks of overexploitation at the population level. We apply IRIIS to case studies of nine commercially important fish species harvested along the west coast of India. Our results demonstrate that the framework can (i) distinguish differential vulnerability among populations of the same species, (ii) explicitly capture management disparities across jurisdictions exploiting shared populations, and (iii) detect sensitivity to temporal changes in fisheries policies. By linking genomics, governance, and risk assessment, IRIIS provides a practical framework for identifying interjurisdictional management gaps and actionable entry points for policy coordination. This approach is particularly relevant for the Global South, where reconciling social equity objectives with ecological sustainability is both essential and challenging.

## 1. Introduction

Historically described as an inexhaustible natural resource ^1^, marine fish today face a crisis of overfishing ^2^. A recent report by the Food and Agriculture Organization of the United Nations flags 37.7 percent of global fish stocks as fished at unsustainable levels ^3^. By some estimates, ∼260 million people depend on marine fisheries for livelihood ^4^, while others suggest that ∼600 million people depend on fisheries and aquaculture at least partially or completely ^3^. These figures highlight that overfishing is not merely an ecological problem, but a crisis of food security, livelihoods and equity.

Since fish has been always viewed as a natural resource, when it comes to managing capture fisheries, economic and political factors override ecological factors ^5,6^. Also, management of marine resources is often done according to terrestrial concepts of management like sovereignty, boundaries, private property etc. Mismatch of these concepts with biological and ecological dynamics of marine ecosystems often impedes effective management of fisheries ^7–9^. Sustainably managing fishes at spatio-temporal scales that are biologically meaningful, economically viable and managerially effective is the principal challenge confronting global fisheries ^10^.

Single species demographic stock assessments provide most effective information for fisheries management ^10^. However, in 2012, 80% of global fish catch lacked formal assessment ^11^ and by 2020, only 25% of world fisheries were numerically assessed ^12^. The most reliable RAM legacy database also involves only 20% of world’s fisheries mostly coming from developed countries in temperate region ^13^.

For multi species tropical fisheries with limited capacities to undertake robust stock assessments, Productivity Susceptibility Analysis (PSA) can be a valuable tool. PSA attempts to evaluate overexploitation risk of multiple species simultaneously by calculating natural susceptibility of each species to resource offtake on a single comparable scale ^14–19^. They attempt to be biologically comprehensive, incorporating population growth rates, trophic level, breeding strategies and other crucial elements of fish biology in their assessments ^15^.

To bridge the gap between being an analytical useful tool to one that is directly applicable to fisheries policy, a few limitations of the PSA approach need to be addressed. It has been long understood that, for effective monitoring and management, genetically distinct populations of the same species should be considered as separate monitoring and management units ^20–22^. For most PSAs, the unit of analysis is the species and not individual population (but see Patrick et al. 2010 ^23^, limiting their reliability ^14,16–19^. Most PSAs also assume a uniform harvest regime along the entire geographical range of analysis. In most tropical countries with large coastlines like Brazil, Indonesia, Philippines, India and Mexico, coastal fisheries are often decentralized and governed by provinces which have significant autonomy in terms of fisheries management ^24–28^. Rarely is there a uniform harvest regime across the geographical area considered within a PSA.

Between the multiple genetically distinct populations of a species and decentralized harvest regimes, there is often a mismatch between the biological boundaries of fish populations and the jurisdictional boundaries of fisheries management. These mismatches lead to an increased risk of local population depletion ^29^.

Here we propose an Integrative Risk Indicator for Interjurisdictional Stocks (IRIIS) as a heuristic extension of the PSA by incorporating 1) the inherent resilience and susceptibility attributes of species characteristic of a typical PSA, 2) information from population genetics to determine geographical boundaries between populations, to indicate whether the genetically distinct population is managed by single or multiple management regimes. 3) the biological relevance of present management regulations on given species.

As a case study, we present a IRIIS assessment for nine commercially important nearshore fish species harvested along the west coast of India. Marine fisheries in India is a paradigm case of a multi-species multi-management fishery. In India, individual coastal states (provinces) have jurisdictional autonomy over adjacent territorial waters. They legislate fisheries regulations in these waters separately, based on local needs, demands from local stakeholder groups and with little dialogue with neighboring states ^28^. This management gives little regard to the underlying geographical spread of fish populations ^28^.

For undertaking IRIIS assessment, we first identified genetically distinct populations of nine chosen species along 3400 km stretch of the west coast of India. Then for each genetically distinct population, we estimated biological congruence management policies of single or multiple management regimes acting on them. This information was then used to construct a population-wise IRIIS. We then discuss the utility of this matrix in guiding policy and management decisions.

## 2. Results

### 2.1. Representing risk of overexploitation associated with species biology (y axis)

IRV values can range between 0 (most vulnerable) to 1 (least vulnerable) ^18^. Among the 133 species Mohammad et al. 2021 analyzed, *Solenocera crassicornis* (coastal mud prawn) had the highest IRV value of 0.512 where *Carcharhinus sorrah* (spot-tail shark) had lowest IRV value of 0.022. Among the nine target species analysed here, *Nemipterus japonicus* had the highest IRV index value of 0.31 and thus the lowest vulnerability to overexploitation on the vertical axis whereas *Epinephelus diacanthus* had the lowest IRV index value of 0.195 making it most vulnerable on the vertical axis. Using the approach outlined in Mohammed et al. 2021, we calculated an IRV index of 0.237 for *Sillago sihama* (Supplementary Material Section 1). Our target species occupied a narrower portion of the range of species analyzed by Mohammad et al. 2021.

### 2.2. Establishing geographical boundaries of target fish populations

We sequenced 311 samples belonging to nine study species across nine locations. After removing samples with low quality during the filtering process, we used a total of 273 samples for the final analysis. On average, we analyzed 4-5 samples per species per location (See figure S4 in supplementary material).

Single Nucleotide Polymorphism (SNP) analysis indicates that the degree of gene flow between individuals across the west coast of India varies considerably with species (Figure 1). The PCA results are presented in Figure 1. For detailed species-wise results which include results of pairwise F_ST_ and Admixture analysis refer to the section 3 of the supplementary material.

**Figure 1.**
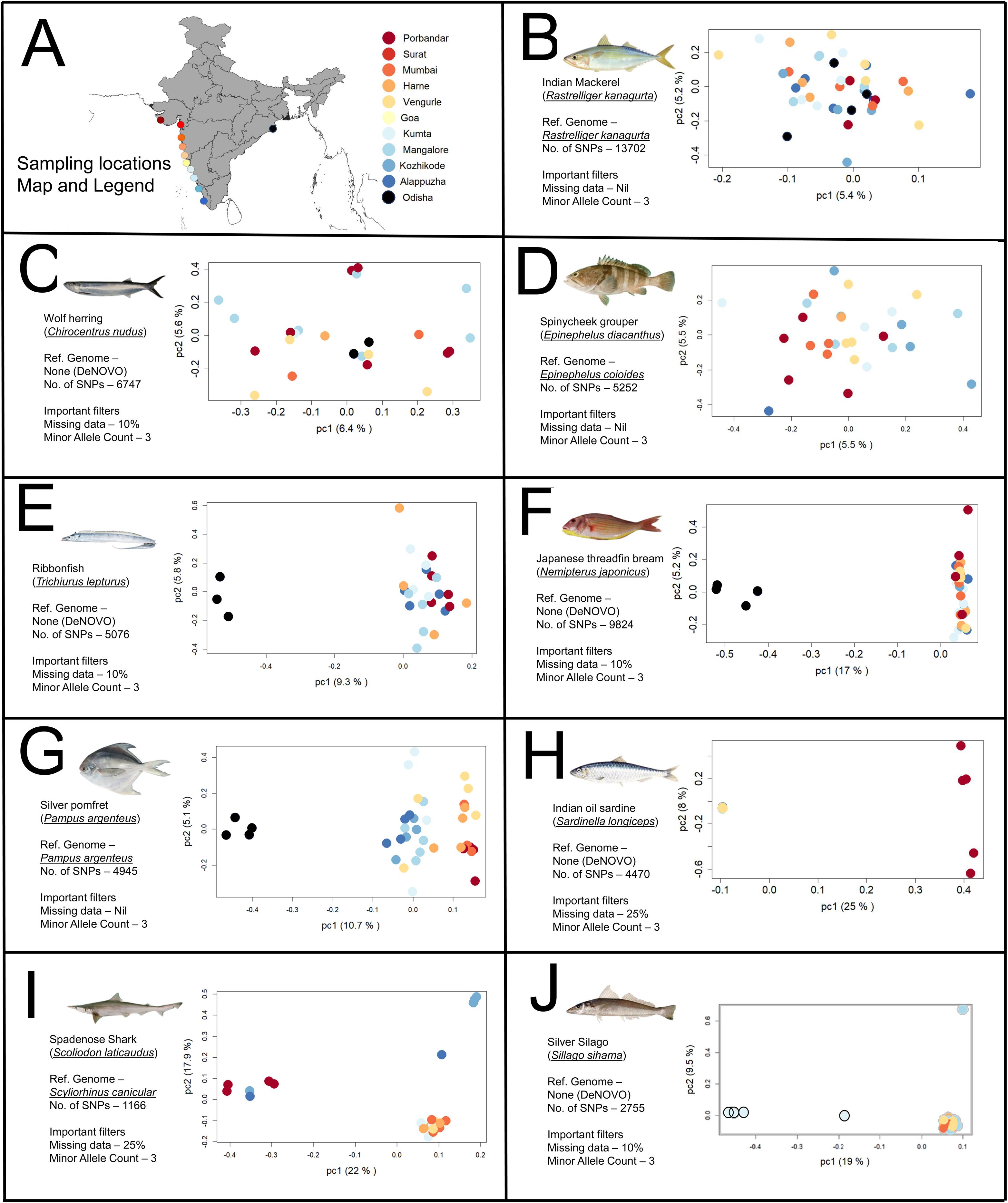
Principal Component plots for nine chosen species across the west coast of India. Panel A – Map of sampling location along the Indian coastline with the colour legend for samples from specific locations. Porbandar = Dark Red, Surat = Red, Mumbai = Light Red, Harne = Orange, Vengurle = Light yellow, Goa = Yellow, Kumta = Light Cyan, Mangalore = Light Blue, Kozhikode = Medium Blue, Alappuzha = Blue, Odisha = Black. Panel B to J – Principal Component Analysis plot of individual species with information about number of SNPs and important filters used for variant calling. See also figures S5 to S13 in supplementary material

*Rastrelliger kanagurta* and *Chirocentrus nudus* show no evidence of population differentiation across the west and east coast of India (Figure 1B & 1C). Due to lack of samples from the east coast, we restricted analysis of *Epinephelus diacanthus* to the west coast. *Epinephelus diacanthus* shows evidence of a single population across the entire west coast (Figure 1D). *Trichiurus lepturus* and *Nemipterus japonicus* also show a single population across the entire west coast, however they have clear divergences from conspecific populations from the east coast (Figure 1E & 1F).

The remaining four species (*Pampus argenteus, Sardinella longiceps, Scoliodon laticaudus, Sillago sihama)* show multiple genetically distinct populations along the west coast of India. Genetic data indicate two potential *Pampus argenteus* populations along the west coast of India, one between northern Gujarat to central Maharashtra, and another from southern south Maharashtra to Kerala (Figure 1G). *Sardinella longiceps* also shows evidence of two populations, with a Gujarat population that separates clearly from the rest of the west coast population (Figure 1H). Genetic data from *Scoliodon laticaudus* shows evidence of three distinct west coast populations: a northern Gujarat population, a middle peninsular (Maharashtra-Goa-Karnataka) population, and a southern Kerala population (Figure 1I). Finally, *Sillago sihama* shows evidence of an isolated population near the town of Kumta which is separate from the coastwide Maharashtra - Karnataka population (Figure 1J).

### 2.3. Identifying percent disparity between species biology and regulations

We identified the percent disparity between biology and regulation for each individual population - state (management regime) combination based on the rationale described in the methods section 2.1.2. To clearly demonstrate how disparity was calculated, results of the disparity analysis for the *Sardinella longiceps* are shown in Figure 2 as a model species. In *Sardinella longiceps* The Gujarat population (which is housed in a single management regime) has a high disparity (66%), while the peninsular India population, despite being shared across four states, has a low disparity across the entire coast. Further results of all species are provided in the Supplementary Material, Section 2.

**Figure 2.**
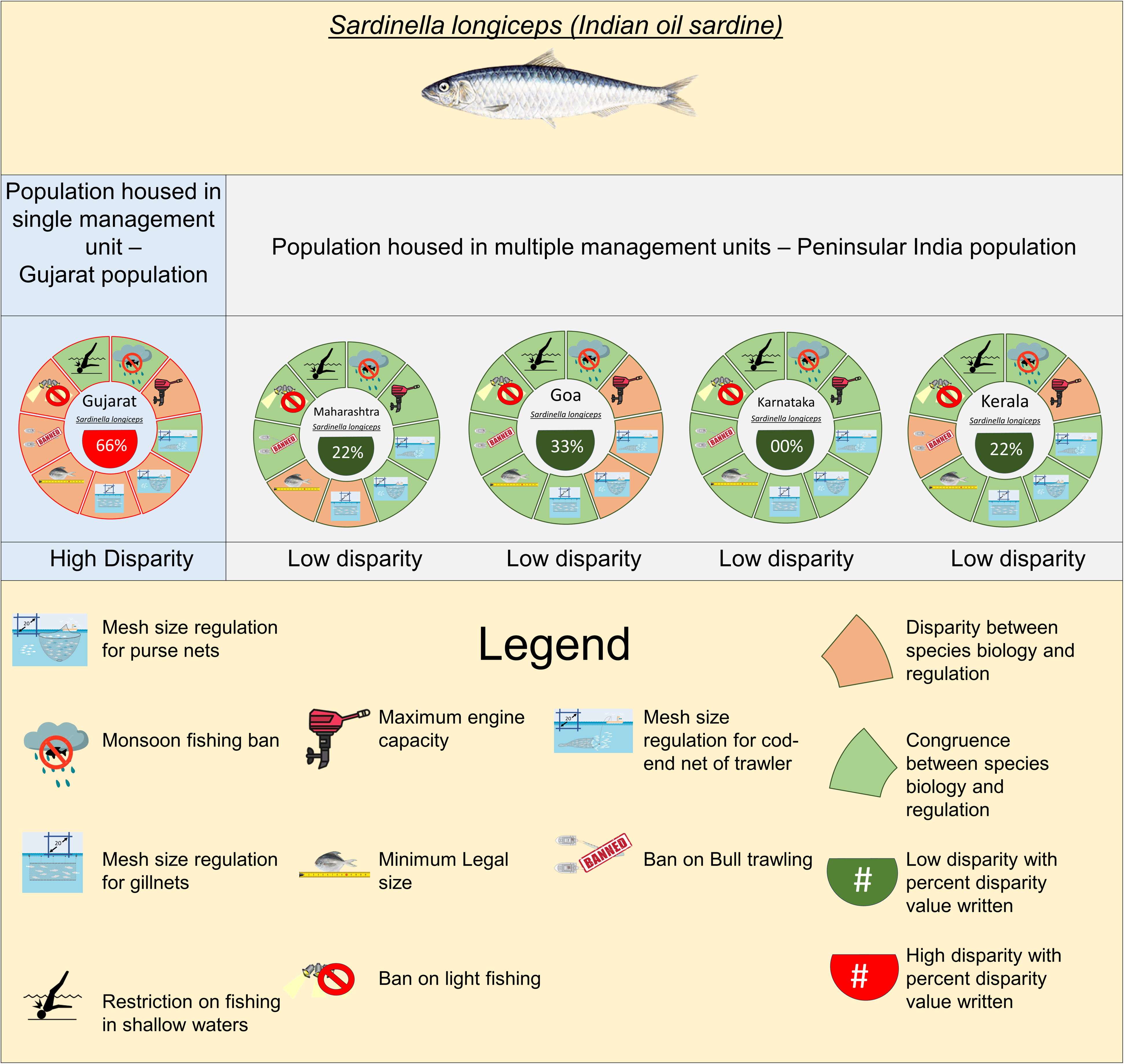
Disparity analysis for Indian oil sardine, with populations housed in single as well as multiple management regimes. Each donut represents a state along the west coast of India. Nine sections are for nine different regulations and the color of each section represents congruence or disparity of the sardine biology with the regulation. Legend for all the symbols and colors are embedded in the figure. See also figure S2 in supplementary material

### 2.4. Populating the vulnerabilities of west coast populations using IRIIS

We used species specific IRV values from Mohammad et al. 2021 ^18^ and state-wise percent disparity for each population identified by genetic analysis, to construct a IRIIS for the west coast (Figure 3). variations in individual populations of different species were evident across both the axis of IRIIS. Critically, species with multiple populations along the coast could have varied overharvest risk on the management disparity axis. This was most evident for *Scoliodon laticaudus*, where based on the disparity scores, the peninsular population had a much higher vulnerability than the Kerala and Gujarat populations. For populations of *Pampus argenteus* and *Sardinella longiceps*, differences in management practices between states still clearly influenced the vulnerability of each population. Widely ranging species (with single populations across the coast) had very different IRV values and were thus very different in their biological susceptibility. However, because their populations spanned several jurisdictions along the coast, they were all much more prone to management disparities.

**Figure 3.**
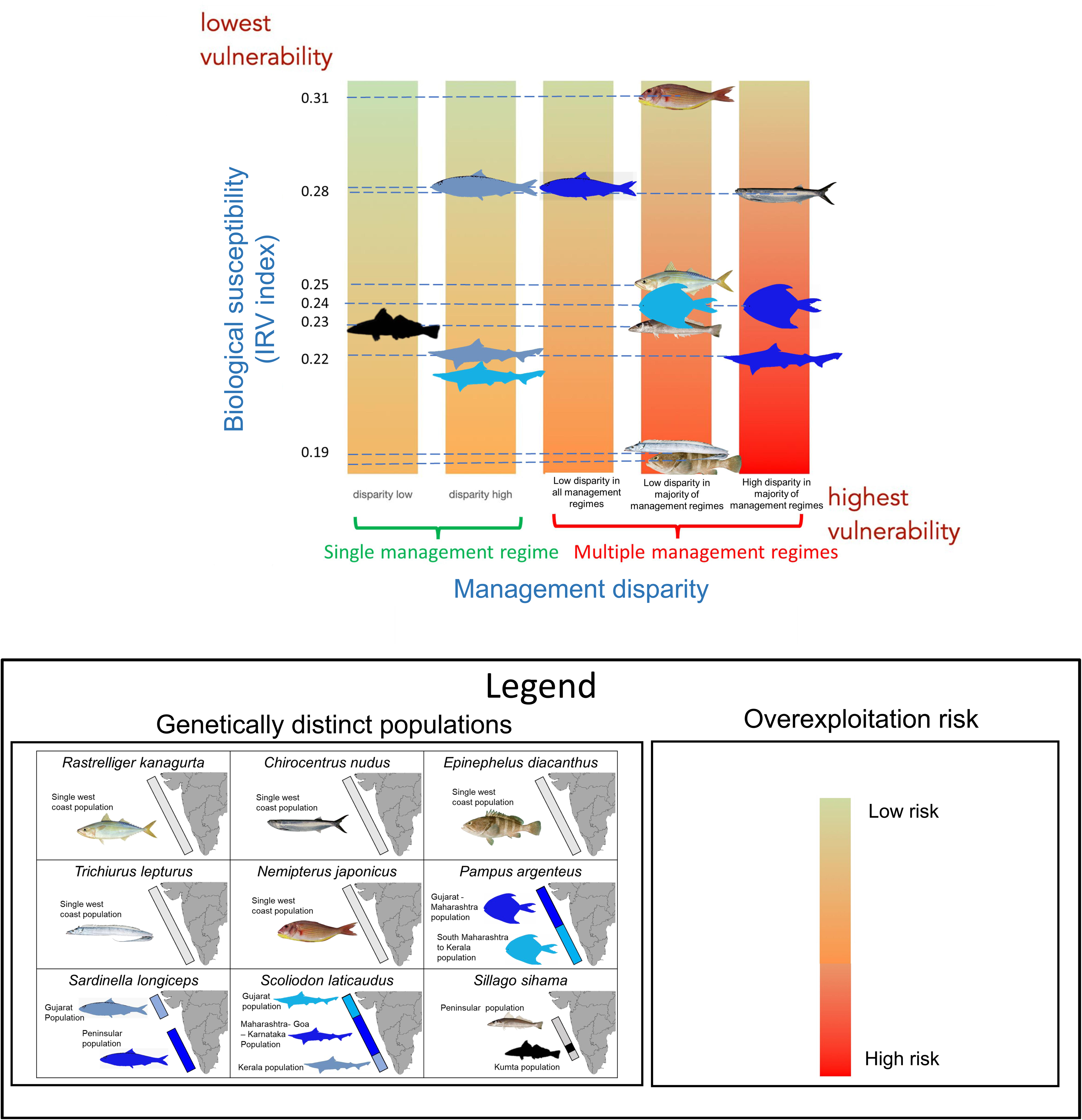
Integrative Risk Indicator for Interjurisdictional Stocks (IRIIS). for nine species along the west coast of India. Five colored bars represent five risks associated with management disparity which is increasing from left to right. vertical axis of each bar represents IRV score. For species presented on the matrix the IRV score is written on the left-hand side. The color gradient on each vertical bar represents overall risk of overexploitation. Different population of each species is indicated using maps in legend section. See also figure S2 in the supplementary materials

## 3. Discussion

Achieving sustainability in multi-species, poorly monitored and data-poor tropical coastal fisheries is often considered a wicked problem with no easy solutions ^30^,. Although recent approaches like the PSA comprehensively incorporate complex species biology, they do not account for management complexities arising from the interjurisdictional nature of most coastal fish populations. IRIIS combines genomic techniques and policy analysis for a more realistic assessment of fish stock vulnerability.

Our case study demonstrates that populations of many commonly targeted coastal species across the Indian west coast are interjurisdictional in nature, spanning multiple harvest regimes and fisheries policies. This has critical implications for tropical fisheries management. Our study demonstrates that within a species, overharvest risk for individual populations varies based on which and how many management regimes are harvesting them. By capturing these complexities IRIIS provides a framework which allows fisheries managers to evaluate different fisheries options and negotiate pragmatic interjurisdictional arrangements based on known boundaries of at-risk species.

### 3.1. A robust approach to predict vulnerability to overharvest

Our analysis demonstrates the vital importance of evaluating overexploitation risk for every population individually, something that most PSA assessments do not allow ^14,16,17,19^. A case in point is the *Pampus argenteus*. With an IRV score of 0.24 (based on the IRV index, see Methods), it is a species of intermediate risk. Our population genetic analysis shows that there are two populations of *Pampus argenteus* along the west coast of India, a northern population that extends from Gujarat to the central coast of Maharashtra, and a southern population that extends further south to Kerala. Although both of these populations share the same IRV score, they are differentially susceptible to overharvest. Kerala have regulations on minimum mesh size for gill nets, and there are *Pampus argenteus* -specific minimum legal-size regulations in place in Goa, Karnataka and Kerala ^31–33^. These regulations are absent in the northern states (refer table S2 in supplementary material). In other words, the southern population (southern Maharashtra to Kerala) is managed by states with harvest rules more congruent to *Pampus argenteus* biology (low disparity), making it much less vulnerable to overfishing than the northern population (Gujarat to central Maharashtra). This example demonstrates how by shifting the assessment focus to spatially explicit populations, our matrix considerably enhances the value of the original IRV assessment. Further, by identifying the specific management attributes responsible for differences in risk, the matrix provides actionable recommendations to policy makers on how to reduce overharvest risk with changes in policy and interjurisdictional cooperation. While the matrix is unable to assess how these management efforts perform on the ground, but provides a useful framework to assess their effectiveness.

Another aspect of this IRIIS is that it is dynamic and sensitive to periodic revisions in fishery policy across management regimes. The case of *Sardinella longiceps* demonstrates this well. The west coast has a largely panmictic population except for Gujarat, which has a distinct population. Until 11 years ago, no west coast Indian states had regulations on fishing methods like bull trawling, light fishing etc ^31,34–36^. In addition, there were no mesh size regulations for purse nets or gill nets and none of the states had defined minimum legal sizes for the *Sardinella longiceps*. As a result, based on IRIIS, until 2016, the *Sardinella longiceps* was in the highest category of interjurisdictional management associated overharvest risk. The subsequent adoption of these regulations by all west coast states has significantly reduced this risk and moved *Sardinella longiceps* on the management axis from “High disparity in majority of the management regimes” to “Low disparity in all management regimes” category (Figure 4). Along with the sensitivity of the matrix, this example shows how the matrix can be used as a tool to formulate biologically meaningful regulations / policies for target species. In addition, the multi-species nature of the matrix can help policy makers to track what impact the new regulations will have on other non-targeted species.

**Figure 4.**
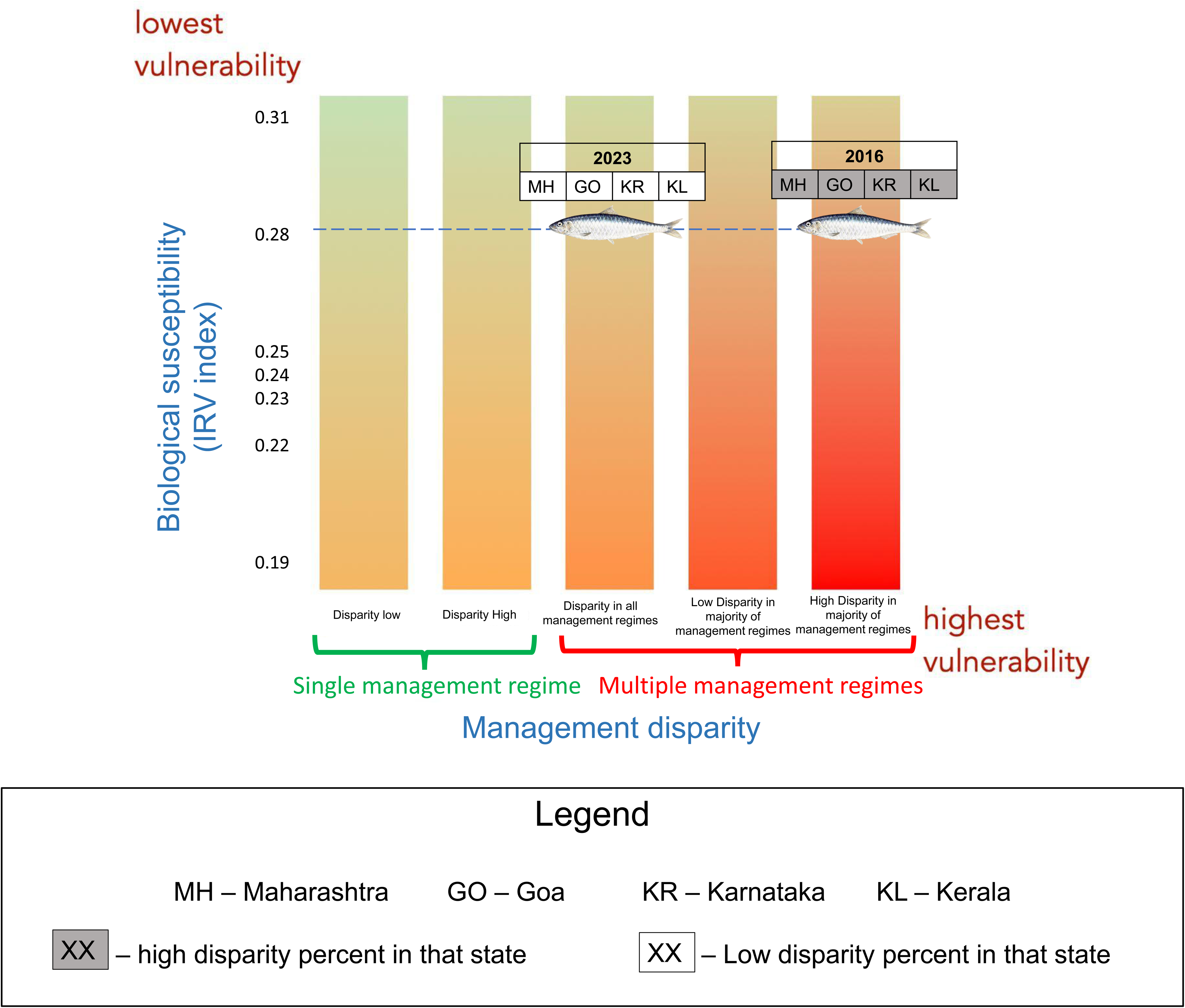
Shift in overexploitation risk of the peninsular population of *Sardinella longiceps* along the management disparity axis as a direct result of changing management practices. Five colored bars represent five risks associated with management disparity which is increasing from left to right. Vertical axis of each bar represents IRV score. For species presented on the matrix the IRV score is written on the left-hand side. The color gradient on each vertical bar represents overall risk of overexploitation. See also table S2, S3 and S5 in supplementary materials

We assume that IRV values which are based on critical life history traits remain relatively constant for a species. However, it is possible that these traits could vary between genetic stocks. For instance, Patrik et al., 2010 show that the PSA vulnerability of different populations of the same species could differ considerably ^23^. In evaluating the disparity between biology and management regime information for our nine target species, we identified considerable differences in basic life history characters like length at first maturity (L50) between *Pampus argenteus* ^37,38^ and *Scoliodon laticaudus* ^39,40^ caught at different locations. These traits could potentially also influence their catchability in gear operating along the coast. Such differences could potentially lead to varying inherent population-level susceptibilities and slightly different IRV values for each species. One benefit of using matrix like IRIIS is that as it incorporates genetically distinct population in risk assessment actually makes these regional differences in life history more apparent.

While IRV values may vary slightly as we learn more about each population, they represent a largely unchanging inherent property of each taxon. However, a central feature of our matrix is that real-world vulnerability is as much a characteristic of the management jurisdictions in question as it is of the biological character of the species. As an example, the IRV value for *Chirocentrus nudus* is high, making it apparently less vulnerable to overexploitation. However, its population is spread all along the west coast which have five different management regimes of five states. In addition, the regulation of four of the five states has very low congruence with *Chirocentrus nudus* biology (higher disparity). This lack of congruence makes it far more vulnerable to overexploitation compared to its species-associated vulnerability indicated by IRV. This explicitly recognizes the fact that fisheries regimes vary widely between locations, and that transboundary agreements (or the lack of them), can strongly mediate the inherent vulnerability of fish stocks. It is critical to keep this in mind when applying the matrix to a specific fishery. For instance, we classified the Gujarat population of the *Sardinella longiceps* as belonging to a single jurisdiction, reducing its overall vulnerability within the matrix. Broadening the geographical scale of our analysis may well have shown that this population is contiguous with *Sardinella longiceps* from Pakistan and beyond, making it potentially subject to trans-national management differences that we did not evaluate here. More comprehensive analysis would require establishing genetic boundaries for the entire range of the species in question before determining its overall overharvest risk. This will have important implications for the kinds of interjurisdictional arrangements required to reduce the overall susceptibility of the population – between coastal nations or between sub-national jurisdictions ^41,42^.

The fact that management differences can so strongly mediate inherent vulnerability indicates that assessments of this kind are most meaningful when designed for every region, making them jurisdictionally and biologically appropriate across the range of the species.

Productivity Susceptibility Analyses or PSAs are limited in their scope of use since they cannot incorporate demographic information and population trends ^43,44^. Single species stock assessments and Bio-economic harvest models based on Maximum Sustainable Yield (MSY) are limited in applicability because of focus on single species ^45,46^. Although the Kobe/Kobi plot incorporates demographic information of multiple species simultaneously it is considered as too complex from the point of view of policy ^47^. IRIIS approach increases applicability of all such assessments by providing a usable framework for policy makers. It incorporates the PSA approach on one axis to incorporate life history-based susceptibility of species. It incorporates policies of individual management regimes and disparity between them. If a single species stock assessment shows a decreasing trend in stocks, it helps to identify potential policy actions and assess how other non-species will react to such policy changes.

The central condition of the IRIIS approach is identifying genetically distinct management units. Larkin highlighted the need of defining stocks based on genetically distinct populations 45 years ago^49^. Multiple studies since them have also demonstrated various issues fisheries face in the absence of this information^29,50^. Even so majority of developing countries operate without this information. We believe our work highlights the importance of identifying genetically distinct populations on both international and regional scales. The techniques we describe in this study offer a relatively cost-effective means of establishing the spatial limits of fish populations. This could inform more nuanced population-level life-history studies and improved risk assessments.

### 3.2. Trans-jurisdictional management of fish populations in tropical waters

How could this or exercises such as IRIIS inform sustainable tropical fisheries? Risk assessments of this sort go a long way in sharply defining the management needs for each population across its natural range and help define the scale at which fisheries management would be most biologically meaningful. It offers a clear set of directions to better align species biology with management, setting the stage for cross-jurisdictional cooperation on specific aspects of policy and practice.

Although we focused on populations which span across multiple jurisdictions within a sovereign nation, the matrix can be applied for trans-boundary situations as well. Countries in western Europe have had a long history of hard-won transboundary fishing arrangements ^48^, driven by decades of conflict over shared resources. There is also substantially better information on geographically identifiable, genetically distinct populations of fish that form the basis of these arrangements ^50^. For some tropical countries transboundary agreements are important means of earning revenue ^51–53^. Engaging in such agreements knowing the geographical extent of genetically distinct populations of transboundary fish and how these species-specific agreements impact other species in broader multi species fisheries, can increase the potential effectiveness of such agreements.

In many countries coastal fisheries are managed in a highly decentralized manner where regions and provinces have considerable autonomy over local fisheries management ^24–27,54^. In addition, the allied problems of equity and social justice associated with coastal fisheries are seeing an increasing movement towards even further localization of fisheries, with approaches like TURFs (Territorial User Rights for Fishing) gaining popularity ^55^. These are usually smaller than regional government fisheries management units. Most TURFs in Chile, Mexico and Japan are smaller than the dispersal ability of fish larvae and adults ^56^. While these efforts are a welcome direction to secure the livelihood of marginalized coastal communities, they are also subject to boundary paradoxes, where increasing fisheries boundaries come at the cost of increasingly complex interjurisdictional negotiations ^57^. Our analysis raises serious questions of how to ensure the success of these pockets of community managed fishing in a wider ocean of a rapidly expanding fishery sector ^58^. Without adequate coordination across biological bounds, these efforts may also result in worsening of current rates of overharvest, and it is important to create forums of dialogue between these community managed and larger regional fisheries, keeping in mind the genetic boundaries of fish stocks.

As far as our case study from India is concerned, recent Indian fisheries policies heavily emphasize ecosystem-based management of fisheries. The vision statement of the 2020 National Fisheries Policy emphasizes “developing an ecologically healthy, economically viable and socially inclusive fisheries sector” with “facilitating coordination among States/UTs, National Agencies and other stakeholders” as its major policy intervention ^59,60^. This policy can create a perfect platform to incorporate spatial information of genetically distinct populations of individual fish species in the fisheries policy framework. So far, no forum has been created to facilitate coordination among states and union territories. We hope our results can provide a strong reason to start this coordination process.

### 3.3. Conclusion

For multi-species tropical fisheries, transitioning to population-specific management of fish stocks may seem like a distant horizon. Eventually though, if coastal tropical nations are serious about stemming the collapse of coastal fish stocks, population-wise management is likely to be critical, at least for the most vulnerable of coastal species. Coastal interjurisdictional arrangements of the kind that are already in place for open-water migratory species like tuna ^61^ offer a possible template that can be built on. Our results demonstrate the need for such agreements for nearshore species of fish and for individual provinces and community managed areas within a single nation state, which was largely considered less important in tropical context. For this, IRIIS can be a useful prioritization tool. Because the management axis allows for dynamic mobility based on changes in policy, it can be used to strategically evaluate which set of policy directions could reduce the susceptibility of the widest number of fish populations across a shared coastline. As our analysis shows, the inherent vulnerability of fish can be strongly mediated by what managers do at local levels, and how closely they coordinate their actions with neighbors that share the same genetic resource pool. This offers the principal challenge but also the best hope for rational management of dwindling fish stocks.

## 4. Methods

We first describe – the Integrative Risk Indicator for Interjurisdictional Stocks (IRIIS) framework to assess the relative Risk of genetically distinct, interjurisdictional fish stocks based on an extension of a standard PSA. We then demonstrate the utility of this framework based on a case study of nine nearshore species along the west coast of India.

### 4.1.1. Constructing the Integrative Risk Indicator for Interjurisdictional Stocks (IRIIS)

We propose that the risk of overharvest for the population of a target species is a function of i). the inherent life history and catchability of each species (PSA) and ii). the biological sustainability of management practices across all the jurisdictional regimes the population spans. Figure 5 outlines the rationale IRIIS assessment. The detailed description can be found in 4.1.1.a and 4.1.1.b

**Figure 5.**
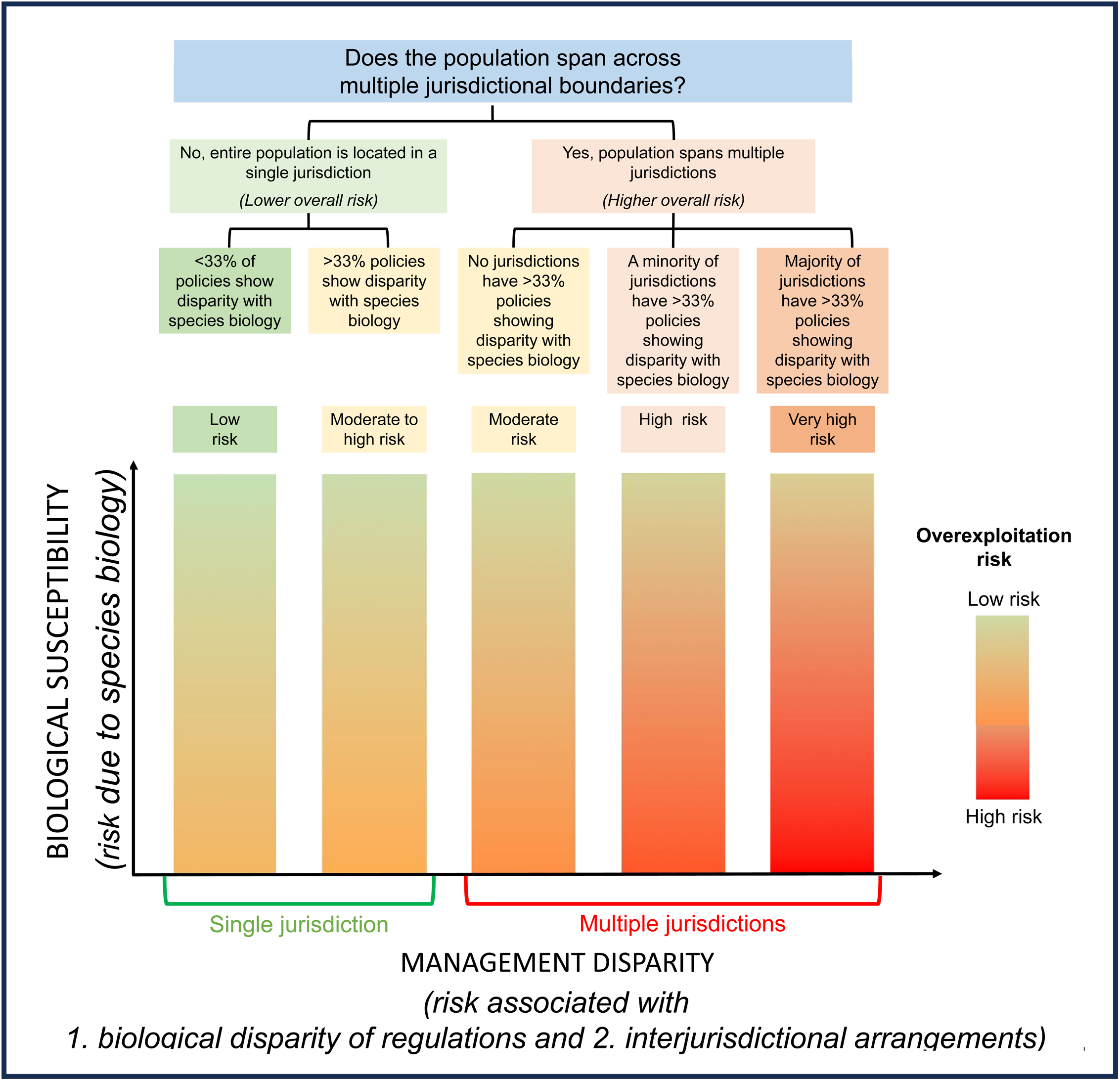
Rationale of development of Integrative Risk Indicator for Interjurisdictional Stocks (IRIIS). The matrix uses indicators like PSA analysis as a starting point to denote biological susceptibility (represented on the y axis). The management disparity axis (x axis) is based on an ordered set of decision rules that evaluate the jurisdictional overlap of each population, and the congruence of management practices in each jurisdiction with known species biology. Five colored bars represent five risks associated with management disparity which is increasing from left to right. The color gradient on each vertical bar represents overall risk of overexploitation.

#### Rationale for constructing the IRIIS

##### 4.1.1.a. Vertical axis of biological susceptibility

The vertical axis of the Integrative Risk Indicator for Interjurisdictional Stocks (IRIIS) is the score assigned by classical Productivity Susceptibility Analyses which typically assign a vulnerability score to their sampling units (usually species) ^16,18,23^.

##### 4.1.1.b. Horizontal axis of management disparity

The horizontal axis of the matrix is premised on the observation that in addition to species-based vulnerability indicated by PSA analysis, the vulnerability of a particular population is mediated by the degree to which the spatial extent of the population matches the spatial extent of the management regime. Therefore, populations can be grouped in two broad vulnerability categories i) Populations located entirely within a single management jurisdiction ii) Population ranging across multiple management jurisdictions. Note that the species-specific vulnerability (based on PSA) does not change in these assessments between populations.

If the assessed population is entirely located within a single jurisdiction, we classify its management-based vulnerability by calculating the percent disparity: Low disparity <33%; High disparity >=33%. Management-based vulnerability increases as we move from bar 1 to bar 2.

If a population spans multiple management regimes, we assess its interjurisdictional management vulnerability by calculating the percent disparity of the population within each jurisdiction separately. Based on the proportion of jurisdictions with more than 33% disparity with species biology we assign the population in three assessment categories. 1) All jurisdictions with <33 % disparity (vertical bar no. 3 in Figure 5), 2) majority of management regimes with <33% disparity (vertical bar no. 4 in Figure 5), 3) Majority of management regimes with >33% disparity (vertical bar no. 5 in Figure 5). Management-based vulnerability increases from left to right.

We assume that populations entirely located in a single management jurisdiction overall have an inherent lower vulnerability than populations spread across jurisdiction of multiple management regimes due to inherent differences in implementation between jurisdictions and the added complications of interjurisdictional co-ordination, if any.

### 4.1.2. Evaluating the biological relevance of present management regimes

For every management regime within the geographical area of the assessment, we first catalogue all available policy instruments, regulations or subsidies related to the fishery. We then evaluate each of these practices and policies based on their congruence (or disparity) with sustainability goals based on what is known of species’ biology. E.g. A policy to restrict mechanized fishing during June-August, is biologically congruent with species that breed during this period. Conversely, the policy would show biological disparity with species that do not breed during this period. The proportion of all policy/practice attributes in that management regime that do not align with the biology of the given species is used to calculate a percent disparity score.

Table 1 demonstrates how we calculated the percent disparity for a combination of a hypothetical province with only two policies related to harvest and a hypothetical species which breeds between June and August and has Length at Maturity (L50) of 20 cm. For simplicity, in the current assessment, we assume that all attributes have an equal weightage on sustainable harvest. However, it is quite possible to assign different weights to each attribute if there is a sufficient justification to do so. Weightage can also be assigned based on reliable knowledge of how effectively each policy attribute is implemented on the ground.

**Table 1.** Evaluating the biological efficacy of management practices for a hypothetical province with only two policies related to harvest and a hypothetical species which breeds between June and September, and has Length at Maturity (L50) of 20 cm

| Policy Number | Attribute and its intended consequence | Prescribed policy | Species biology | Assessment |
| --- | --- | --- | --- | --- |
| 1 | Temporal/seasonal ban on mechanized fishing to protect breeding individuals | Ban period: June – August | Breeding period of target species: June- September | Congruence between species biology and policy attributes |
| 2 | Prescribed Minimum Legal Size (MLS) to protect juveniles from overfishing | No prescription (all size individuals are legal to catch) | Length at Maturity (L50) – 20 cm | Disparity between species biology and policy attribute |
|  | Total disparity percentage |  |  | 50% policy attributes show disparity – **High disparity combination** |

### 4.2. Case study: West coast of India

We apply IRIIS to evaluate the vulnerability of populations of nine commercially important fish species along the west coast of India. We use SNP-based genomic data to identify species population boundaries for these target species and an analysis of management practices across all regimes to populate the matrix.

#### 4.2.1. Study area

The study region extends from the western shores of the Kutch peninsula to Kanyakumari at the southern tip of India - a stretch of approximately 3400 km of coastline (Figure.6) The region represents a single eco-region based on its faunal community composition ^62^. The governance of the coast is distributed across five states (Gujarat, Maharashtra, Goa, Karnataka and Kerala). Each state regulates its own fisheries with policy instruments in the form of state specific Marine Fisheries Regulation Acts (MFRAs). Such disjunct governance in a single ecoregion makes the west coast an ideal case study to understand how differences in fisheries management can mediate overharvest risk of individual fish populations.

#### 4.2.2. Choice of species

Indian fisheries are typically multi species, with several species constituting the target catch. For such fisheries, the comparison between biological boundaries of fish and the jurisdictional boundaries of fisheries needs to happen for multiple species simultaneously. Our attempt was to include as many fisheries target species as was logistically possible.

We identified nine fish species commonly caught in trawl fisheries along the west coast of India. The choice of species was dependent on their ubiquity, the frequency with which they occur in trawl catches across the west coast, and ease of identification in the field. They were also chosen to represent different trophic groups, habitats and taxonomic groups.

The selected fish species were silver pomfret (*Pampus argenteus*), ribbonfish (*Trichiurus lepturus*), Japanese threadfin bream (*Nemipterus japonicus*), spiny cheek grouper (*Epinephelus diacanthus*), spade nose shark (*Scoliodon laticaudus)*, wolf herring (*Chirocentrus nudus*), Indian oil sardine (*Sardinella longiceps),* ladyfish (*Sillago sihama*), and Indian mackerel (*Rastrelliger kanagurta*).

#### 4.2.3. Calculating risk of overexploitation associated with species biology

Mohamed et al. in 2021 ^18^ formulated an Index of Resilience and Vulnerability (IRV) based on the growth coefficient (K), longevity: asymptotic length (L∞), reproductive load, number of spawning months (NSM), fecundity and mean trophic level (MTL) as vulnerability attributes and species geographic distribution, body length depth ratio (BLD), exploitation ratio, recruitment, susceptibility to gear and price as resilience attributes. Although not explicitly framed as PSA, the IRV uses same basic principles. This is the most rigorous assessment of species susceptibility of marine fish from the region. we used it to formulate our y axis which represents risk of overexploitation associated with species biology.

We used IRV values for all nine species (Figure 3). Mohammad et al. 2021 did not assess the IRV index of *Silago sihama*, so we calculated its IRV based on the formula given by Mohammad et al 2021 (Supplementary Material, Section 1).

#### 4.2.4. Identifying the geographical extent of individual fish populations

##### 4.2.4.1. Data collection

We sampled individuals of each species from ten sites located at roughly 200 km intervals along the entire west coast from Gujarat to Kerala. We have listed the sites with the map in Figure 6. At each site we located the places where fishing boats land their catch (landing sites) and we sampled freshly landed fish from these landing sites. Since the study uses genetic data to identify geographical boundaries between populations, we also sampled individuals from Balasore - a single location on India’s east coast. Balasore is located at least 1700 km away from southernmost sampling locations on the west coast and was expected to act as a separate population in the/our analysis.

**Figure 6.**
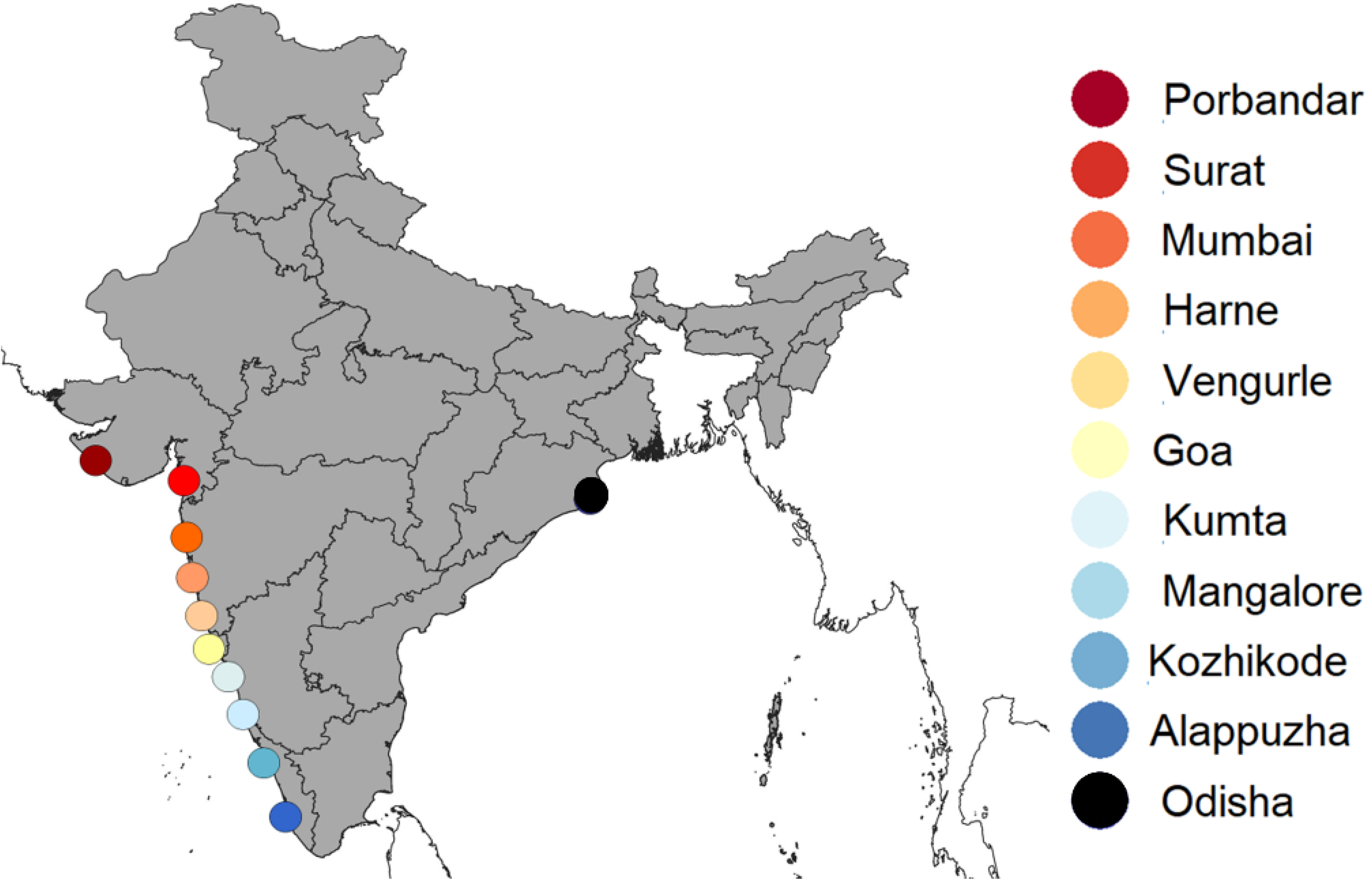
Location of sampling sites along the west coast of India. Map of sampling location along the Indian coastline with the color legend for samples from specific locations.

Tissue samples were collected using fin clips extracted from the fish directly at the landing site. Fin clips of 10-18 individuals from each species were collected at each sampling location using sterilized instruments to avoid cross-contamination, and stored in absolute ethanol in a 2 ml tube. We ensured that the fish sampled were sourced from local waters by sampling from boats where the sailors confirm that they were fishing in waters no further than 50 km from sampling location. As an additional safeguard we sampled only boats with lower engine capacity and cannot travel beyond 50-60 km from the landing site due to fuel constraints. Sampling fish without considering their life stages posed the risk of mixing individuals from two distinct populations, leading to potential misidentification. To avoid this, we collected samples from only the adult population at each location.

##### 4.2.4.2. Data generation

We used genome-wide SNP (Single Nucleotide Polymorphism) markers to identify genetically distinct populations and their geographical extent for all nine chosen species. We chose SNP-based methods to get a high number of regions of differentiation across the DNA to overcome the constraints of sample size ^64^. We used the Double Digest Restriction Associated DNA (ddRAD) approach to generate genomic data ^65^. The sample collection protocol along with detailed protocols of DNA extraction, library preparation is given in the Supplementary Material, section 3.

##### 4.2.4.3. Data analysis

We used three approaches for SNP calling: 1) For silver pomfret (*Pampus argenteus*) and Indian mackerel (*Rastrelliger kanagurta*), we used available reference genomes from the NCBI database for SNP calling (*Rastrelliger kanagurta -* GCA_024605255.1, *Pampus argenteus* - GCA_000697985.1). 2) For spiny cheek grouper (*Epinephelus diacanthus*) and spade nose shark (*Scoliodon laticaudus)* we used the reference genomes of the nearest taxonomically related species. We used the *Epinephelus coioides* (GCA_900536245.1) genome for *Epinephelus diacanthus* and *Scyliorhinus canicula* (GCA_902713615.1) genome for *Scoliodon laticaudus* from the NCBI database. 3) There were no available reference genomes that we could use for any of the other species, so we used a *de novo* variant calling pipeline by STACKS. The detailed bioinformatic protocol (quality control of reads and SNP discovery) are provided in Supplementary Material, section 3 and 4.

After SNP discovery, we used three indices to identify evidence of genetic similarity and dissimilarity between samples and ultimately identify genetically distinct populations for all nine species. These indices were:

1. Visualization based on a principal component analysis (PCA) using PLINK ^66^.
2. Visualization based on admixture probabilities using the program ADMIXURE ^67^. We ran an admixture for K ranging from 1 to n, where n = number of sites in which we have information of that particular species, for 10 replicates each, the best K was determined by estimating the cross-validation error. The results were analyzed and plotted using a web-based server, CLUMPAK ^68^.
3. A pairwise mean F_ST_ calculation conducted using STACKS ^69^.

#### 4.2.5. Identifying disparity between species biology and management attributes

Across the west coast of India, we examined fisheries regulations and policy instruments relevant to our study species. Following Gangal et. al ^28^, we identified nine different regulations as the key management attributes of the regimes prevalent in the five states of the Indian west coast ^28^. They were 1) partial or complete fishing ban during monsoon months (June-August) 2) maximum allowable engine capacity of a fishing vessel 3) minimum allowable mesh size for cod end of trawl operation 4) minimum allowable mesh size for gill nets 5) minimum allowable mesh size for purse nets / ring seine nets 6) minimum allowable legal size for a species 7) ban on bull / paired trawling 8) ban on fishing using lights 9) restriction on mechanized fishing in shallow waters. These nine regulations were used to assess disparity between regulations and fish biology for each state. The details and justifications for the percent disparity calculations are provided in detail in Supplementary Material, Section 2.

#### 4.2.6. Populating the IRIIS matrix

We first mapped all nine species based on their IRV values along the vertical axis. Species with higher IRV values were positioned higher on the vertical axis. The horizontal (management) axis was populated based on the geographic spread of each population in relation to jurisdictional boundaries. Based on the geographic extent of each population, as per rationale explained in figure 5, we classified populations into two categories - populations ranging within a single management regime and populations ranging across multiple management regimes.

Within the first category (populations ranging within a single management regime), we further classified the populations into two subcategories (from left to right): 1. populations with low disparity (disparity index less than 33%) and 2. populations with high disparity (disparity index more than 33%).

In the second category, along the management axis (population ranging across multiple management regimes), we classified populations along the horizontal axis into 3 subcategories (from left to right): 1. Low disparity in all the management regimes, 2. Low disparity in majority of the management regimes, 3. High disparity in majority of the management regimes.

## Resource availability statement

The raw demultiplexed data generated in this study will be deposited in the Sequence Read Archive (SRA) database of the online repository of NCBI (National Center for Biotechnology Information) and the accession numbers will be provided upon acceptance. The references of government regulations used for disparity analysis are provided as citations in Supplementary material, Section 2. All analytical scripts are included in the Supplementary Materials

## Acknowledgements

We thank all the fisherfolk, fish venders and retailors who allowed us to collect samples from their fish. We thank Isha Bopardikar, Alisa Barns, Al Badush Gafoor, Rajeshwari BT and Amit Nayak for help at the time of sample collection. We thank Meghana Natesh, Vinay Sagar, Aparna Krishnan, Tejali Naik, Laxminarayan CP, Vinay Kumar, Gauri Gharpure for their help in Lab work. We thank Shri Amm Murugappa Chettiar Research Centre, Cholamandalam Investment and Finance Company Limited (Chola) and Mr.Arvind Dattar for providing financial assistance for the project. We also thank Central Marine Fisheries Research Institute, Department of Fisheries – Government of India, Department of Fisheries of Gujarat, Maharashtra, Goa, Karnataka and Kerala for publicly available information on policies, schemes and subsidies and rules and regulations.

## Authors contributions

MG - Conceptualization, Data curation, Formal analysis, Funding acquisition, Investigation, Methodology, Project administration, Validation, Visualization, Writing – original draft, Writing – review & editing

RA - Conceptualization, Funding acquisition, Methodology, Project administration, Supervision, Validation, Visualization, Writing – review & editing

VS - Methodology, Supervision, Validation, Writing – review & editing

AP - Formal analysis, Methodology, Resources, Software, Validation

AT - Formal analysis, Methodology, Software, Validation

PP - Formal analysis, Software, Validation

MS - Data curation, Writing – review & editing

IM - Data curation, Writing – review & editing

JK - Methodology, Supervision, Validation, Writing – review & editing

UR - Conceptualization, Methodology, Project administration, Resources, Supervision, Validation, Writing – review & editing

## Supplemental information

Document S1. Figures S1–S13, Tables S1 to S8, and supplemental references

