## Supplementary Material for "Integrating fish biology, population genomics and fishery policies for informed management of interjurisdictional multi-species fisheries"

**Supplementary material – section 1**

**Calculating an IRV index values for ladyfish/ Silver Silago (*Sillago sihama*)**

For our analysis, we used the IRV index developed by Mohammad et al. 2021 ^1^. We requested the authors for IRV values for the target species in our analysis, which they generously provided (See Table below). However, an IRV value was not calculated for silver Silago (*Sillago sihama*) by Mohammad et al. 2021 requiring us to calculate it ourselves using the approach outlined below:

Table S1. Resilience, Vulnerability, IRV and Intrinsic rate of population growth (rm) values for eight study species

| **No.** | **Species** | **rm** | **Resilience** | **Vulnerability** | **IRV** |
| --- | --- | --- | --- | --- | --- |
| 1 | *Chirocentrus dorab* | 1 | 2 | 1.833333333 | 0.2848 |
| 2 | *Epinephelus diacanthus* | 3.2 | 1.91666667 | 2.222222222 | 0.1915 |
| 3 | *Nemipterus japonicus* | 5.2 | 2.54658385 | 2.123188406 | 0.3188 |
| 4 | *Pampus argenteus* | 3.4 | 2.39285714 | 2.333333333 | 0.2449 |
| 5 | *Rastrelliger kanagurta* | 7.64 | 2.40336135 | 2.306535948 | 0.2525 |
| 6 | *Sardinella longiceps* | 6.3 | 2.28571429 | 2.083333333 | 0.2866 |
| 7 | *Scoliodon laticaudus* | 2.84 | 1.79365079 | 1.888888889 | 0.2274 |
| 8 | *Trichiurus lepturus* | 1.26 | 1.80357143 | 2.09375 | 0.1932 |

To calculate IRV index of Silver Silago (*Sillago sihama*) we undertook the following steps

1. We identified that both the Resilience score and vulnerability score of eight out of nine study species from Mohammad et al. 2021 showed a good correlation with the intrinsic rate of population growth (rm) value reported in the online dataset FishBase ^2^

Resilience score = 0.092*rm + 1.7883, R^2^ = 0.5253

Vulnerability score = 0.1454 * In (rm) + 1.9437, R^2^ = 0.3424

2. Based on this, we used the intrinsic rate of population growth (rm) value reported in Fishbase for *Sillago sihama* (rm = 3.68) to calculate its Resilience score and vulnerability score

Resilience score = 2.126

Vulnerability score = 2.025

Based on equations in Mohammad et al. 2021 ^1^ we calculated the IRV index of *Sillago sihama* from its Resilience score and vulnerability score*.* Equation to calculate IRV index

Equation to calculate IRV index. Reproduced here from Mohammad et al. 2021^3^

$$d\_i = sqrt((3 - x\_i)^2 + (y\_i - 1)^2)$$

$$\theta\_i = tan^(-1)(y\_i / x\_i)$$

The maximum possible distance for a point in the graph is √8 and the maximum angle is π/2 radians. Hence, the authors of Mohammad et al. 2021defined IRV score ranging from 0 to 1 as

$$IRV = (1 - d\_i / sqrt(8)) (1 - \theta\_i / (\pi/ 2))$$

Here x_i_ is the species resilience value and y_i_ is the species vulnerability value

3. Based on the calculations, the **IRV value of *Sillago sihama* was 0.23**

**Supplementary material – section 2**

**Identifying the disparity between species biology and management attributes**

We identified the disparity between species biology and management attributes using following steps

1. Step 1 – identifying key management attributes and corresponding state wise regulations.

As explained in the methods section we identified nine regulations as key management attributes to assess state-wide disparity with fish biology. They were 1) partial or complete fishing ban during monsoon months (June to August) 2) Maximum allowable engine capacity of a fishing vessel 3) Minimum allowable mesh size for cod end of trawl operation 4) Minimum allowable mesh size for gill nets 5) Minimum allowable mesh size for purse nets / ring seine nets 6) Minimum allowable legal size for a species 7) Ban on Bull / Paired trawling 8) Ban on fishing using lights 9) Restriction on mechanized fishing in shallow waters. These nine regulations were used to assess state-wide disparity with fish biology. Table S2 describes the state specific regulations for these management attributes (regulations) for five states (Gujarat, Maharashtra, Goa, Karnataka and Kerala) along the west coast of India

Table S2. state specific regulations for nine chosen management regulations

|  | **Monsoon fishing ban** | **Minimum engine size** | **Mesh size - cod end trawler** | **Mesh size - gill net** | **Mesh size - purse net** | **Minimum legal size** | **Bull trawling** | **Light fishing** | **Shallow water ban** |
| --- | --- | --- | --- | --- | --- | --- | --- | --- | --- |
| **Gujarat** | 10th June to 15th August (Ports-and-Fisheries-Department-Gujarat.)^4^ | No specific regulation found | 40mm square (Ports-and-Fisheries-Department-Gujarat.) ^4^ | No specific regulation found | No specific regulation found | Shark 1-1.5m (Ports-and-Fisheries-Department-Gujarat) ^4^ | No specific regulation found | No specific regulation found | No mechanized fishing up to 5 nautical miles from shore (Ports-and-Fisheries-Department-Gujarat.)^4^ |
| **Maharashtra** | 1st June to Narali Poornima (Agriculture-and-Co-operation-Department)^5^ | Max 6 cylinder (Agriculture-and-Co-operation-Department.)^5^ | 35mm, 25mm – Ratnagiri (Government-of-Maharashtra.)^6^ | No specific regulation found | 25mm - sardine, 49mm – mackerel (MATSYAVI-1115/CR-282/ADF-14.)^7^ | No specific regulation found | Banned (Law-and-Judiciary-Department-Maharashtra.)^8^ - | No Night fishing (Agriculture-and-Co-operation-Department.)^5^ | No mechanized vessels in less than 5 fathoms depth up to Burdondi, and in less than 10 fathoms south of Burdondi (Agriculture-and-Co-operation-Department.)^5^ |
| **Goa** | 1st June to 31st August (Shamila Monteiro, Samir R Korgaonkar, Chandrakant D Velip, Smita A Mazumdar, Megha S Kerkar.) ^9^ | No specific regulation found | 24mm (Order 2–1-81FSH (V).)^10^ | No specific regulation found | No specific regulation found | Indian oil sardine - 10 TL  Indian Mackerel - 14TL  Ribbonfish - 46 TL  Silver silago - 11.3 TL  Japanese threadfin bread - 12 TL  Silver pomfret - 13TL,  Spinycheek grouper - 18TL (Department-of-Fisheries-Goa.)^11^ | Banned (No. DF/ENF/ORDER/2016-17.)^12^ | Banned (No. DF/ENF/ORDER/2016-17.)^12^ | No trawl and purse net allowed within 5 km from the shore (DF/ENF/RULE AMENDMENT/2021-22.)^13^ |
| **Karnataka** | 1st June to 31st July (Government-of-Karnataka.)^14^ | Regulated (Department-of-Fisheries-Karnataka.)^15^ | 35 mm (Government-of-Karnataka.)^16^ | No specific regulation found | 20mm (Notification on mesh size regulation for Purseine net.)^17^ | Indian oil sardine - 10 TL  Indian Mackerel - 14TL  Ribbonfish - 46 TL  Silver silago - 11.3 TL  Japanese threadfin bread - 12 TL  Silver pomfret - 13TL,  Spinycheek grouper – 18TL (Department-of-Fisheries-karnataka.)^18^ | Banned (Department-of-Fisheries-karnataka.)^19^ | Banned(Department-of-Animal-Husbandry-Dairying-&-Fisheries.)^20^ | No mechanized fishing up to 5 km from the shore  (Karnataka_Marine_fishing_regulation_Act1986.)^21^ |
| **Kerala** | June 9th to July 31^st^ (Fisheries.and.Ports.(B).Department. )^22^ | Specified (Fisheries-and-ports(B)-Department-Kerala.)^23^ | 35sq, 25sq (Fisheries-and-ports(B)-Department-Kerala.)^23^ | Various rules (Fisheries-and-ports(B)-Department-Keral.)^23^ | 10dimond, 22 diamond (Fisheries-and-ports(B)-Department-Kerala.)^23^ | TL  Indian Mackerel - 14TL  Indian oil sardine - 10 TL  Silver pomfret -13TL  Spiny cheek grouper - 18TL  Japanese threadfin bream - 10TL(K S Mohamed et al.)^24^ | No specific regulation found | Max12 v is allowed (Fisheries-and-ports(B)-Department-Kerala.)^23^ | No mechanized fishing in waters less than 20 m in depth (Karnataka_Marine_fishing_regulation_Act1986.)^21^ |

2. Step 2 – Developing criteria to identify whether regulations are congruent with species biology.

We scored each regulation - state combination between zero and one, depending on if the state-specific regulation was ‘congruent’ with species biology (0) or not (1). We assessed ‘congruence’ based on whether the regulation in question aligned with sustainable harvest requirements for that species, based on what is currently known of species’ biology). A lack of congruence (or disparity) reflects either that the regulation does not align with sustainable harvest requirements, or the absence of regulation in that state. If for particular species, a regulation is not relevant, we excluded it from the disparity analysis. For eg. silver sillago / ladyfish are not found in the area where purse nets operate and thus, they are not caught by purse nets. Thus, purse net mesh size regulation is irrelevant for silver sillago and we remove it from the disparity score calculation

Table S3. Criteria used to assess congruence or disparity with the species biology of each regulation

| **Regulations** | **Biological congruence / disparity** | **Explanation** | **Disparity score** |
| --- | --- | --- | --- |
| Monsoon ban | congruence | Known breeding season of species in question overlaps with the monsoon ban | 0 |
|  | disparity | - Regulation absents in the state in question **OR** - Known breeding season species in question does not overlap with the monsoon ban | 1 |
| Maximum legal engine capacity | congruence | State in question has specified a maximum engine capacity for mechanized and motorized boats | 0 |
|  | disparity | Regulation absent in the state in question | 1 |
| Minimum mesh size for cod end of trawl nets | congruence | - The l50^[1]^ value of specified mesh size is at least 85% of Lm^[2]^  **OR** - Regulation present, but there exists no study identifying l50 value for given species | 0 |
|  | disparity | - Regulation absent in the state in question **OR** - The l50 value of specified mesh size is less than 85% of Lm | 1 |
| Minimum mesh size for gill nets | congruence | - The l50 value of specified mesh size is at least 85% of Lm  **OR** - Regulation present, but there exists no study identifying l50 value for given species | 0 |
|  | disparity | - Regulation absent in the state in question **OR** - The l50 value of specified mesh size is less than 85% of Lm | 1 |
| Minimum mesh size for purse nets | congruence | - The l50 value of specified mesh size is at least 85% of Lm  **OR** - Regulation present, but there exists no study identifying l50 value for given species | 0 |
|  | disparity | - Regulation absent in the state in question **OR** - The l50 value of specified mesh size is less than 85% of Lm | 1 |
| Minimum legal size of a species | congruence | Minimum allowable legal size specified for species in question | 0 |
|  | disparity | - Regulation absent in the state in question **OR** - Minimum allowable legal size is not identified for the species in question in the regulation | 1 |
| Ban on bull trawling | congruence | Regulation present in the state in question | 0 |
|  | disparity | Regulation absent in the state in question | 1 |
| Light fishing ban | congruence | Regulation present in the state in question | 0 |
|  | disparity | Regulation absent in the state in question | 1 |
| Shallow water fishing restriction | congruence | Regulation present in the state in question | 0 |
|  | disparity | Regulation absent in the state in question | 1 |

3. Step 3 – Identifying disparity between species biology and individual management regulations

In theory, the congruence and disparity for each regulation could change based on the biology of individual species. However, the only species-specific information pertinent to these regulations available for species along the Indian west coast of India are a) information on breeding seasons and b) size frequency spectrum information of fish caught in the cod end of trawl net. We were therefore able to assess a) how closely the partial or complete fishing ban during monsoon months (June to August) overlapped with known breeding seasons for each species and b) if the I50 value of minimum allowable mesh size for the cod end of trawl operations was at least 85% of Lm for each assessed species (see below). For all other management regulations, given the lack of data, we assumed congruence with species biology if the regulation was present in given the state, as per the table S3.

a) Partial or complete fishing ban during monsoon months (June to August).

Based on available information complied in figure S1 we can observe that the peak spawning periods of Japanese threadfin bream (*Nemipterus japonicus*), Spiny cheek grouper (*Epinephelus diacanthus*), Spade nose shark (*Scoliodon laticaudus)* and wolf herring (*Chirocentrus nudus*), do not overlap with ban period for all the states along west coast of India^25^. Thus, the populations of these four species show disparity with the monsoon fishing ban regulation.

Figure S1. Spawning periods of nine study species along the west coast of India

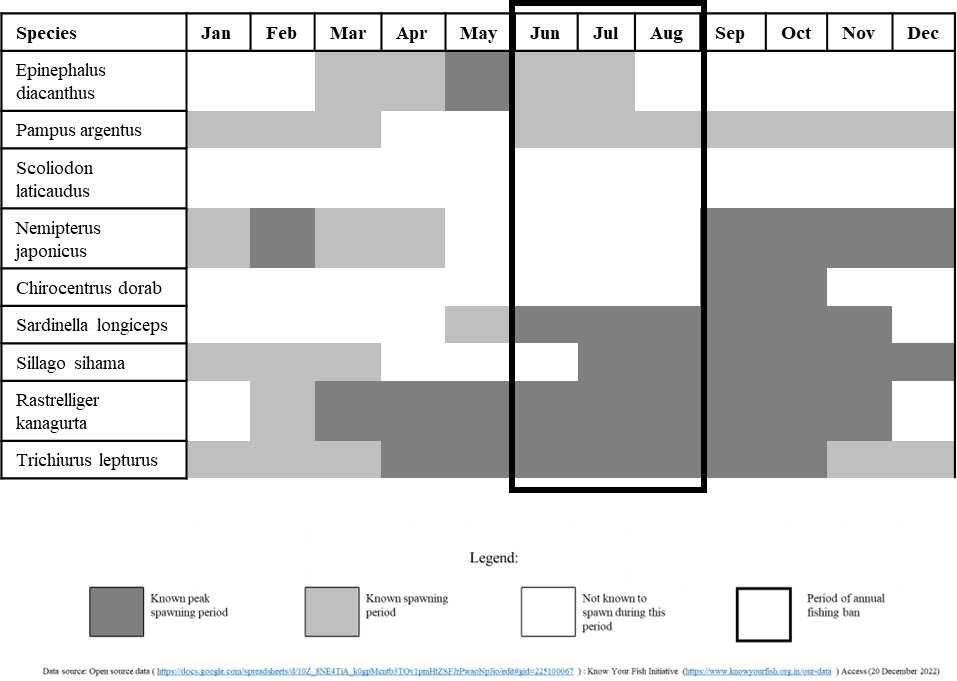

b) Minimum allowable mesh size for cod end of trawl operation

For the trawl cod end mesh size regulation, we assessed the biological congruence or disparity based on L50/Lm ratios. L50 is the length at 50% retention (L50) for a species in a particular fishing gear and Lm is the length of maturity for that species / population. Higher L50/Lm values signify a higher proportion of mature individuals caught in that fishing gear. Since mesh size regulations are designed to keep juveniles out of nets, we consider higher L50/Lm ratios for any species – gear combination to be congruent with species biology. For this analysis we assume L50/Lm ≤ 0.85 (85%) to have low biological congruence (high disparity between biology and regulation).

We first collated information on known L50 / Lm ratio from published data in the study area (table S4). We then compared this ratio with trawl cod end mesh size regulations of individual states to identify congruence or disparity between regulation and species biology (table S5).

Table S4. L50 / Lm ratios for nine study species from published data in the study area

| **Species** | **Length at maturity**  **(FishBase)** | | **Length of maturity (Lm) based on most recent study from the region** | | **Size of fish at L50** | | **l50/Lm** |
| --- | --- | --- | --- | --- | --- | --- | --- |
|  | **Mean** | **Range** | **Lm Mean** | **Study area (most recent study from the region)** | **Net attributes** | **l50** |  |
| spiny cheek grouper (*Epinephelus diacanthus*) | 30.9 | 23.1-41.4 | 20 | Southwest coast of India (Ramachandran & Ramalingam.)^26^ | 60 mm upper and 20 mm lower(Boopendranath, Pravin, Gibinkumar, Sabu, & Madhu.) ^27^ | 10.59 | 0.52 |
| silver pomfret (*Pampus argenteus*) | 30.9 | 23.1-41.4 | 27.5 | Gujarat (Ghosh et al.)^28^ | 30 mm (Paighambari & Eighani.)^29^ | 11 | 0.4 |
|  |  |  | 23 | Maharashtra (Gladston, Devi, & Xavier.)^30^ | 30 mm (Paighambari & Eighani.)^29^ | 11 | 0.47 |
| Japanese threadfin bream (*Nemipterus japonicus*) | 15.8 | 11.8-21.2 | 24 | Saurashtra Gujarat (Sarman, Hitesh, Vinaykumar, Mahendra, & Piyush.)^31^ | 40 mm (Bhairol, Dora, & Talwar.)^32^ | 12 (diamond) 16 (square) | 0.5 diamond, 0.66 square |
| Indian oil sardine (*Sardinella longiceps*) | 12.9 | 9.6-17.3 | 15.5 | Gujarat (Motivarash, Kardani, Katariya, & Bhatt.)^33^ | 60 mm upper and 20 mm lower (Boopendranath et al.)^27^ | 12.69 | 0.81 |
|  |  |  | 15.5 | Southern population (Nair, Joseph, Kripa, Remya, & Pillai.)^34^ | 60 mm upper and 20 mm lower (Boopendranath et al.)^27^ | 12.69 | 0.81 |
| Silver silago /ladyfish (*Sillago sihama*) | 15.6 | 11.7-20.9 | 16.9 | Karnataka (Vinod & Basavaraja.)^35^ | 40 mm Bull trawl (Ratanawalee Phoonsawat, Udomsin Auksonphaob, Thitipon Cheuamankong, & Sampan Panjarat.)^36^ | 19.2 | 1.13 |
| Indian mackerel (*Rastrelliger kanagurta*) | 11.5 | 11.4-20.5 | 19.8-20.6 | Maharashtra (Bhendarkar, Naik, Mohite, & Kulkarni.)^37^ | 40 mm Bull trawl (Ratanawalee Phoonsawat, Udomsin Auksonphaob et al.)^36^ |  | 0.63-0.65 |
| Ribbonfish (*Trichiurus lepturus*) | 72.9 | 54.4-97.7 | 55.4 | Karnataka (Rajesh, Rohit, Vase, Sampathkumar, & Sahib.)^38^ | 40 mm (Vettiyattil, Herrmann, & Bharathiamma.)^39^ | 46.95 (diamond) | 0.84 |
|  |  |  |  |  | 40 mm (Vettiyattil et al.)^39^ | 56.75 (square) | 1.02 |
| wolf herring (*Chirocentrus nudus*) | 46.2 | 34-61 | 41-43 (*Chirocentrus dorab*) | Tamil Nadu Abdussamad, Pillai, Zacharia, & Jeyabalan.)^40^ | No available information |  |  |
|  |  |  | 48-49 (*Chirocentrus nudus*) | Tamil Nadu (Abdussamad et al.)^40^ |  |  |  |
| Spade nose shark (*Scoliodon laticaudus*) | 36.2 | 27-48 | 35.79 | Gujarat (Sen et al.)^41^ | No available information (Boopendranath et al.)^27^ | all are caught |  |
|  |  |  | 36 | Maharashtra (Barnes, Sutaria, Harry, & Jabado.)^42^ |  |  |  |

Table S5. congruence or disparity between trawl cod end mesh size regulation and biology of study species.

| **Species** | **L50/Lm** | **State** | **Minimum allowable mesh size** | **Judgement** | **Rationale** |
| --- | --- | --- | --- | --- | --- |
| spiny cheek grouper (*Epinephelus diacanthus*) | 0.5295 | Gujarat | 40 | Congruence | L50/Lm ratio was calculated for 20mm mesh size for the lower cod end. We do not have information on L50 values for higher mesh sizes. We thus give the benefit of the doubt to states other than Goa, which have higher minimum allowable mesh size. |
|  |  | Maharashtra | 35 | Congruence |  |
|  |  | Goa | 24 | Disparity |  |
|  |  | Karnataka | 35 | Congruence |  |
|  |  | Kerala | 35 | Congruence |  |
| Silver pomfret (*Pampus argenteus*) | 0.4, 0.47 | Gujarat | 40 | Congruence | L50/Lm ratio was calculated for 30 mm mesh size for lower cod end. We do not have information on L50 values for higher mesh sizes. We thus give the benefit of the doubt to states other than Goa, which have higher minimum allowable mesh size. |
|  |  | Maharashtra | 35 | Congruence |  |
|  |  | Goa | 24 | Disparity |  |
|  |  | Karnataka | 35 | Congruence |  |
|  |  | Kerala | 35 | Congruence |  |
| Japanese threadfin bream (*Nemipterus japonicus*) | 0.5 for Dimond mesh, 0.66 for square mesh | Gujarat | 40 | Disparity | Gujarat has made 40 mm mesh sizes mandatory, but square meshes are not mandatory, and diamond meshes have poor results |
|  |  | Maharashtra | 35 | Congruence | In Maharashtra, square mesh sizes beyond 40mm are mandatory |
|  |  | Goa | 24 | Disparity | Mesh sizes less than 40 mm, and no shape regulation |
|  |  | Karnataka | 35 | Disparity | Mesh sizes less than 40 mm, and no shape regulation |
|  |  | Kerala | 35 | Congruence | Square meshes are mandatory, but minimum cod end mesh size is 35 mm. However, due to lack of data, we give the benefit of the doubt to Kerala |
| Indian oil sardine (*Sardinella longiceps*) | 0.818 | Gujarat | 40 | Congruence | Even though l50 is 20% lower than Lm, L50 is calculated with a smaller mesh size (20 mm lower cod end) than legal mesh sizes in all the states except Goa. We therefore give the benefit of the doubt to other states. In contrast, the legal mesh size requirement in Goa is very low (maximum 24), which is why we score it as a disparity |
|  |  | Maharashtra | 35 | Congruence |  |
|  |  | Goa | 24 | Disparity |  |
|  |  | Karnataka | 35 | Congruence |  |
|  |  | Kerala | 35 | Congruence |  |
| Silver silago /ladyfish (*Sillago sihama*) | 1.13 | Gujarat | 40 | Congruence | There is a good biological congruence between l50 and Lm in 40 mm cod end net fished by bull trawling. We assume that otter trawling operations even at lower cod end mesh sizes will show better L50/Lm ratios. As a result, even though Goa only has a 24 mm minimum mesh size requirement, we will give it the benefit of the doubt |
|  |  | Maharashtra | 35 | Congruence |  |
|  |  | Goa | 24 | Congruence |  |
|  |  | Karnataka | 35 | Congruence |  |
|  |  | Kerala | 35 | Congruence |  |
| Indian mackerel (*Rastrelliger kanagurta*)) | 0.63-0.65 | Gujarat | 40 | Congruence | Since the known L50 findings are from bull/paired trawling, we expect poor L50/Lm ratios to improve from bull trawling to otter trawling. However, because of low legal mesh size we cannot extend this to Goa |
|  |  | Maharashtra | 35 | Congruence |  |
|  |  | Goa | 24 | Disparity |  |
|  |  | Karnataka | 35 | Congruence |  |
|  |  | Kerala | 35 | Congruence |  |
| Ribbonfish (*Trichiurus lepturus*) | 0.84-1.02 | Gujarat | 40 | Congruence | Goa has very low mesh size restrictions and even Karnataka has no restriction on types of net (diamond or square mesh) on a lower mesh size than 40 mm. Thus, we expect L50/Lm ratios to decline in those cases. Maharashtra allows 40 mm mesh sizes and square nets. Kerala allows 35 mm mesh sizes but square nets. Although Gujarat has no restrictions on nets, it allows 40 mm mesh sizes, which varies only 15% from the Lm. We therefore gave Gujarat the benefit of the doubt and consider Gujarat mesh size regulation to be in congruence with ribbonfish biology along with Maharashtra and Kerala |
|  |  | Maharashtra | 35 | Congruence |  |
|  |  | Goa | 24 | Disparity |  |
|  |  | Karnataka | 35 | Disparity |  |
|  |  | Kerala | 35 | Congruence |  |
| wolf herring (*Chirocentrus nudus*) | No information available | Gujarat | 40 | Congruence | Since there is no available information on L50 in trawl nets of any mesh size we gave the benefit of the doubt to these species |
|  |  | Maharashtra | 35 | Congruence |  |
|  |  | Goa | 24 | congruence |  |
|  |  | Karnataka | 35 | Disparity |  |
|  |  | Kerala | 35 | Congruence |  |
| Spade nose shark (*Scoliodon laticaudus*) | No information available | Gujarat | 40 | Congruence | Since there is no available information on L50 in trawl nets of any mesh size we gave the benefit of the doubt to these species |
|  |  | Maharashtra | 35 | Congruence |  |
|  |  | Goa | 24 | congruence |  |
|  |  | Karnataka | 35 | Disparity |  |
|  |  | Kerala | 35 | Congruence |  |

4. Step 4 – constructing a disparity matrix: We constructed a disparity matrix for every genetically distinct population of our target species using the protocols described above. See the main text for more details on our approach. In figure S2 we have catalogued disparity matrix for each genetically distinct population of target species

Figure S2. Disparity matrix for each genetically distinct population of target species

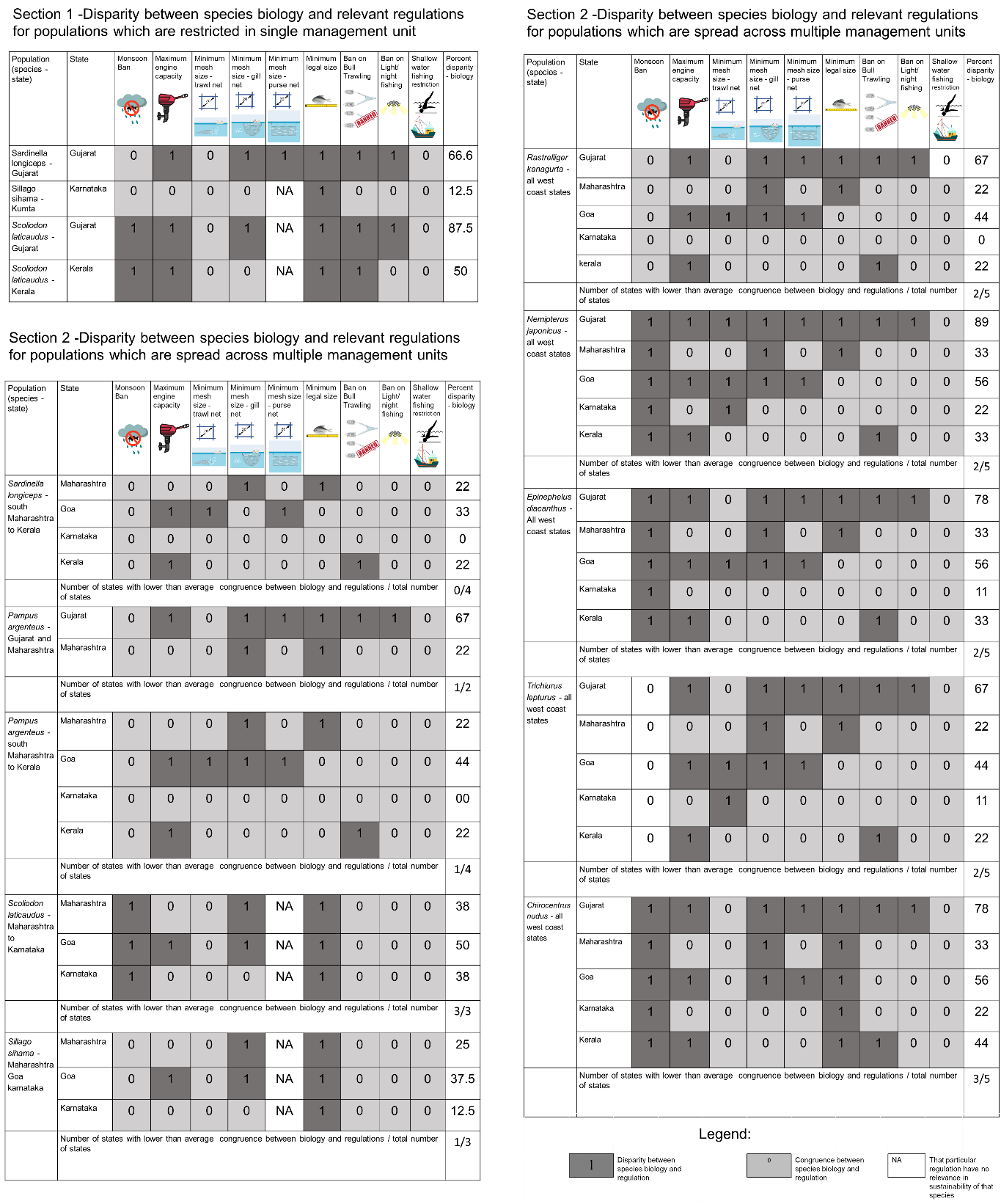

**Supplementary material – section 3**

**Using Single Nucleotide Polymorphisms (SNPs) to identify the geographical extent of individual fish populations along the Indian west coast.**

1. **Sample collection**

We sampled 9 species of fish commonly caught in commercial fishing operations along the Indian west coast. The selected fish species were silver pomfret (*Pampus argenteus*), ribbonfish (*Trichiurus lepturus*), Japanese threadfin bream (*Nemipterus japonicus*), spiny cheek grouper (*Epinephelus diacanthus*), spade nose shark (*Scoliodon laticaudus)*, wolf herring (*Chirocentrus nudus*), Indian oil sardine (*Sardinella longiceps),* ladyfish (*Sillago sihama*), and Indian mackerel (*Rastrelliger kanagurta*),

Table S6. Target species with their habitat and comparative commercial values. Habitat and Food sourced from Fishbase profiles for target species ^63^.

|  | **Species name** | **Habitat** | **Food** |
| --- | --- | --- | --- |
| 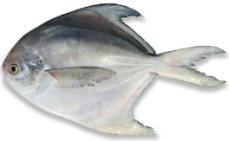 | Silver pomfret (*Pampus argenteus*) | Muddy Bottoms | Zooplankton |
| 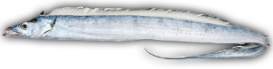 | Ribbonfish (*Trichiurus lepturus*) | Muddy Bottoms | Fish, squid etc |
| 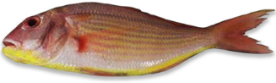 | Japanese threadfin bream  (*Nemipterus japonicus*) | Mud and sand bottoms | Small fish, crustaceans etc. |
| 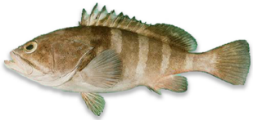 | Spiny cheek grouper (*Epinephelus diacanthus*) | Muddy substrate | Fish |
| 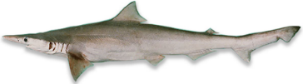 | Spade nose shark (*Scoliodon laticaudus)*) | Rocky substrate | Small fish, shrimp |
| 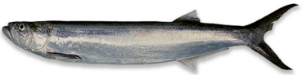 | Wolf herring (*Chirocentrus nudus*) | Inshore waters | Small fish, crustaceans etc |
| 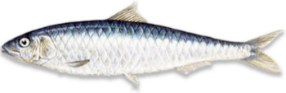 | Indian oil sardine (*Sardinella longiceps)* | Coastal pelagic | Phytoplankton, crustaceans etc. |
| 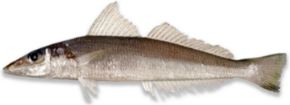 | Ladyfish/ Silver Silago  (*Sillago sihama*) | Beach, sandbank, estuaries | Polychaete, shrimps |
| 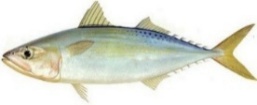 | Indian mackerel (*Rastrelliger kanagurta* | Coastal pelagic | Small zooplankton |

We sampled individuals of each species from landing sites located at roughly 200 km intervals along the entire west coast from Gujarat to Kerala (See figure S3)

Figure S3. sampling locations along the west coast of India

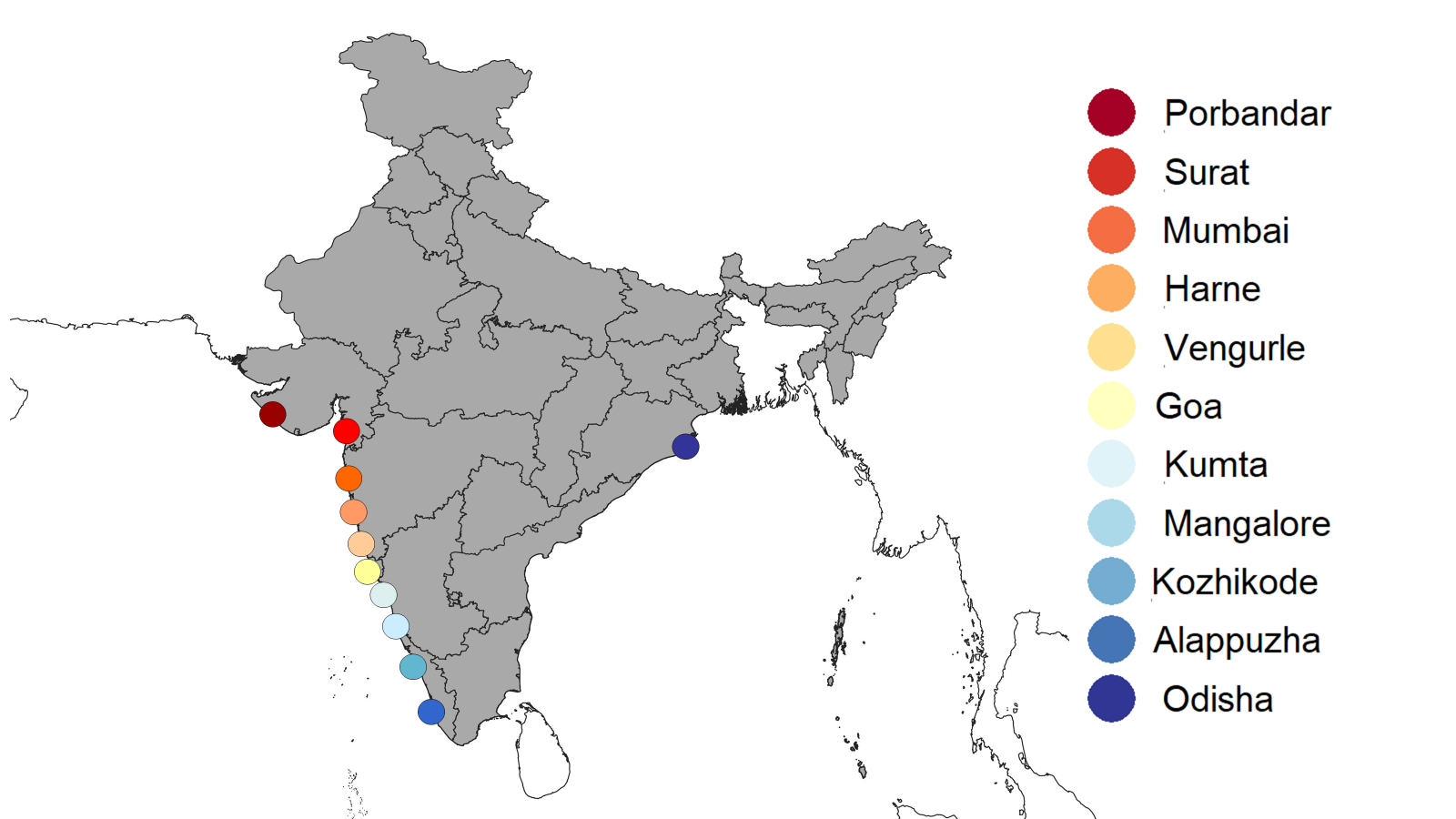

Tissue samples were collected using fin clips extracted from the fish directly at the landing site. Fin clips of 10-18 individuals from each species were collected at each sampling location using sterilized instruments to avoid cross-contamination, and stored in absolute ethanol in a 2 ml tube. We ensured that the fish sampled were sourced from local waters since many fish breed and spawn at different locations. Sampling fish without considering their life stages posed the risk of mixing individuals from two distinct populations, leading to potential misidentification. To avoid this, we collected samples from only the adult population at each location.

2. **Genetic Analysis**

DNA extraction was performed for all the collected samples using QIAamp DNA Blood Mini Kit (Qiagen, catalogue no 69504) following protocols provided by the manufacturer. Extractions were done in batches of 11 samples each, along with one extraction control to check for contamination.

We used a genome-wide approach (Single Nucleotide Polymorphism (SNPs) to identify population structure and genetic differentiation for all species across multiple locations. We used double-digest restriction-site associated DNA (ddRAD) sequencing to generate the data and used a de-novo assembly approach for analysis as there was no genomic information available for most of the species studied^43^.

**2.1.** **Library preparation and ddRAD sequencing**

ddRAD sequencing involves digesting the input DNA with two different enzymes to compare fragments across different individuals. To maximize the number of common SNPs and to understand the population structure of nine fish species, choosing an enzyme combination that yields sufficient fragments of 300-500 bp was crucial. To determine the optimal combination, we digested the extracted DNA of three taxonomically diverse species: Spiny cheeked grouper (*Epinephelus diacanthus*), Largehead hairtail (*Trichiurus lepturus*), and Spade nose shark (*Scoliodon laticaudus*), to maximise taxonomic diversity. We used multiple enzyme combinations (EcoRI-MspI, EcoRI-MlucI, SpHI-MucI), and bioanalyzer (Agilent) profiles to finalize the enzyme combination for ddRAD library preparation. Based on the results, we identified a combination of SpH1 and MluC1 as the ideal choice (a combination of four and six base cutters), providing the maximum number of sequential fragments for all target species.

After finalizing the enzyme combination, we prepared ddRAD libraries following the protocol developed and standardized by Tyagi et al., 2024^44^. Digested products were subjected to adapter ligation, purification, and indexing. To reduce PCR bias, indexing PCR was done in three replicates, and we pooled all replicates before dual-size selection using AMPure XP magnetic beads^45^. These libraries were sequenced on the Illumina HiSeq2500/Novaseq platform using the PE 2X100 sequencing kit.

**2.2. SNP calling and filtering**

The demultiplex fastq reads were then checked for quality using FastQC^46^. For Silver Pomfret (*Pampus argenteus*) and Indian Mackerel (*Rastrelliger kanagurta,*) we used available reference genomes from NCBI database for SNP calling (*Rastrelliger kanagurta -* GCA_024605255.1, *Pampus argenteus* - GCA_000697985.1). For Spiny cheek grouper (*Epinephelus diacanthus*) and Spade nose shark (*Scoliodon laticaudus)* we used reference genomes of taxonomically nearest related species. We used the *Epinephelus coioides* (GCA_900536245.1) genome for *Epinephelus diacanthus* and *Scyliorhinus canicular* (GCA_902713615.1) genome for *Scoliodon laticaudus* from the NCBI database. For the rest of the species due to lack of available reference genome we used the *de novo* variant calling pipeline by STACKS^47^

**2.2 a. SNP calling for species with reference genomes.**

The reads were trimmed using trimmomatic and mapped to respective reference genomes using the **BWA mem algorithm**^48^. Post mapping variant calling was done using **freebayes**^49^. Vcftools was used for filtration^50^. Scripts of all the codes used to process the demultiplex fastq reads to VCF files are given in section 4

**2.2.b. SNP calling for species without reference genome**.

Preliminary filtering was done using the function **process_radtags** in the program STACKS^47^. The parameters used for this filtering were based on Paris et al. 2017^51^ (M = 5, m = 3, n = 5). We used the **denovo.map_pl** algorithm in STACKS for creating bam files. Variant calling was done later with the help of function **populations**. After variant calling, vcftools was used for filtration^50^. Scripts of all the codes used to process the demultiplex fastq reads to VCF files are given in section 4

**2.3. Filtration**

We use the sequence written in table S7 for filtration in a given sequence. After steps 5,6 and 7 we assessed individuals for proportion of missing data. Individuals with a high proportion of missing data were removed.

Table S7. Filtration sequence of variants to identify Single Nucleotide Polymorphisms (SNPs)

| Sequence | Command | Meaning | Value |
| --- | --- | --- | --- |
| 1 | –remove-indels | Removing any variant that alters the length of the REF allele |  |
| 2 | –minQ (for sp with reference genome) | Minimum mapping quality required | 30 |
| 3 | –minGQ | Minimum quality required | 30 |
| 4 | –minDP | Minimum depth required | 10 |
| 5 | –min-alleles <integer>  –max-alleles <Integer> | Include only those sites which have number of alleles greater than or less than 2 | 2 |
| 6 | –hwe | Assess for Hardy Weinberg equilibrium and remove sites which show deviation from it | P value 005 |
| 7 | –mac | Include sites with minor allele count greater than or equal to | 3 |
| 8 | –max-missing | Exclude sites on the basis of proportion of missing data | *Rastrelliger kanagurta* - 1  *Chirocentrus nudus* - 0.9  *Epinephelus diacanthus* - 1  *Trichiurus lepturus* - 0.9  *Nemipterus japonicus* - 0.9  *Pampus argenteus* - 1  *Sardinella longiceps* - 0.75  *Scoliodon laticaudus -* 0.75  *Sillago sihama* - 0.9 |

**2.4. Population genetic analysis**

The genetic population structure was determined using three approaches.

1. The visualization is based on a principal component analysis (PCA) using PLINK^52^.

2. The visualization based on admixture probabilities using program ADMIXTURE^53^. We ran Admixture for K ranging from 1 to n, where n = the number of sites in which we have got information of that particular species, for 10 replicates each, the best K was determined by estimating the cross-validation error. The results were analyzed and plotted using a web-based server, CLUMPAK^54^

3. A pairwise mean FST calculation done using STACKS^47^

**Results –**

We sequenced 311 samples belonging to nine study species across nine locations. After removing samples with low quality during the filtering process, we used a total of 273 samples for the final analysis. On average, we analyzed 4-5 samples per species per location

Figure S4. Sample sizes used for the SNP analysis for each species across all study sites

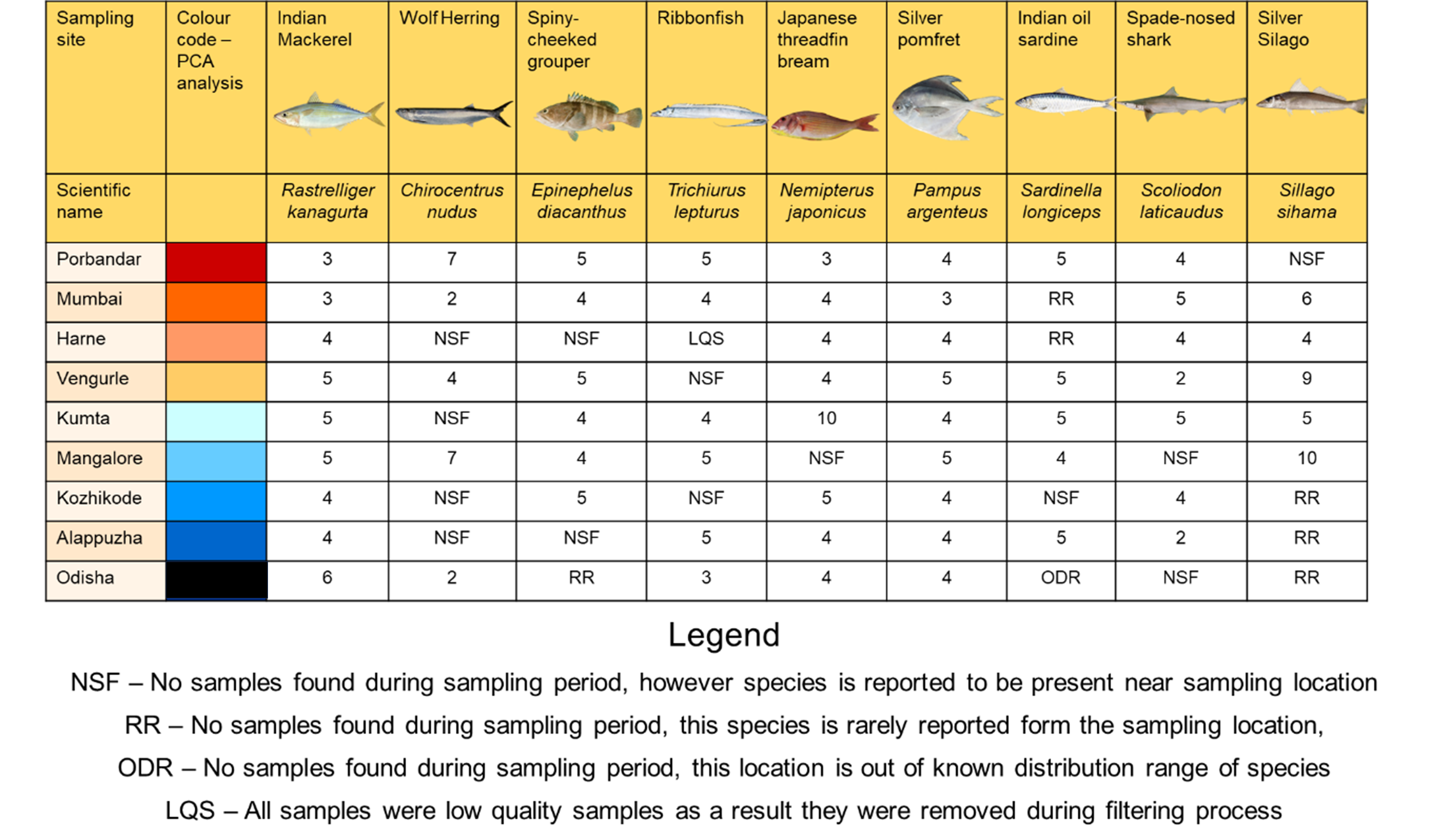

We employed three analytical methods (PCA, Admixture and FST) to comprehensively investigate the genetic structure of all species. In next few pages we have presented results of all the three analytical approaches mentioned above for each of the nine species followed by an explanatory paragraph

**1. Indian Mackerel (*Rastrelliger kanagurta*)**

Our findings indicate no signature of population structure or differentiation within the Indian mackerel samples on the west coast of India. PCA analysis demonstrates that the mackerel exhibit a panmictic population (Fig. S5B). These conclusions were further supported by Admixture analysis, where the best-supported model was found to be K=1 (Fig. S5D), along with pair-wise Fst values showing no differentiation (Fig. S5F). Despite including samples from the east coast of India in our analysis to examine any potential east-west structure, we found that the Indian mackerel comprises a single panmictic population, with no observed differentiation (Fig. S5.A, S5.C, and S5E).

**Figure S5. Genetic population structure results of Indian Mackerel**

| 1. *Rastrelliger kanagurta (*Indian Mackerel)  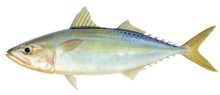 | | | | | | |
| --- | --- | --- | --- | --- | --- | --- |
| **Sampling locations:** Map of sample points and Legend of PCA plots  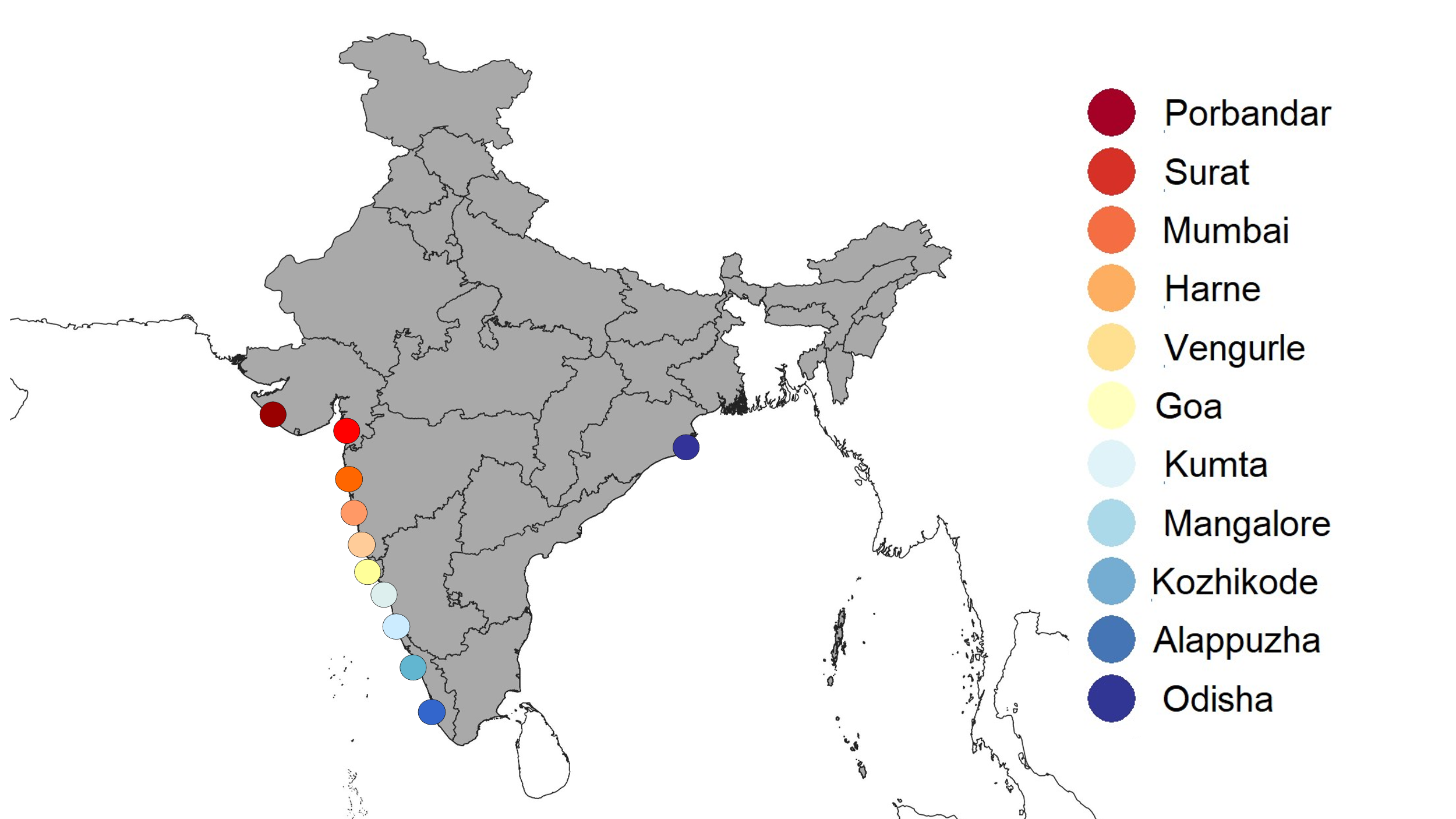 | | **A:** PCA plot - West coast Indian mackerel samples with Odisha  No Of SNPs – 13702  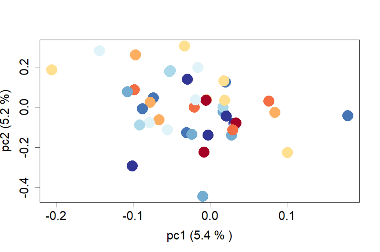 | | | **B:** PCA plot - West coast Indian mackerel samples  Number of SNPs – 7780  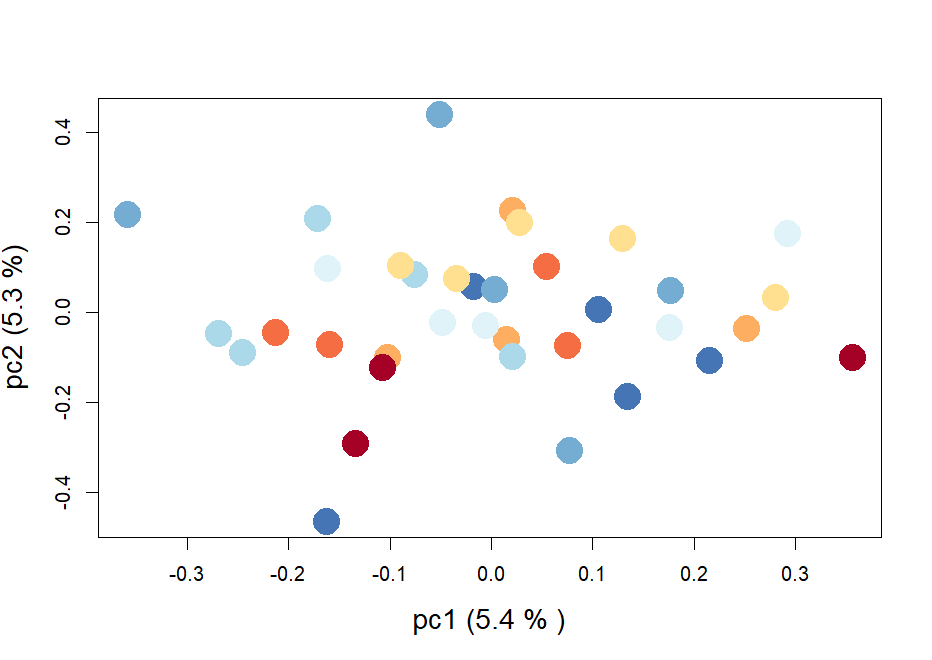 | |
| **Admixture plots (Best K is identified by black border)** | | | | | | |
| **C:** Admixture plots – west coast Indian mackerel samples with Odisha (east coast) | | | CV errors | **D:** Admixture plots – west coast Indian mackerel samples | | CV errors |
| 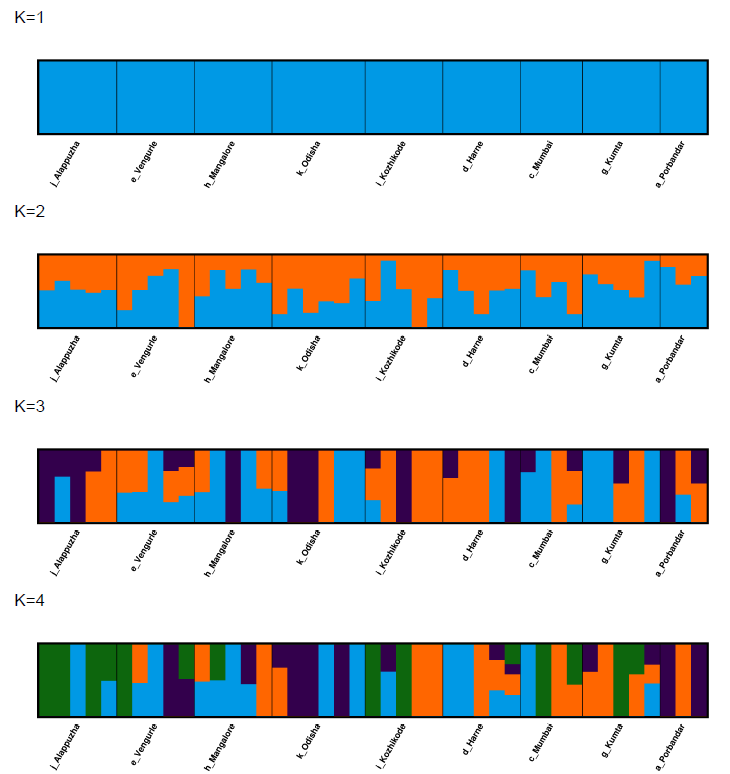 | | | 0.4323  0.5285  0.6095  0.6928 | 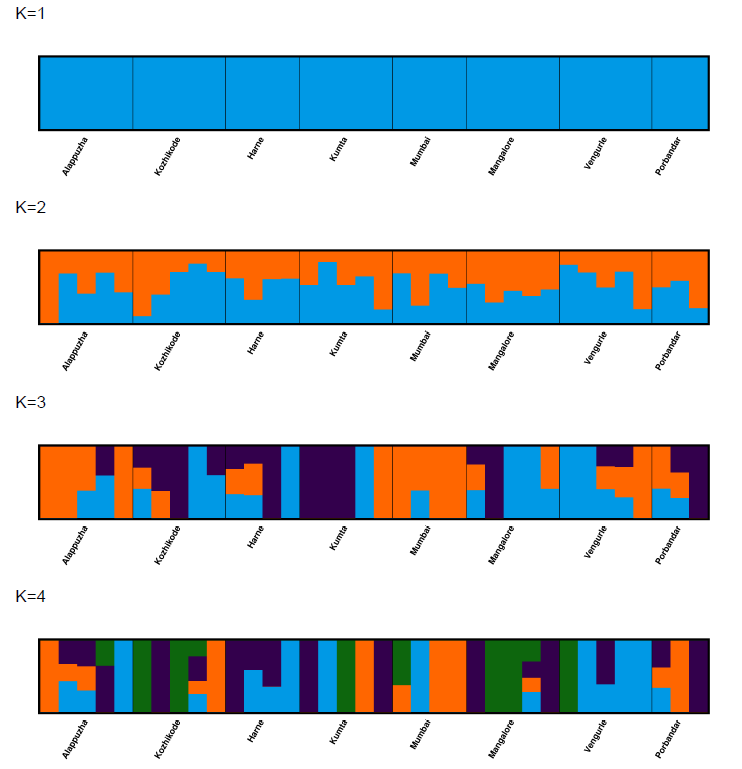 | | 0.4752  0.5876  0.6836  0.7960 |
| **Pairwise fixation index (FST) matrix for samples from each site** | | | | | | |
| **E:**  **F:** | 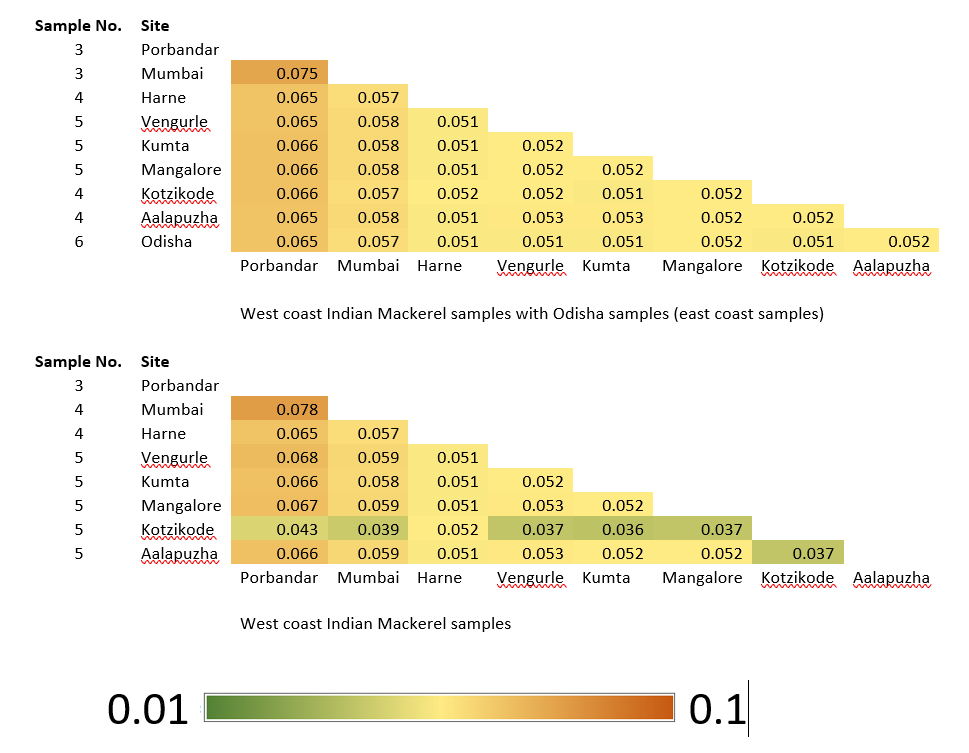 | | | | | |

**2. Wolf herring (*Chirocentrus nudus*)**

Our findings indicate no signature of population structure or differentiation within the wolf herring samples on the west coast of India. PCA analysis demonstrates that the mackerel exhibit a panmictic population (Fig S6B). These conclusions were further supported by Admixture analysis, where the best-supported model was found to be K=1 (Fig S6D), along with pair-wise Fst values showing no differentiation (Fig. S6F). Despite including samples from the east coast of India in our analysis to examine any potential east-west structure, we found that the wolf herring comprises a single panmictic population, with no observed differentiation (Fig. S6A, S6C, and S6E).

**Figure S6. Genetic population structure results of wolf herring**

| 2. *Chirocentrus nudus (*Wolf herring)  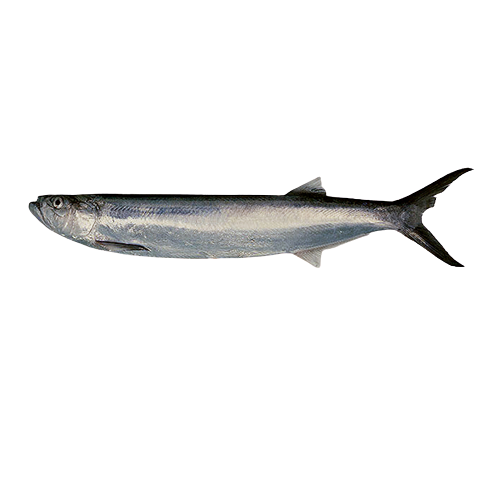 | | | | | | |
| --- | --- | --- | --- | --- | --- | --- |
| **Sampling locations:** Map of sample points and Legend of PCA plots  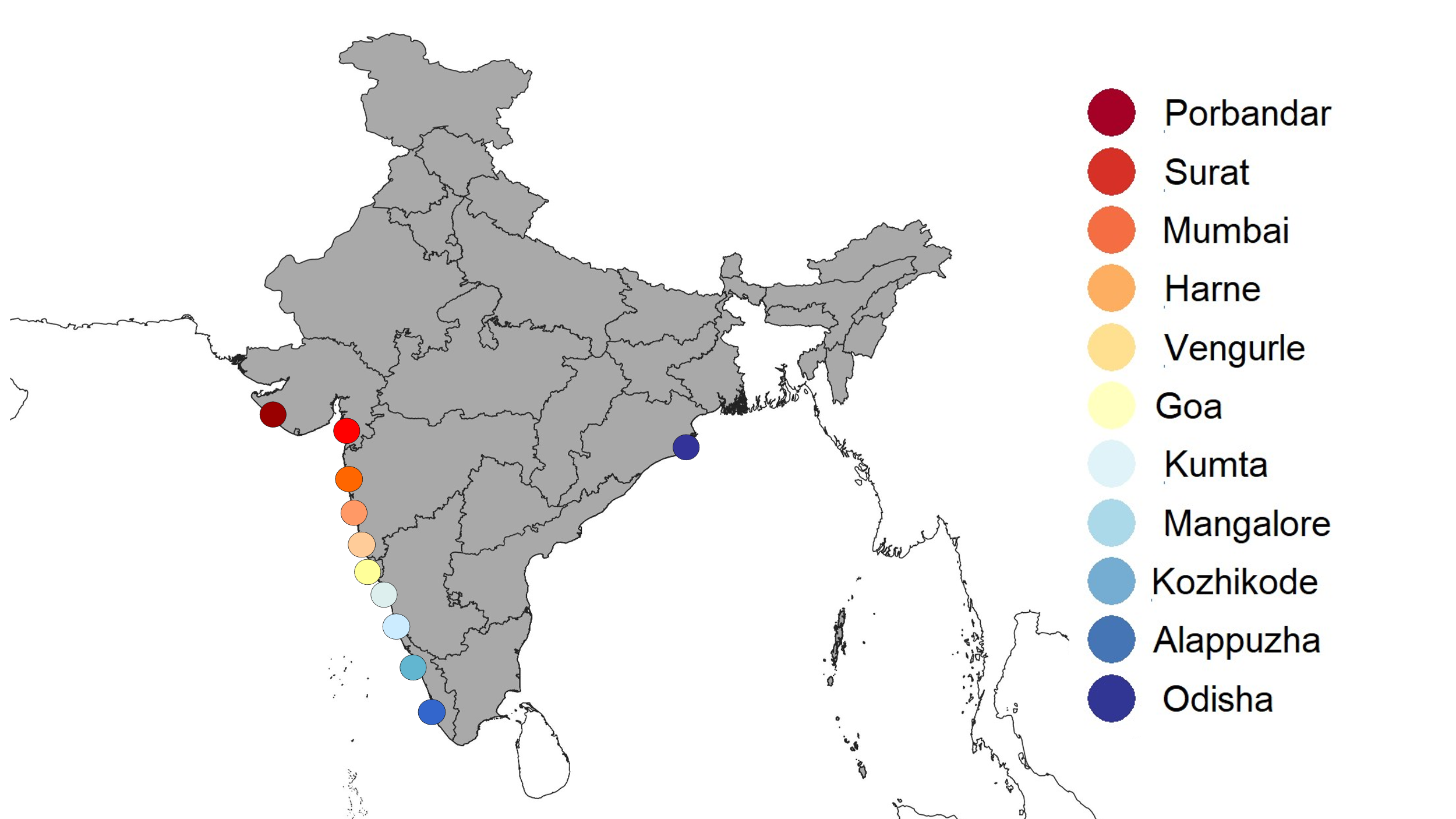 | | **A:** PCA plot - West coast Wolf herring samples with Odisha  No Of SNPs – 6747  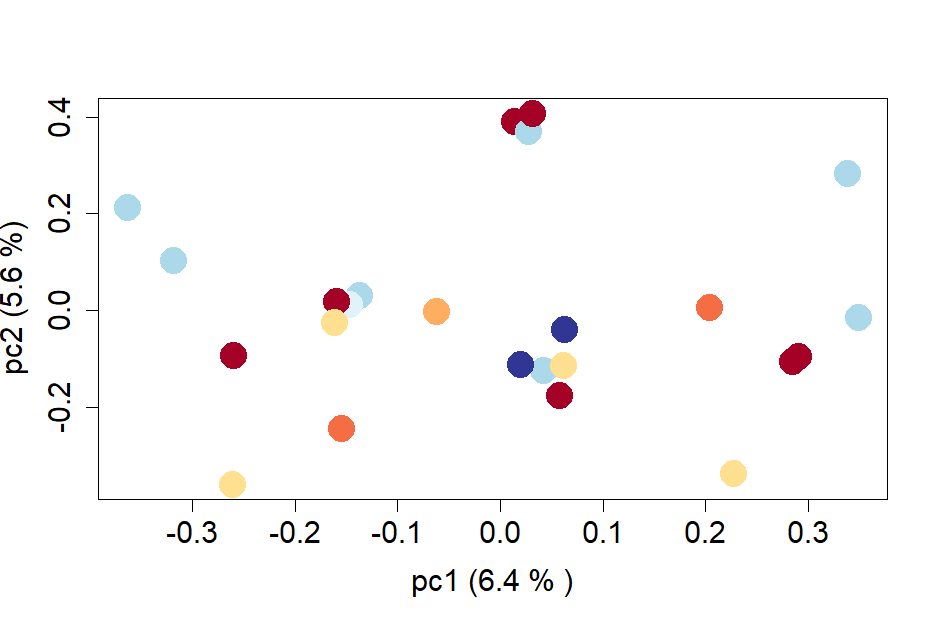 | | | **B:** PCA plot - West coast Wolf herring samples  Number of SNPs – 4188  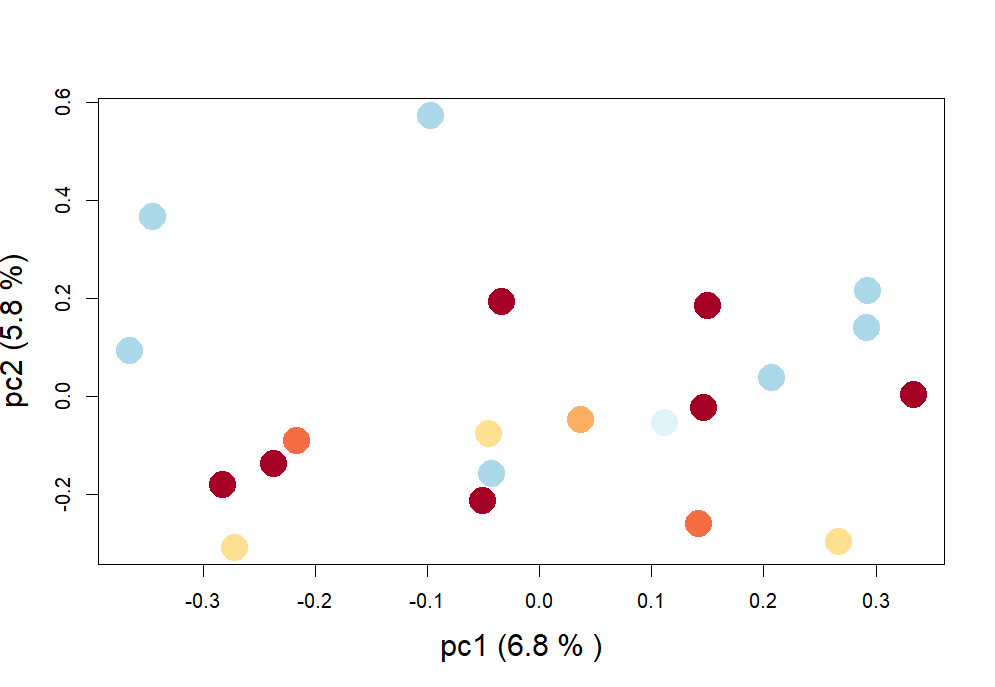 | |
| **Admixture plots (Best K is identified by black border)** | | | | | | |
| **C:** Admixture plots – west coast Indian mackerel samples with Odisha (east coast) | | | CV errors | **D:** Admixture plots – west coast Indian mackerel samples | | CV errors |
| 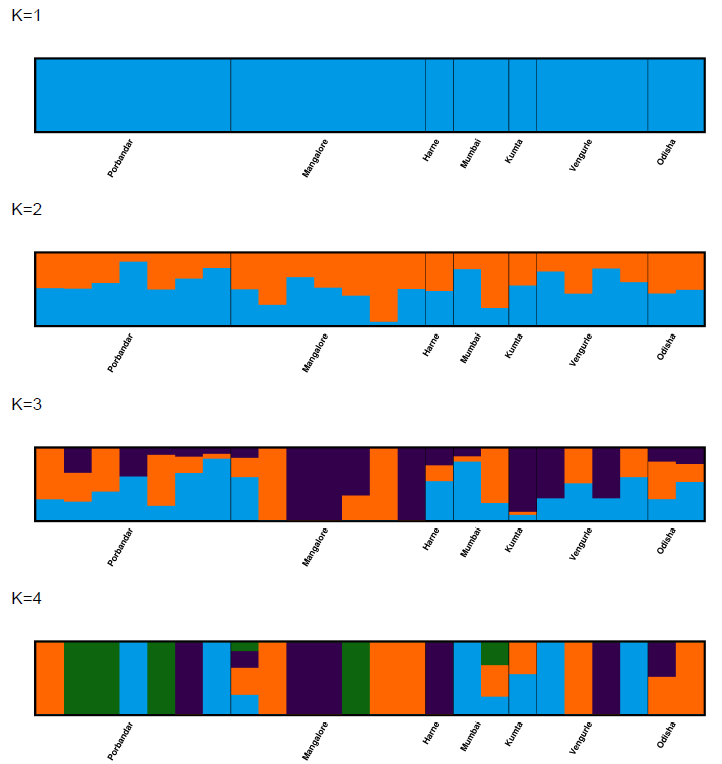 | | | 0.5729  0.7715  0.9838  1.199 | 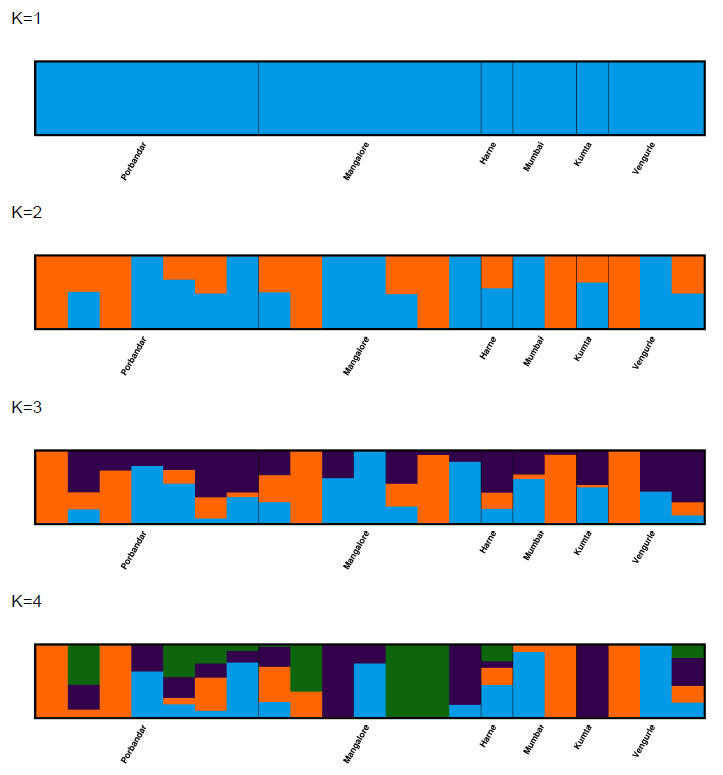 | | 0.62  0.84  1.1  1.37 |
| **Pairwise fixation index (FST) matrix for samples from each site** | | | | | | |
| **E:**  **F:** | 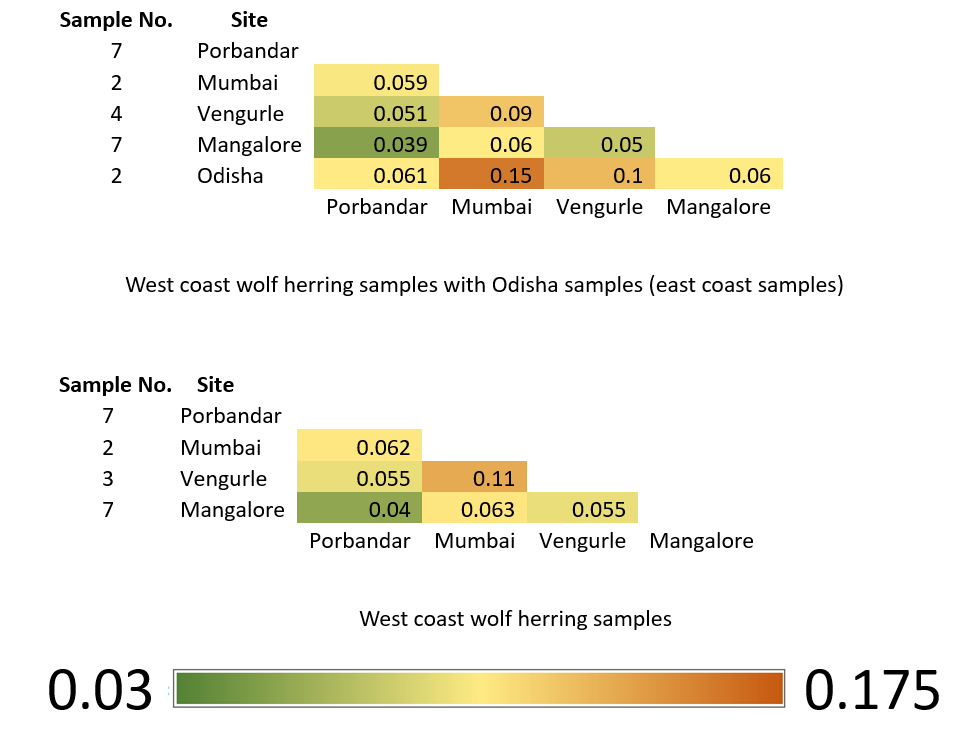 | | | | | |

3. **Spiny cheek grouper (*Epinephelus diacanthus*)**

Our findings indicate no signature of population structure or differentiation within the Spiny check grouper samples on the west coast of India. PCA analysis demonstrates that the Spiny check grouper exhibits a panmictic population (Fig. S7A). These conclusions were further supported by Admixture analysis, where the best-supported model was found to be K=1 (Fig. S7B), along with pair-wise Fst values showing no differentiation (Fig. S7C). We did not find this species along the east coast so we could not include samples from east coast for this analysis.

**Figure S7. Genetic population structure results of Spiny cheek grouper**

| 3. *Epinephelus diacanthus (Spinycheek grouper)*   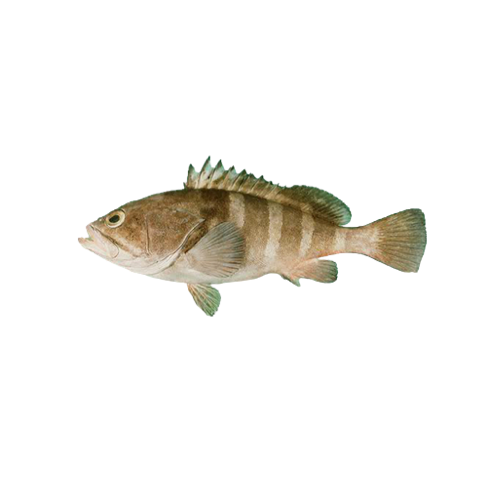 | | | |
| --- | --- | --- | --- |
| **Sampling locations:** Map of sample points and Legend of PCA plots  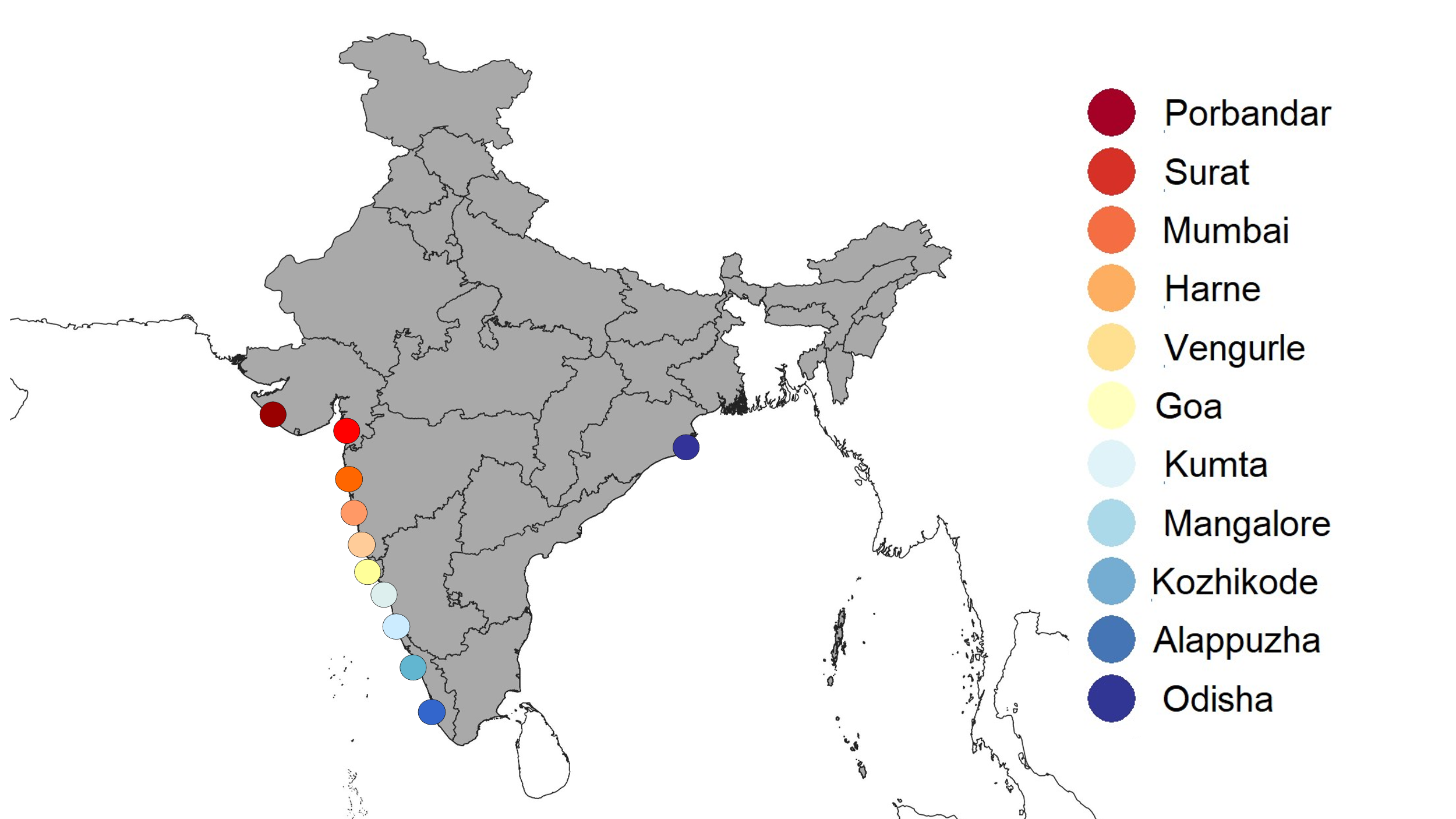 | **A:** PCA plot - West coast Spiny cheek grouper samples, Number of SNPs – 5252  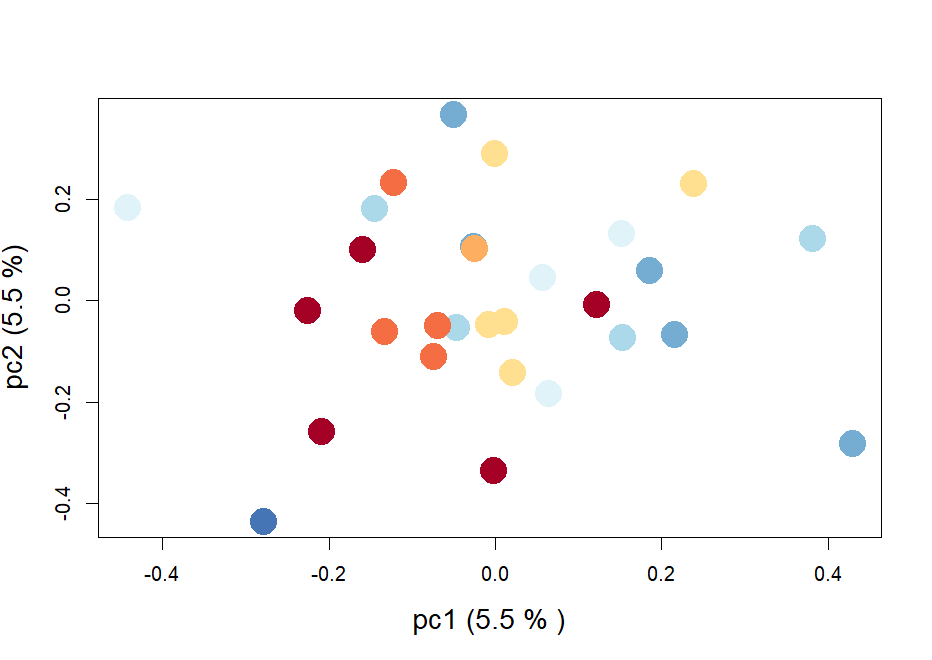 | | |
| **B:** Admixture plots – west coast Spiny cheek grouper samples | | CV errors | **C:** Pairwise fixation index (FST) matrix for samples from each site |
| 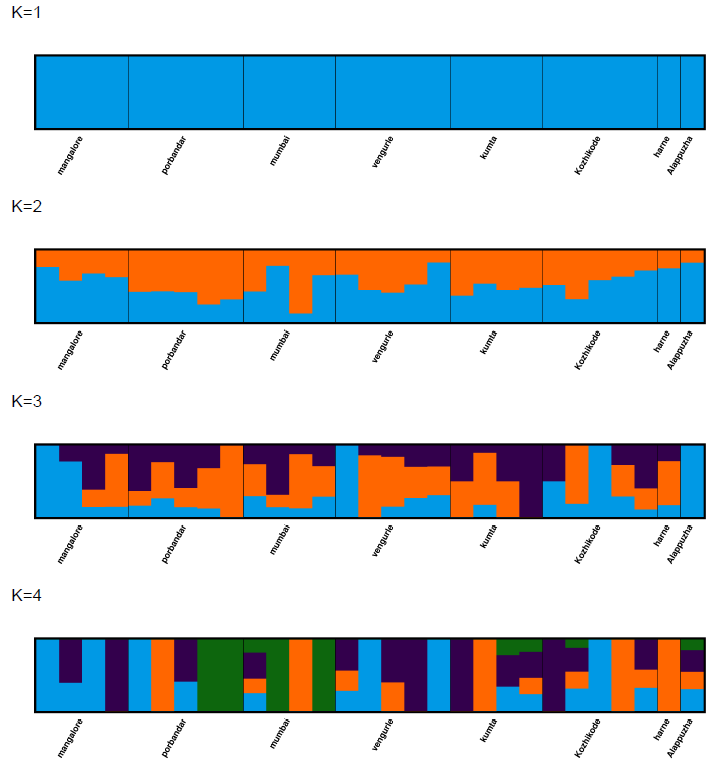 | | 0.5295  0.6614  0.7992 | 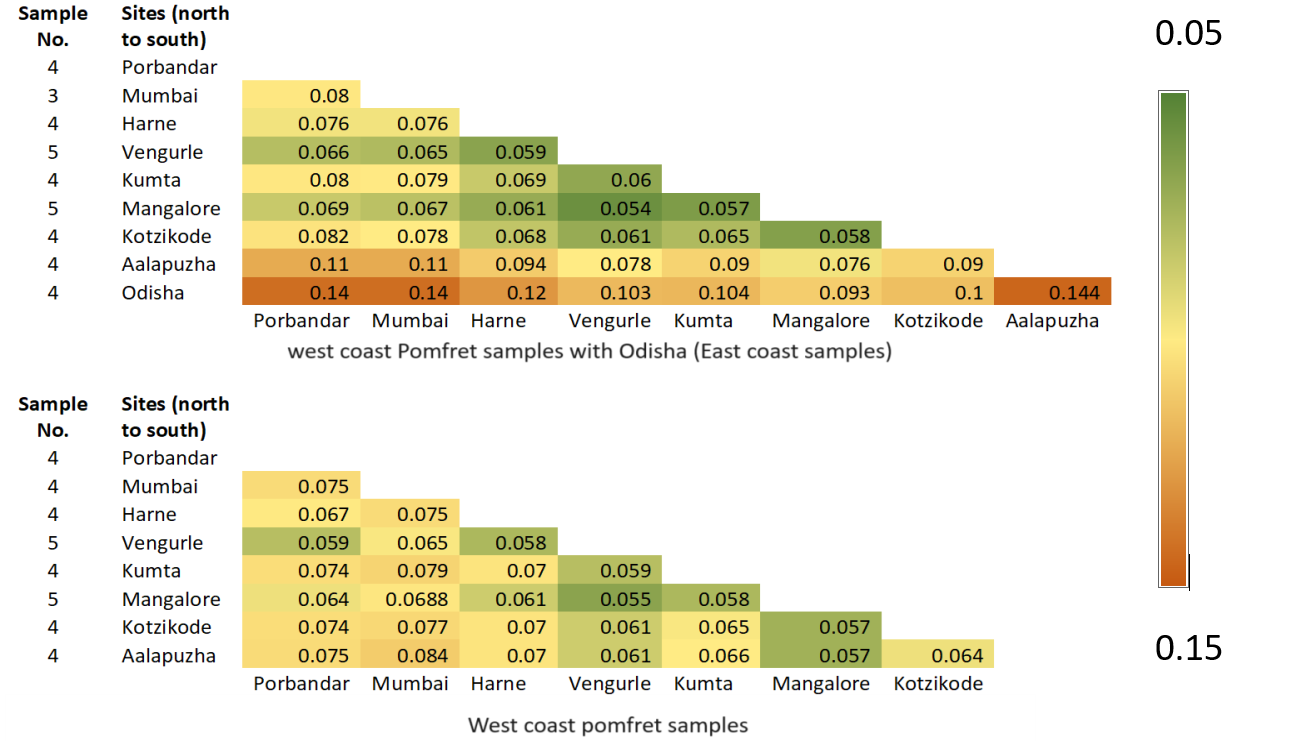  0.0  0.1  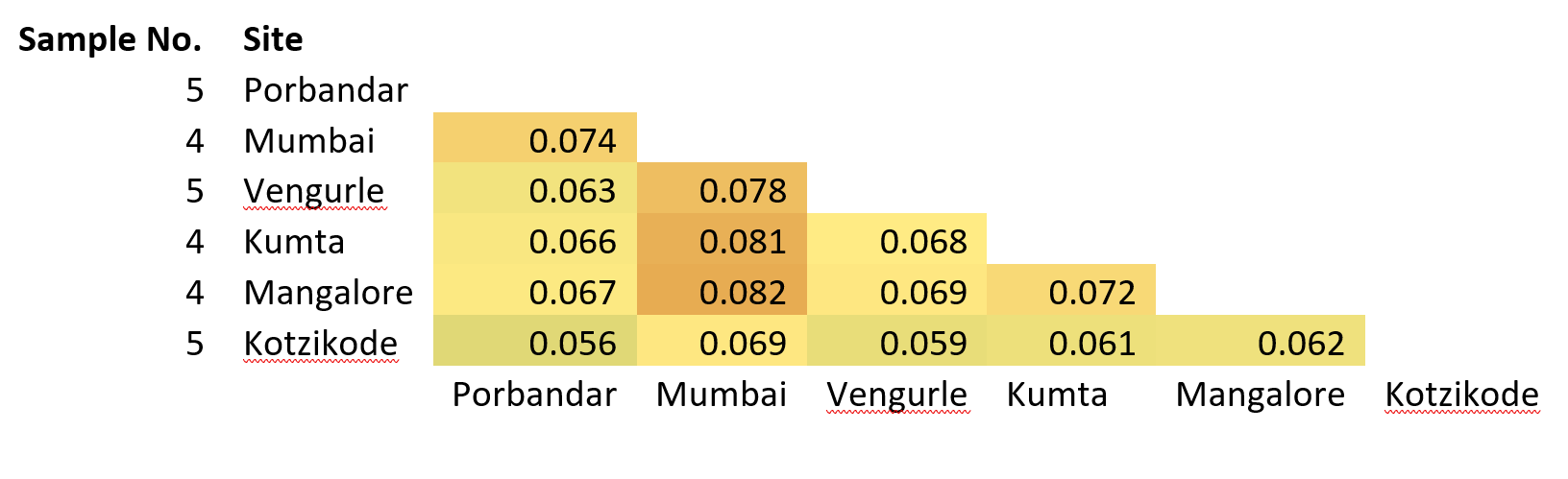  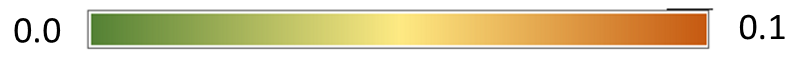 |

4. **Ribbonfish (*Trichiurus lepturus*)**

Our results highlight that Ribbonfish exhibit very strong structure and genetic differentiation between the east and west coasts of India. All three results, PCA (Fig.S8A), admixture (best supported model is K=2) (Fig.S8C) and pairwise Fst (Fig. S8E) shows a clear differentiation of the east coast (Odisha) samples.

However, our findings indicate no signature of population structure or differentiation within the Ribbonfish samples along the west coast of India. PCA analysis demonstrates that the Ribbonfish exhibit a panmictic population (Fig. S8B). These conclusions were further supported by Admixture analysis, where the best-supported model was found to be K=1 (Fig. S8D), along with pair-wise Fst values showing no differentiation (Fig. S8F).

**Figure S8. Genetic population structure results of Ribbonfish**

| 4.*Trichiurus lepturus* (Ribbonfish)  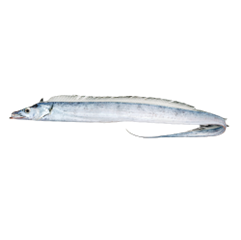 | | | | | | |
| --- | --- | --- | --- | --- | --- | --- |
| **Sampling locations:** Map of sample points and Legend of PCA plots  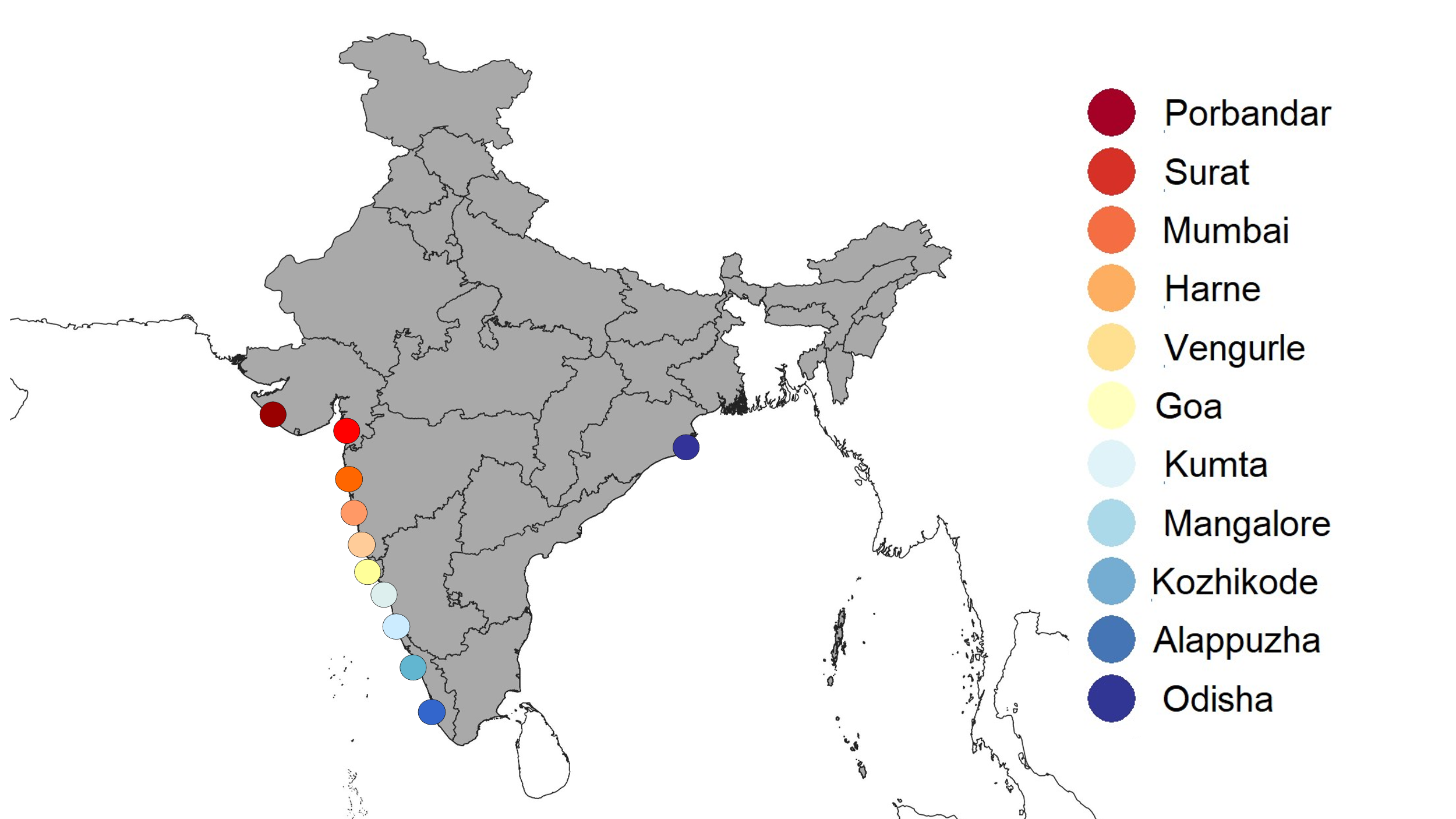 | | **A:** - PCA plot - West coast Pomfret samples with Odisha  No Of SNPs – 5076  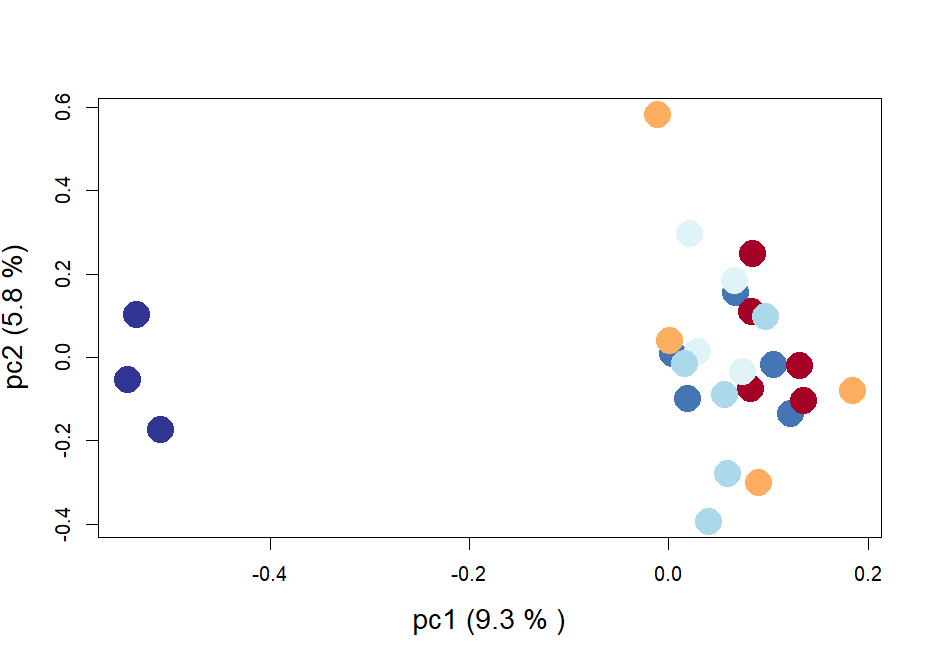 | | | B: PCA plot - West coast pomfret samples  Number of SNPs – 9960  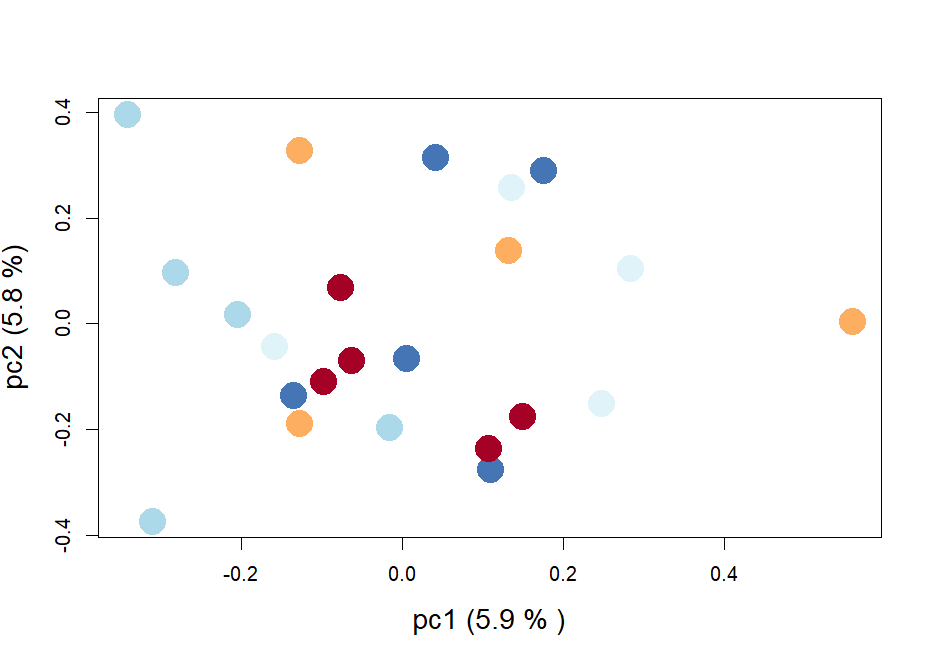 | |
| **Admixture plots (Best K is identified by black border)** | | | | | | |
| **C:** Admixture plots – west coast ribbon samples with Odisha (east coast) | | | CV errors | **D:** Admixture plots – west coast ribbon samples | | CV errors |
| 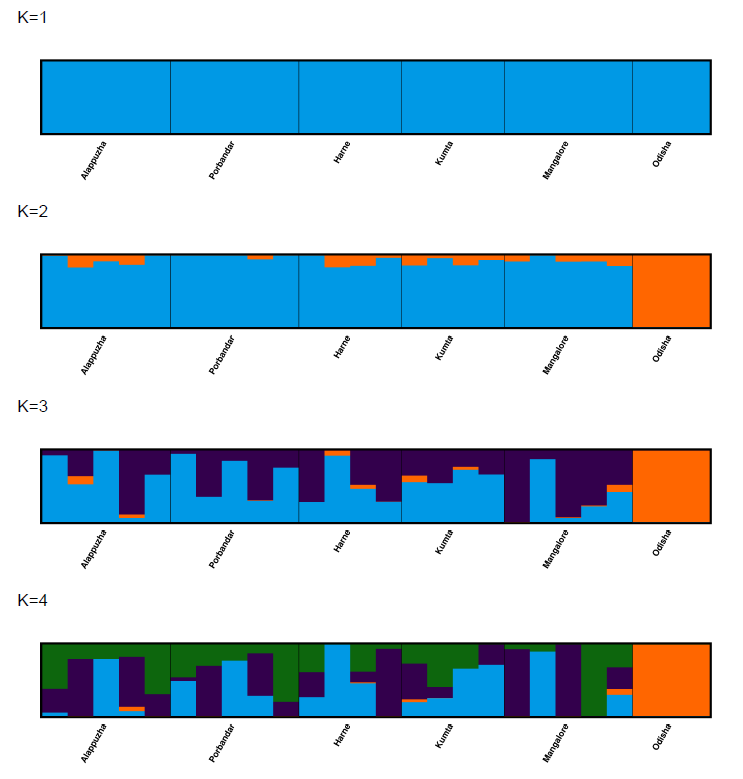 | | | 0.5879  0.7340  0.9447  1.1082 | 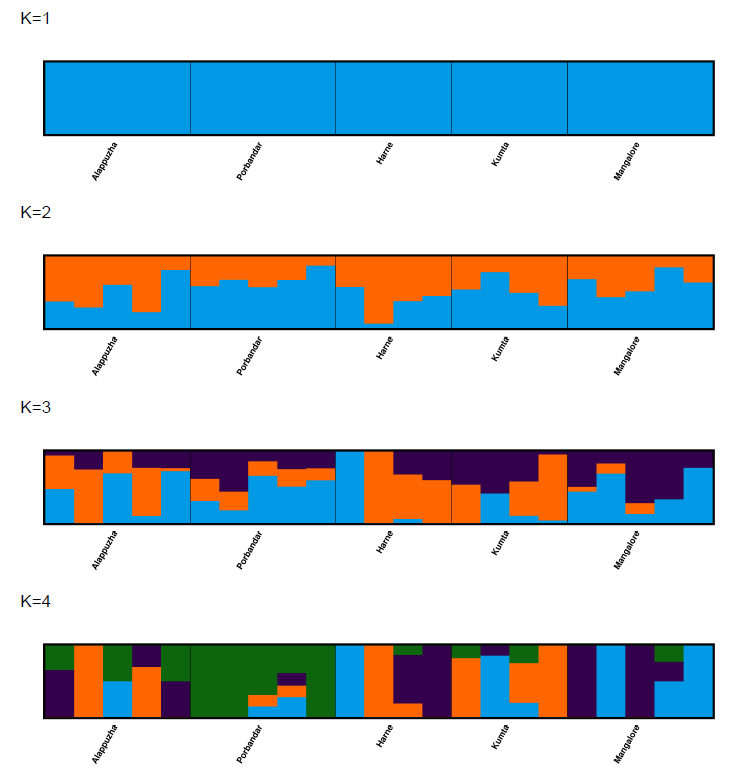 | | 0.6101  0.8150  0.9951  1.2444 |
| **Pairwise fixation index (FST) matrix for samples from each site** | | | | | | |
| **E:**  **F:** | 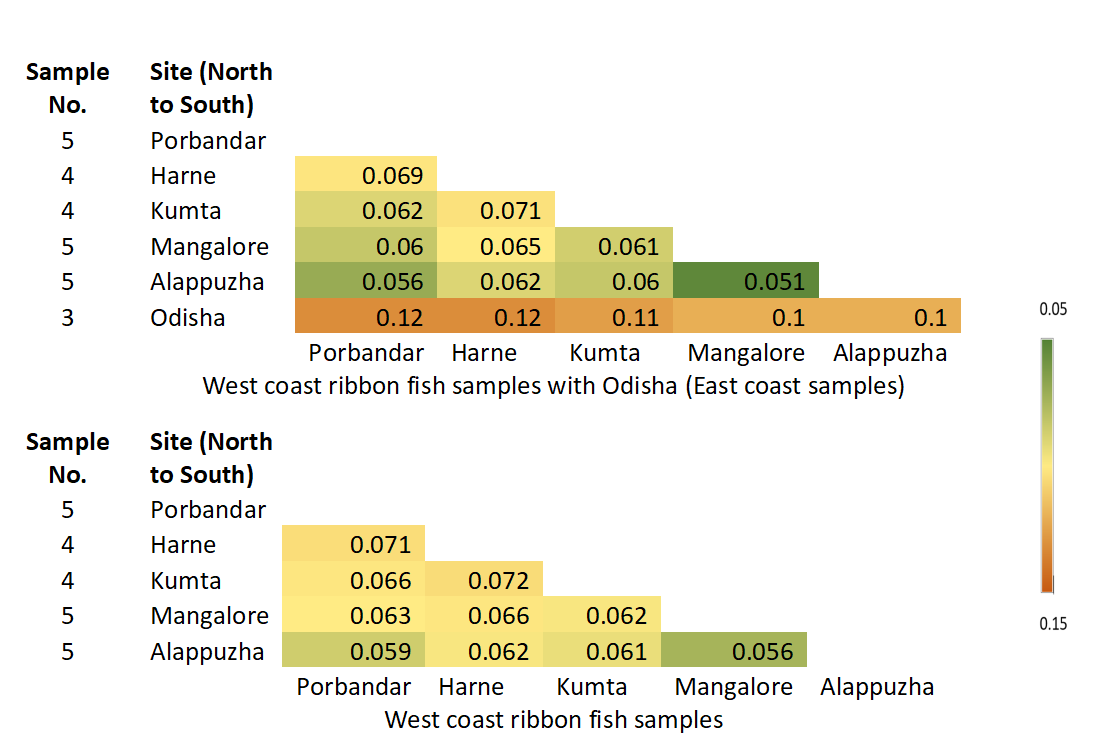 | | | | | |

5. **Japanese threadfin bream (*Nemipterus japonicus*)**

Our results highlight that Ribbonfish exhibit very strong structure and genetic differentiation between the east and west coasts of India. All three results, PCA (Fig.S9A), admixture (best supported model is K=2) (Fig. S9C) and pairwise Fst (Fig. S9E) show a clear differentiation of east coast (Odisha) samples.

However, our findings indicate no signature of population structure or differentiation within the Japanese threadfin bream samples along the west coast of India. PCA analysis demonstrates that the Japanese threadfin bream exhibits a panmictic population (Fig. S9B). These conclusions were further supported by Admixture analysis, where the best-supported model was found to be K=1 (Fig. S9D), along with pair-wise Fst values showing no differentiation (Fig. S9F).

**Figure S9. Genetic population structure results of Japanese threadfin bream**

| 5. *Nemipterus japonicus* (Japanese threadfin bream)  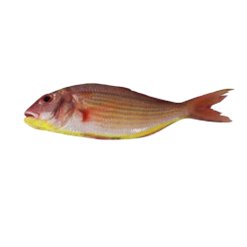 | | | | | | |
| --- | --- | --- | --- | --- | --- | --- |
| **Sampling locations:** Map of sample points and Legend of PCA plots  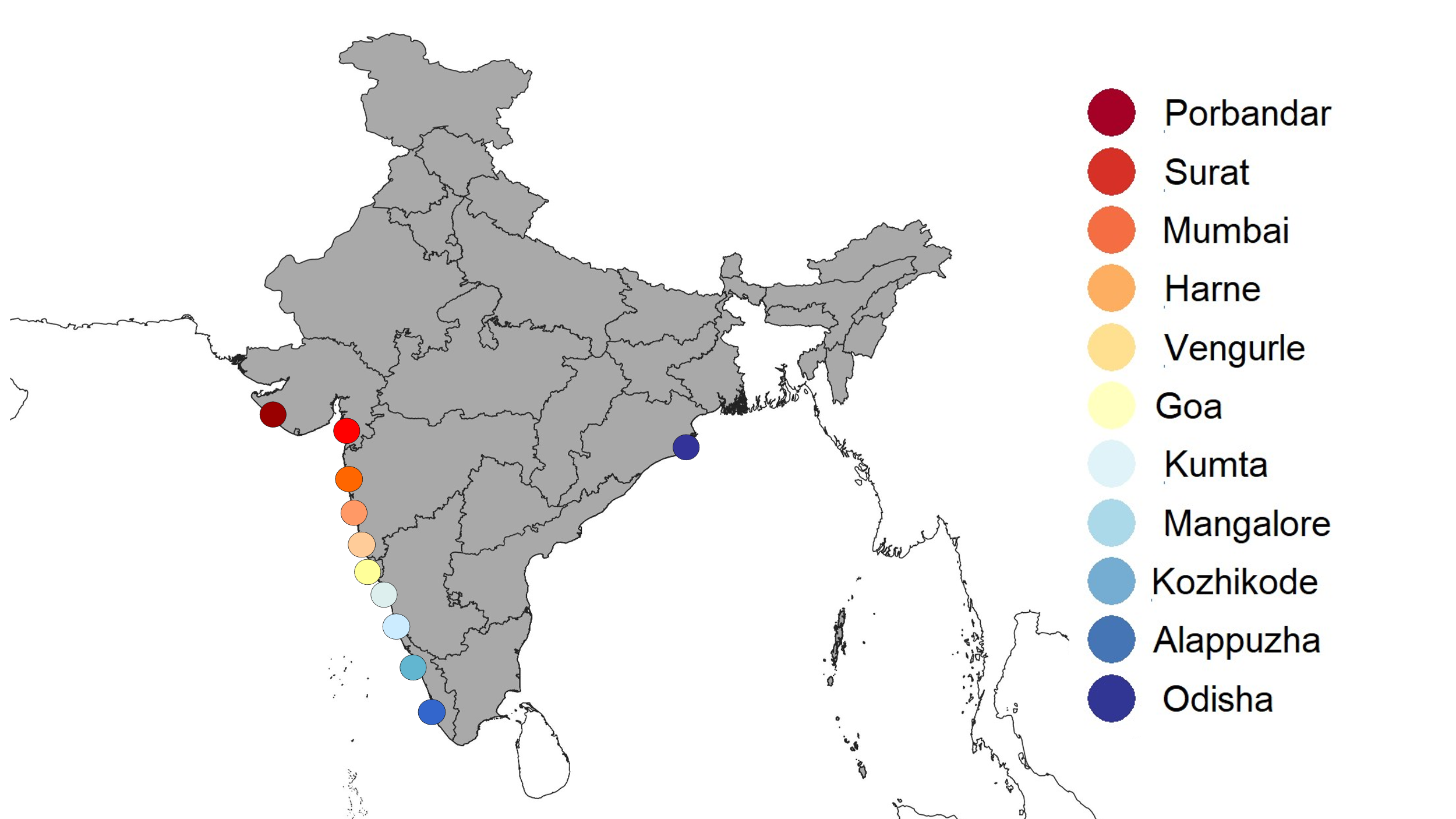 | | **A:** PCA plot - West coast Japanese threadfin bream samples with Odisha  No Of SNPs – 9824  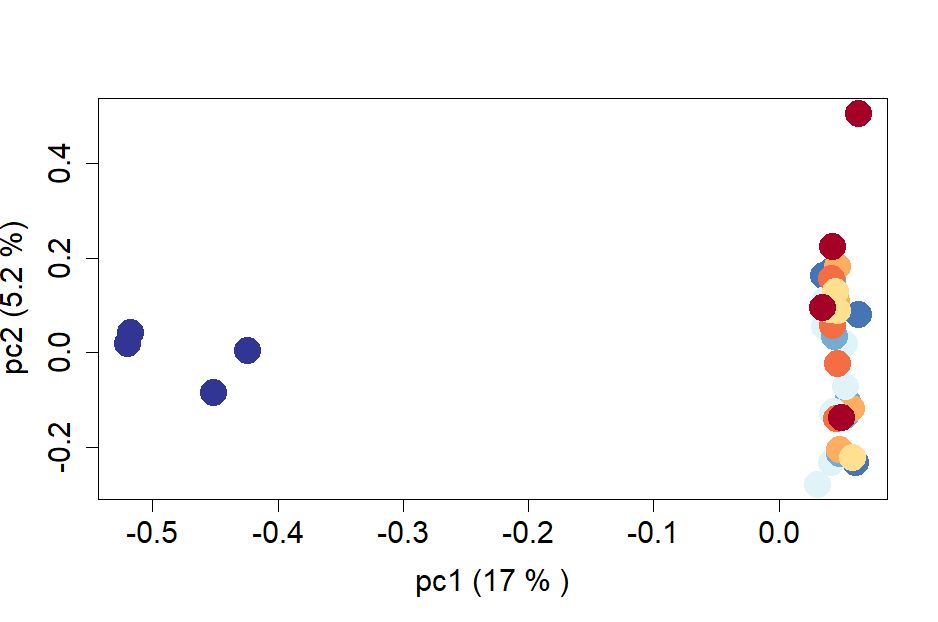 | | | **B:** PCA plot - West coast Japanese threadfin bream samples  Number of SNPs – 4774  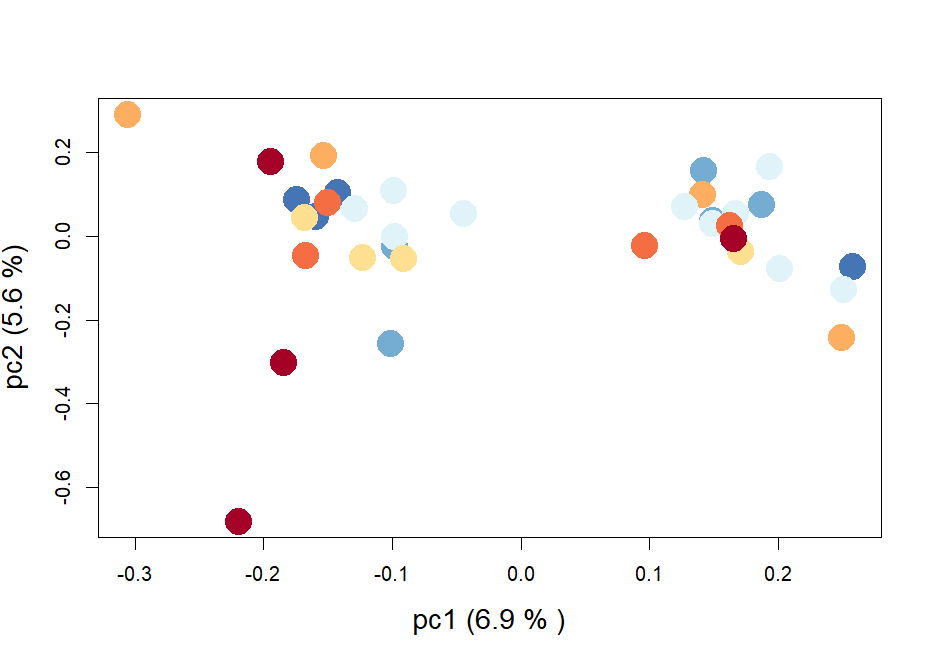 | |
| **Admixture plots (Best K is identified by black border)** | | | | | | |
| **C:** Admixture plots – west coast Japanese threadfin bream samples with Odisha (east coast) | | | CV errors | **D:** Admixture plots – west coast Japanese threadfin bream samples | | CV errors |
| 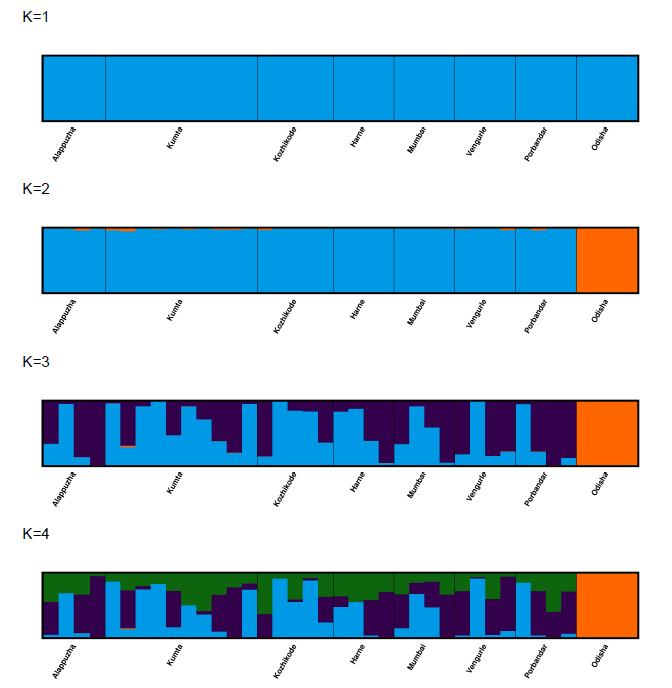 | | | 0.5116  0.5735  0.6494  0.7584 | 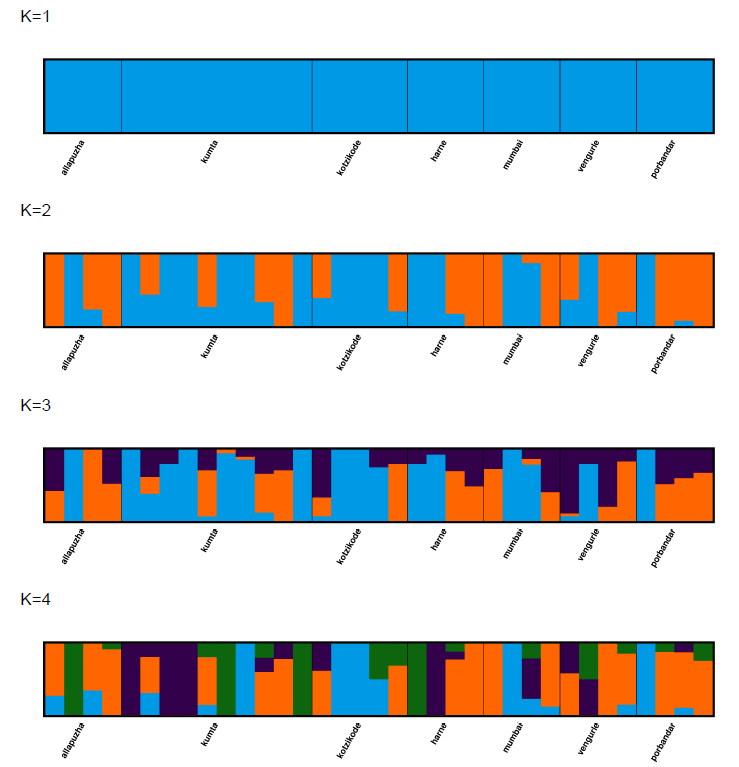 | | 0.4964  0.5706  0.6766  0.7986 |
| **Pairwise fixation index (FST) matrix for samples from each site** | | | | | | |
| **E:**  **F:** | 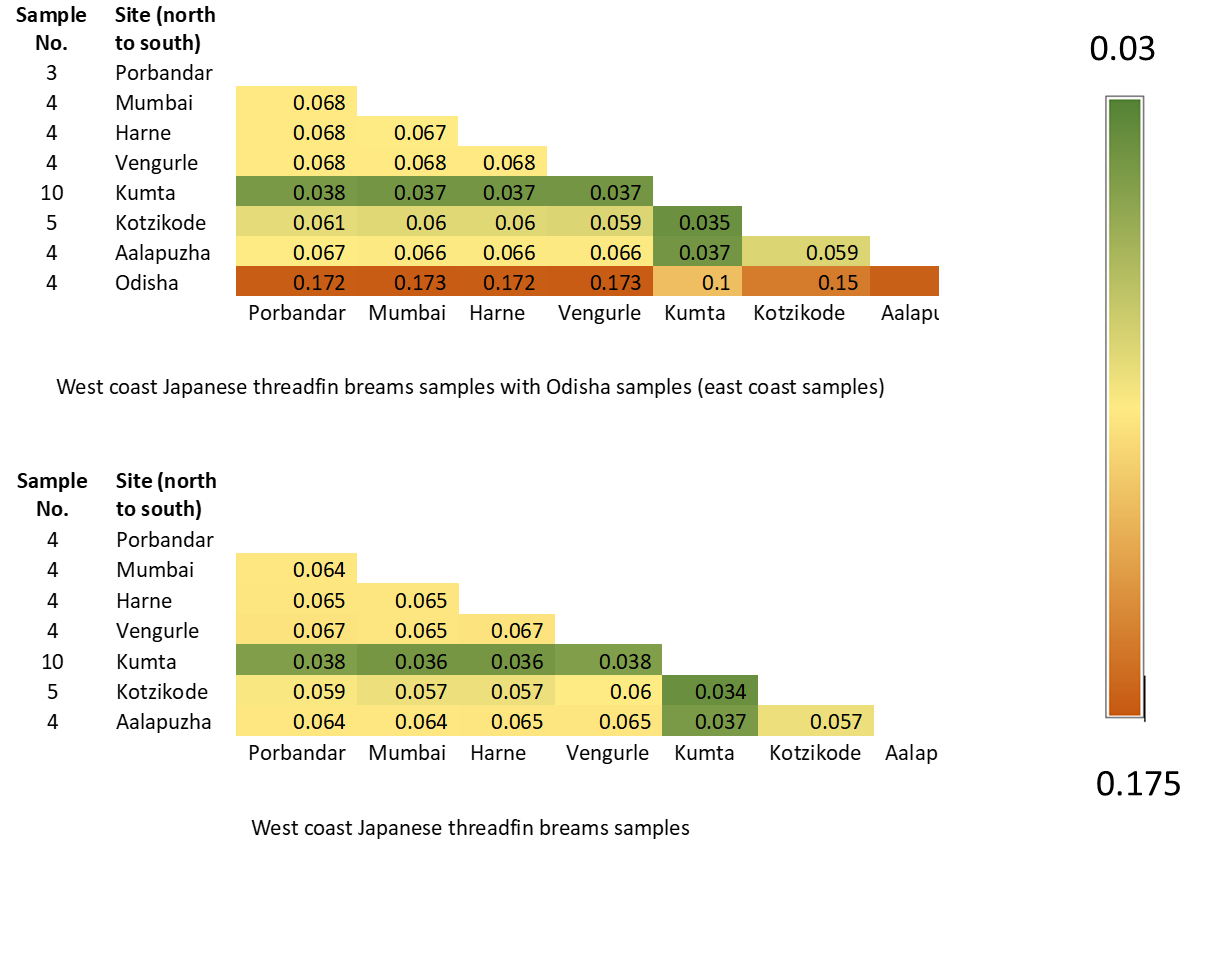 | | | | | |

6. **Silver Pomfret (*Pampus argenteus*)**

Our results highlight that silver pomfret exhibit very strong structure and genetic differentiation between east and west coasts of India. All three results, PCA (Fig.S10A), admixture (Fig.S10C) and pairwise Fst (Fig.S10E) show a clear differentiation of the east coast (Odisha) samples.

Interestingly, we also observe population structure in this species along the west coast of India. The PCA analysis demonstrates that the silver pomfret exhibit two genetic clusters north and south of Vengurle (Fig. S10B). These conclusions were further supported by Admixture analysis, where the best-supported model was found to be K = 2 (Fig. S10D), along with pair-wise Fst values showing slight differentiation (Fig. S10F). There are few samples from Vengurle at the junction of two clusters, which are clubbing together with the southern samples in FSt, increasing their chances to be part of the southern population.

**Figure S10. Genetic population structure results of Silver pomfret**

| 7. *Pampus argenteus* (Silver Pomfret)   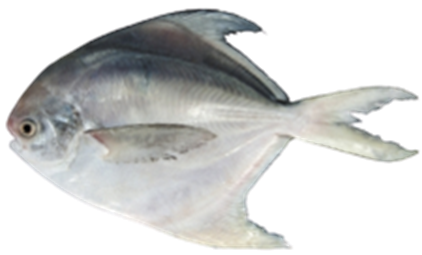 | | | | | | |
| --- | --- | --- | --- | --- | --- | --- |
| **Sampling locations:** Map of sample points and Legend of PCA plots  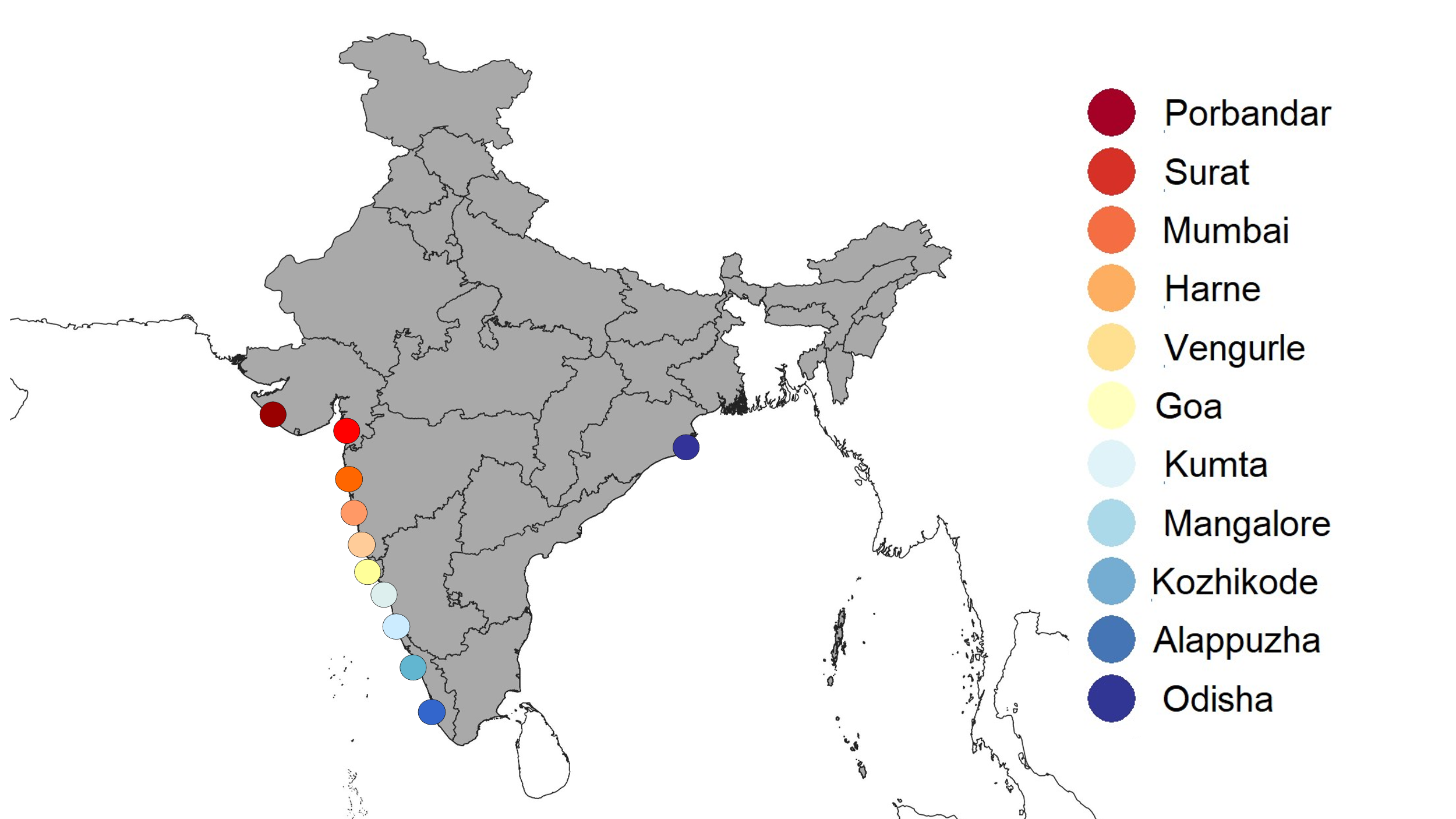 | | **A:** PCA plot - West coast Silver Pomfret samples with Odisha  No Of SNPs – 4995    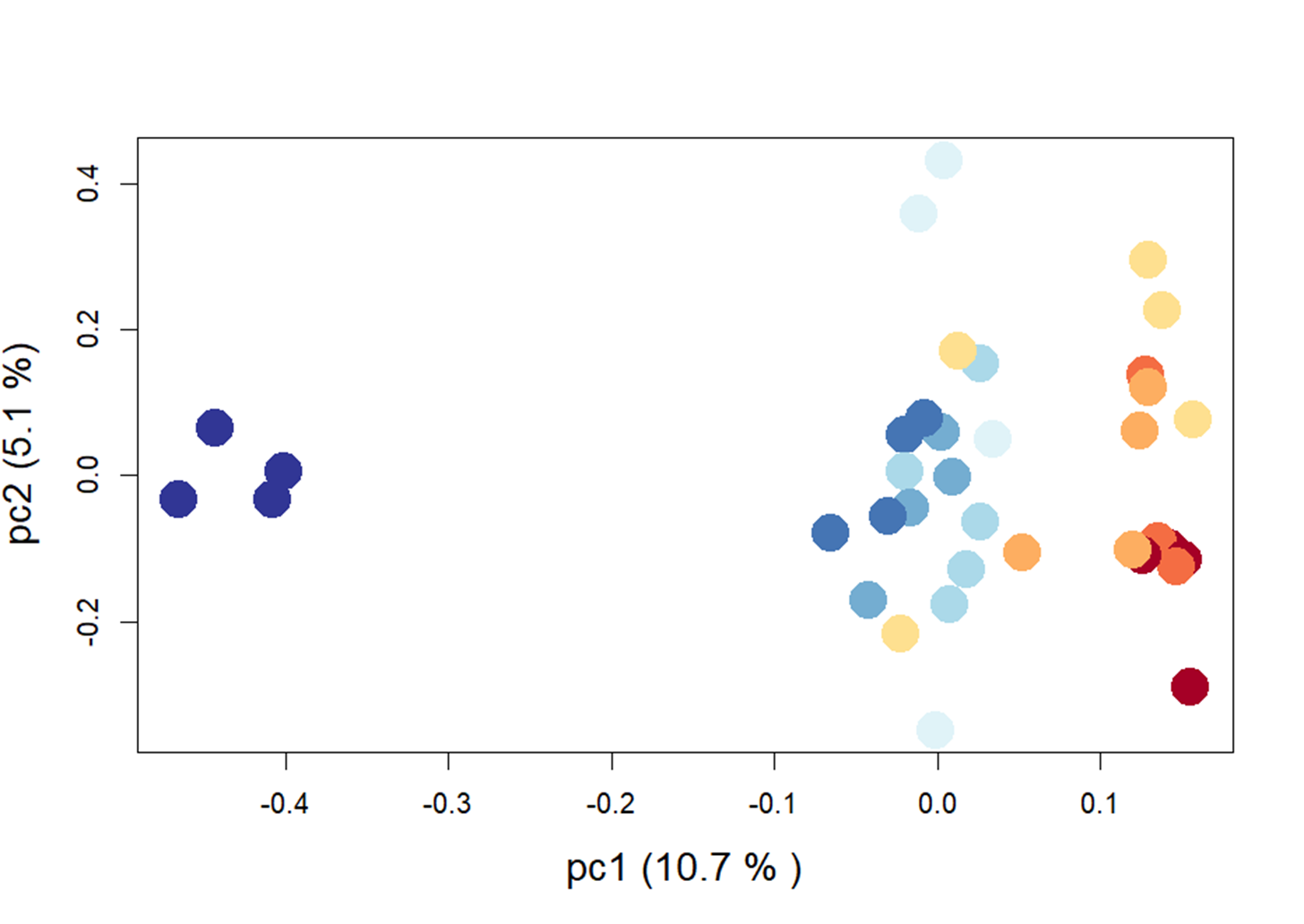 | | | B: PCA plot - West coast Silver Pomfret samples  Number of SNPs – 5068  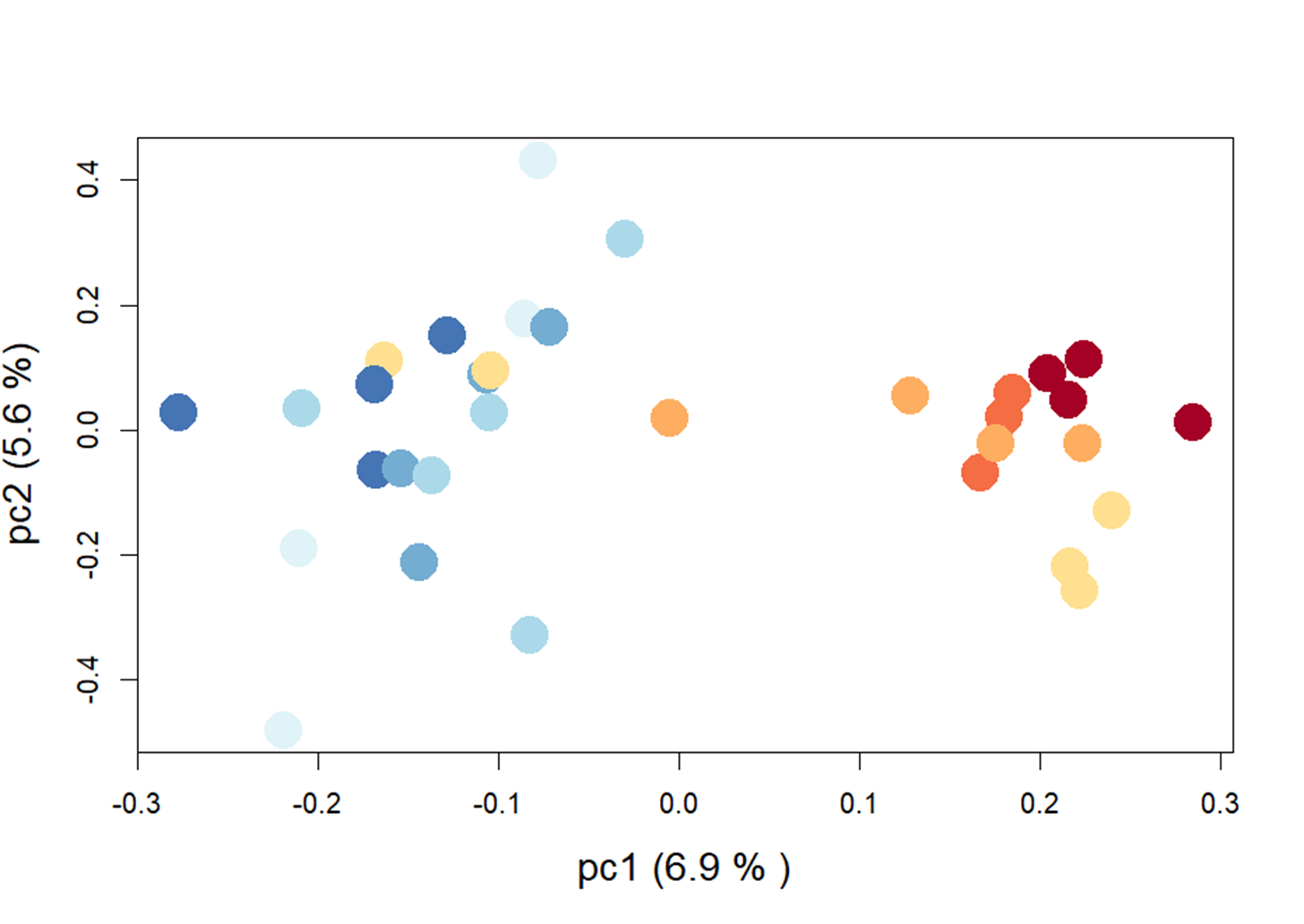 | |
| **Admixture plots (Best K is identified by black border)** | | | | | | |
| **C:** Admixture plots – west coast silver pomfret samples with Odisha (east coast) | | | CV errors | **D:** Admixture plots – west coast silver pomfret samples | | CV errors |
| 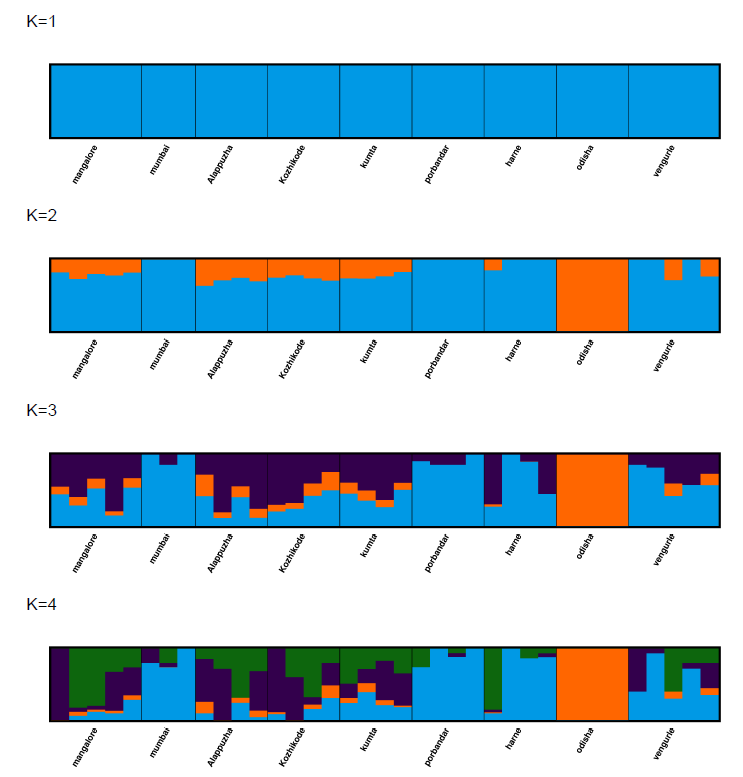 | | | 0.4619  0.5069  0.5966  0.7170 | 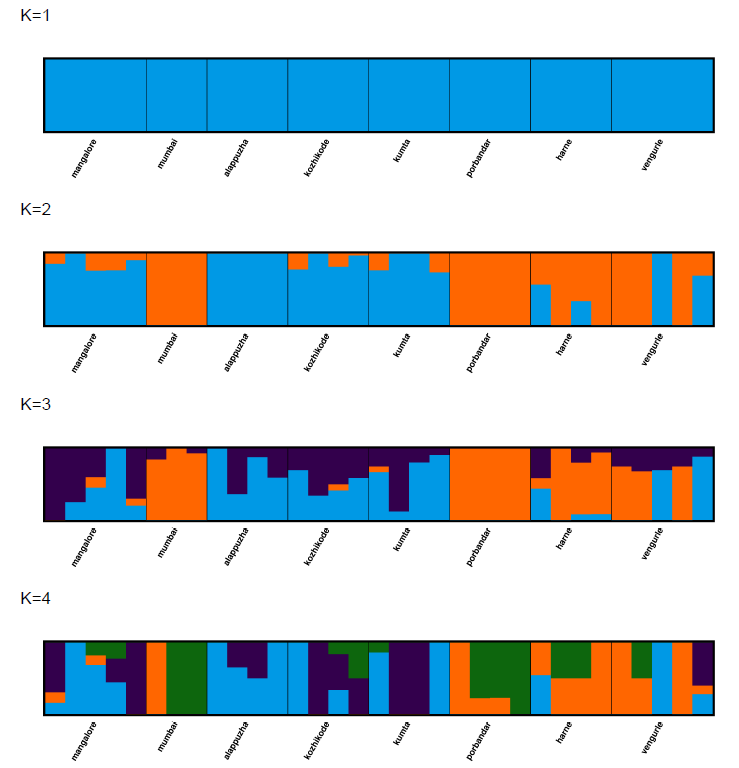 | | 0.4821  0.5814  0.7005  0.8129 |
| **Pairwise fixation index (FST) matrix for samples from each site** | | | | | | |
| **E:**  **F:** | 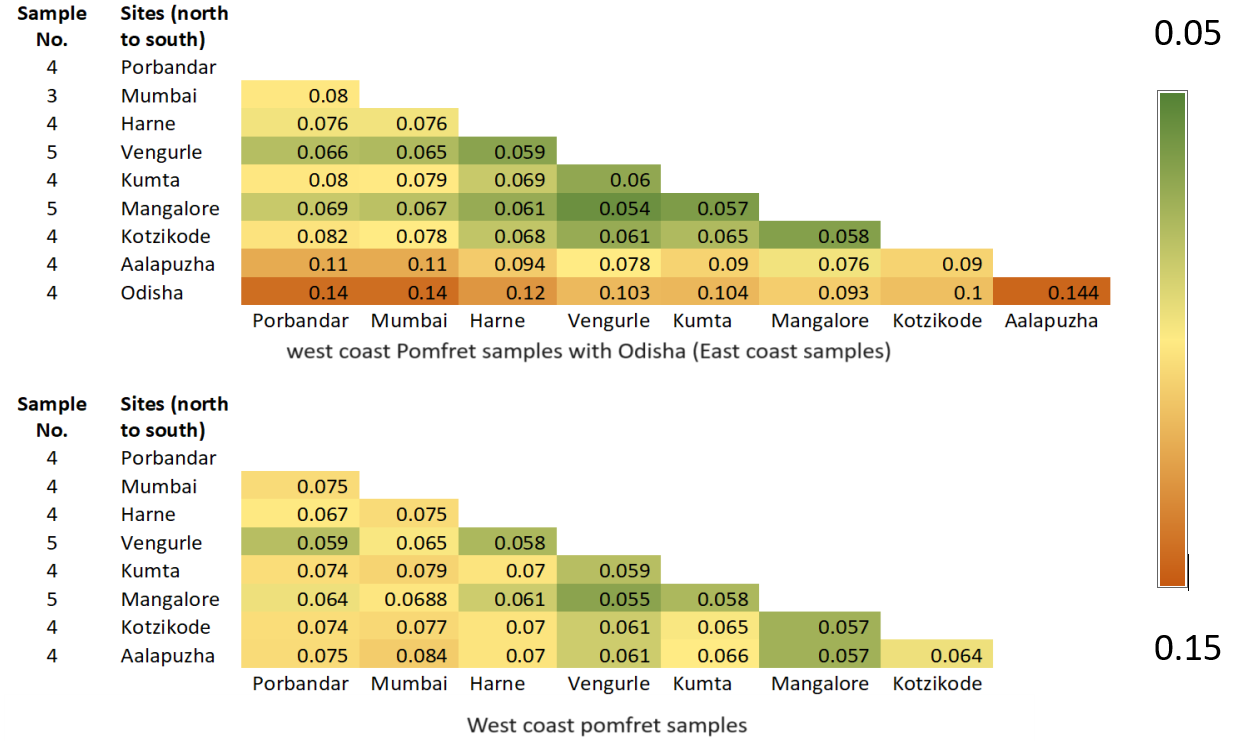 | | | | | |

7. **Indian oil sardine (*Sardinella longiceps*)**

Our results indicate that Indian oil sardines exhibit very strong structure and genetic differentiation between Gujarat and the rest of west coast samples. The PCA shows two distinct populations (Fig.S11A). These conclusions were further supported by Admixture analysis, where the best-supported model was found to be K = 2 (Fig.S11B) and pairwise Fst (Fig.S11C), also shows a clear differentiation of Gujarat samples from rest of the west coast samples. We did not find this species along the east coast, as a result comparative analysis of east coast and west coast could not be conducted.

**Figure S11. Genetic population structure results of Indian Oil sardines**

| 7. *Sardinella longiceps (Indian oil sardine)*  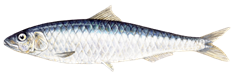 | | | |
| --- | --- | --- | --- |
| **Sampling locations:** Map of sample points and Legend of PCA plots  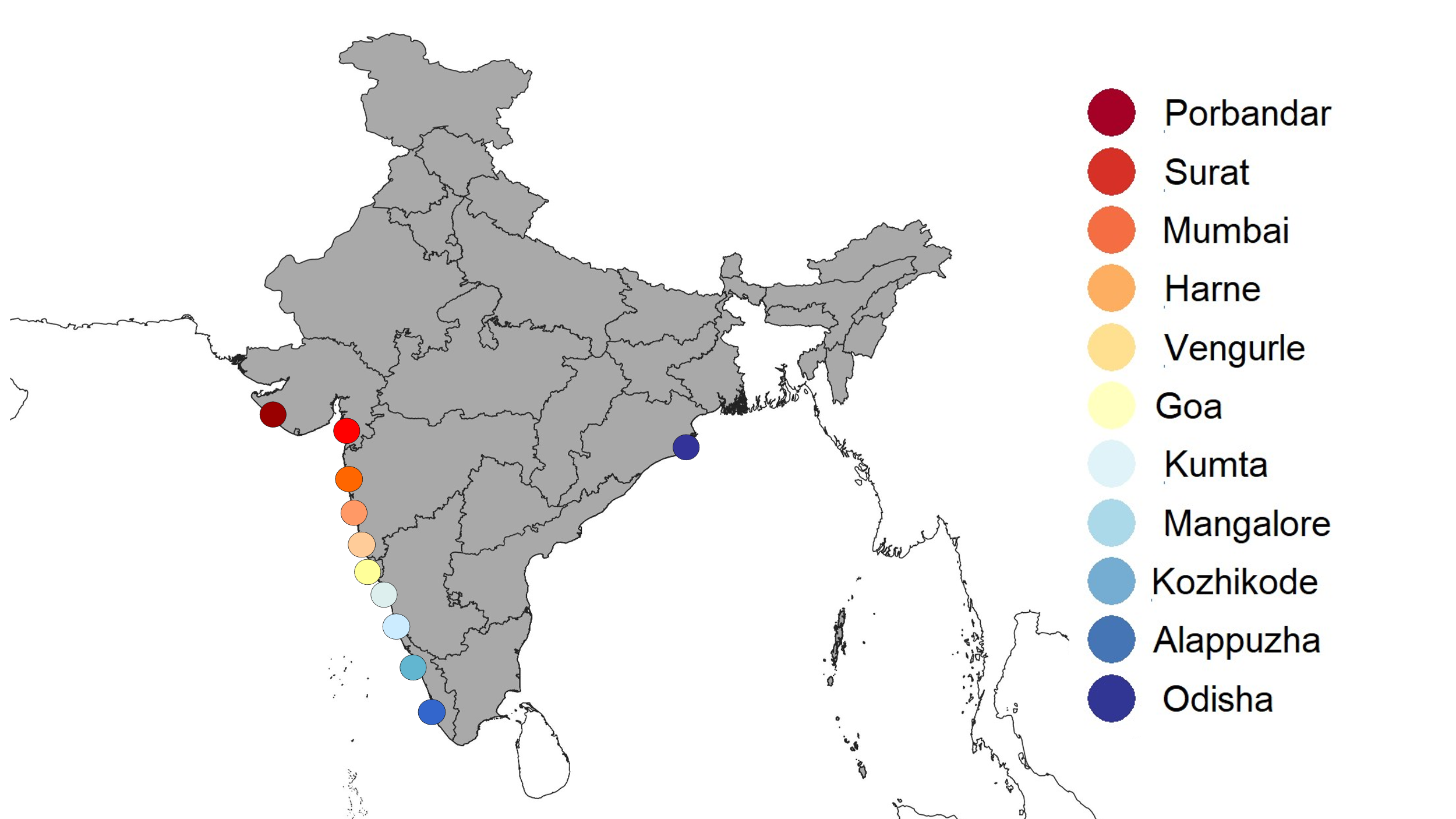 | **A:** PCA plot - West coast Indian oil sardine samples, Number of SNPs – 4470  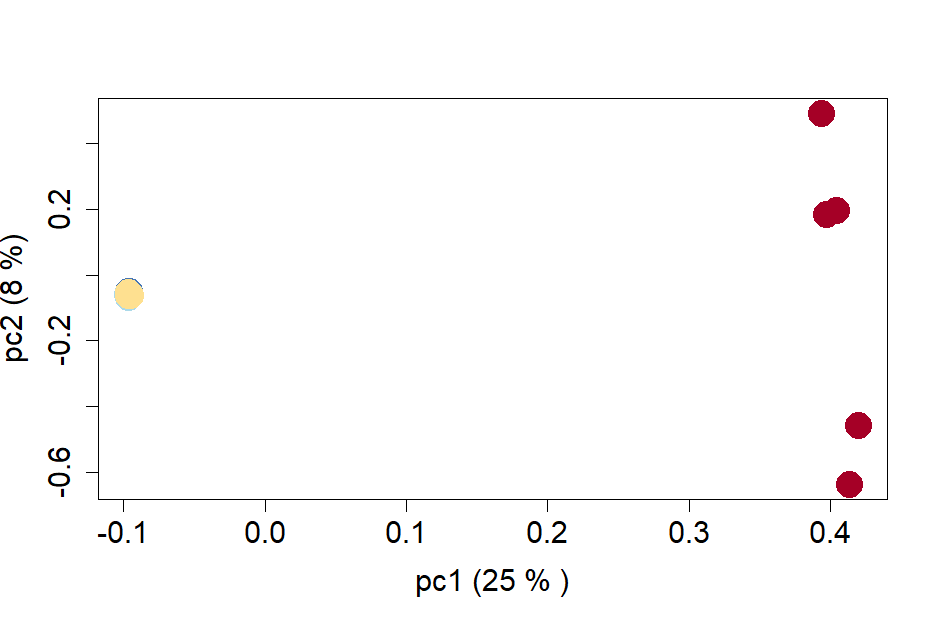 | | |
| **B:** Admixture plots – west coast Indian oil sardine samples | | CV errors | **C:** Pairwise fixation index (FST) matrix for samples from each site |
| 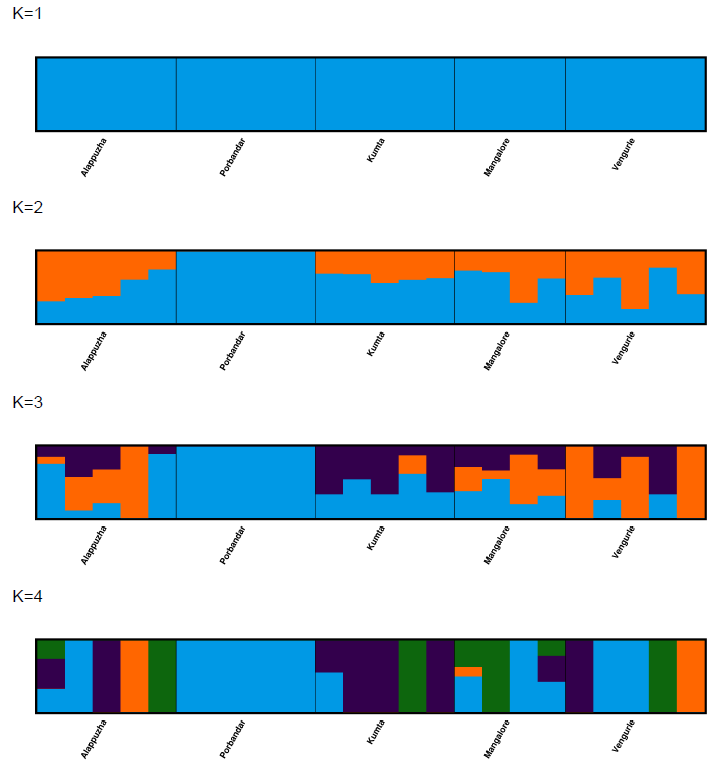 | | 0.6612  0.9094  1.2055  1.6287 | 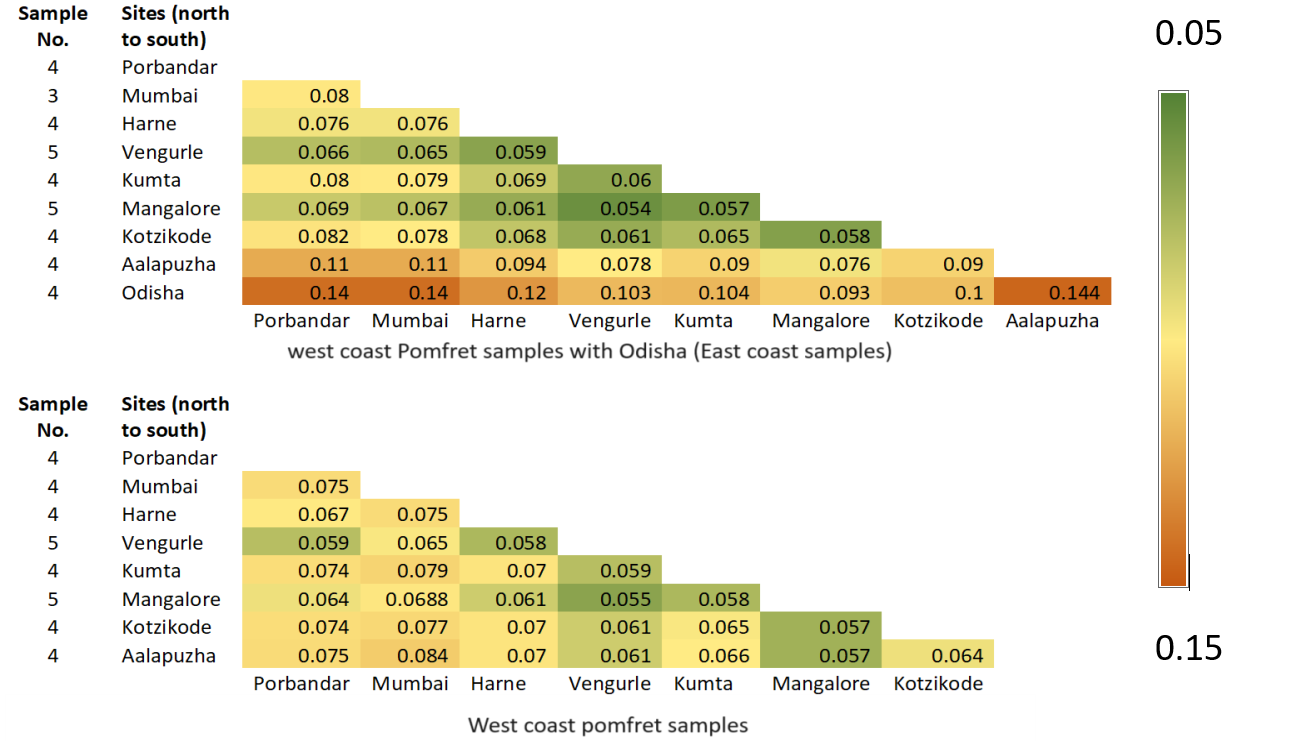  0.0  0.1  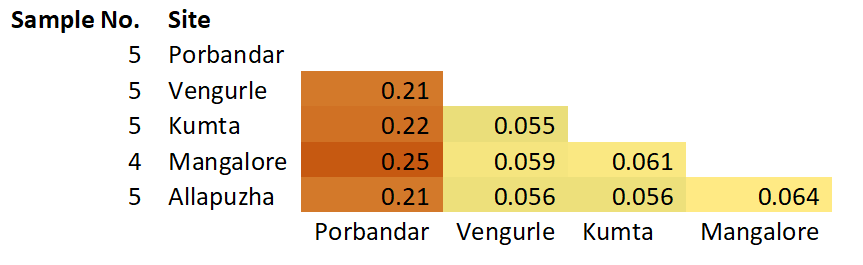  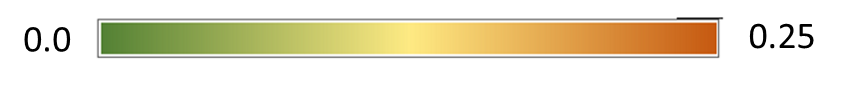 |

8. **Spade nose shark (*Scoliodon laticaudus*)**

Our results indicate that Spade nose shark exhibit very strong structure and genetic differentiation between the Gujarat samples, Maharashtra – Karnataka samples and Kerala samples, making three distinct populations along the west coast of India. PCA (Fig.S12A) and admixture (Best k = 3) (Fig.S12B) show similar results with a couple of Kerala samples featuring in the Gujarat group. Gujarat exports considerable amount of spade nose shark to Kerala which are sometime sold on fish landing sites (Personal Observation). This might explain the observed pattern. Fst results clearly show that Gujarat and Kerala samples are very different from the Maharashtra – Karnataka samples (Fig. S12C).

**Figure S12. Genetic population structure results of Spade nosed shark**

| 9. *Scoliodon laticaudus (Spadenose Shark)*   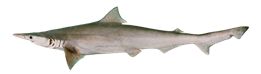 | | | |
| --- | --- | --- | --- |
| **Sampling locations:** Map of sample points and Legend of PCA plots  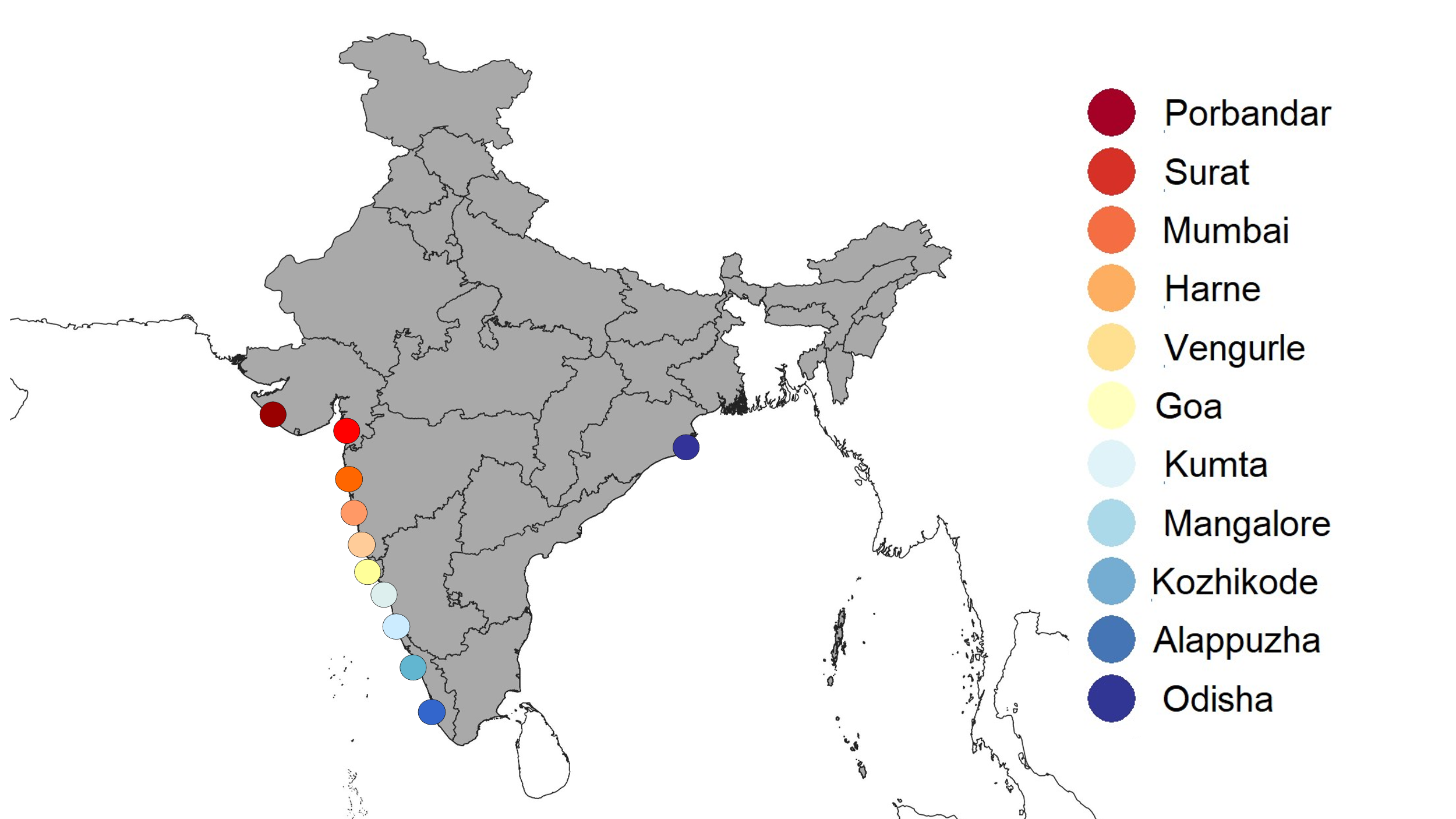 | **A:** PCA plot - West coast Spadenose shark samples, Number of SNPs – 1166  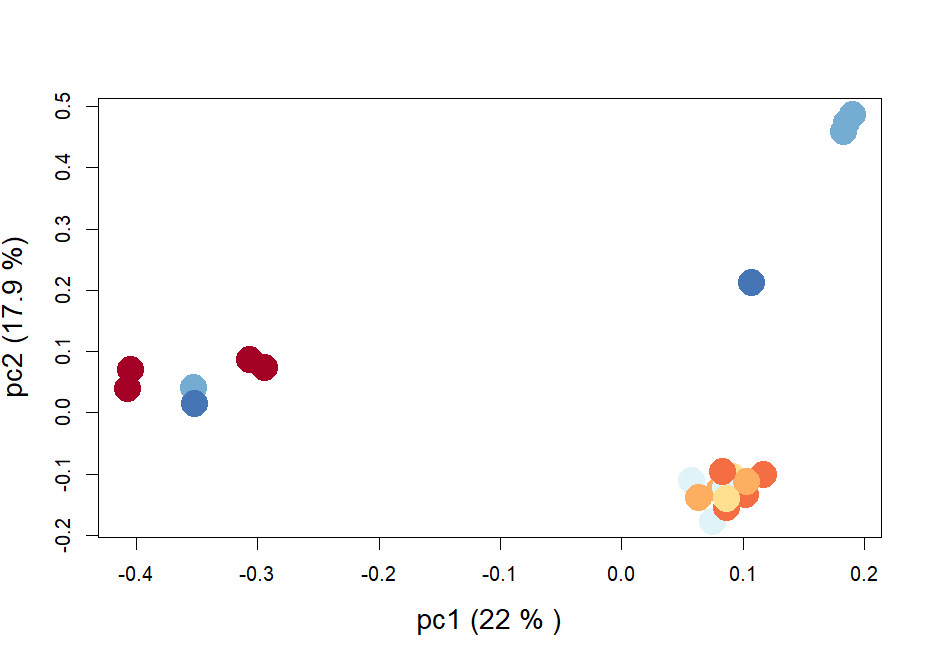 | | |
| **B:** Admixture plots – west coast Spadenose shark samples | | CV errors | **C:** Pairwise fixation index (FST) matrix for samples from each site |
| 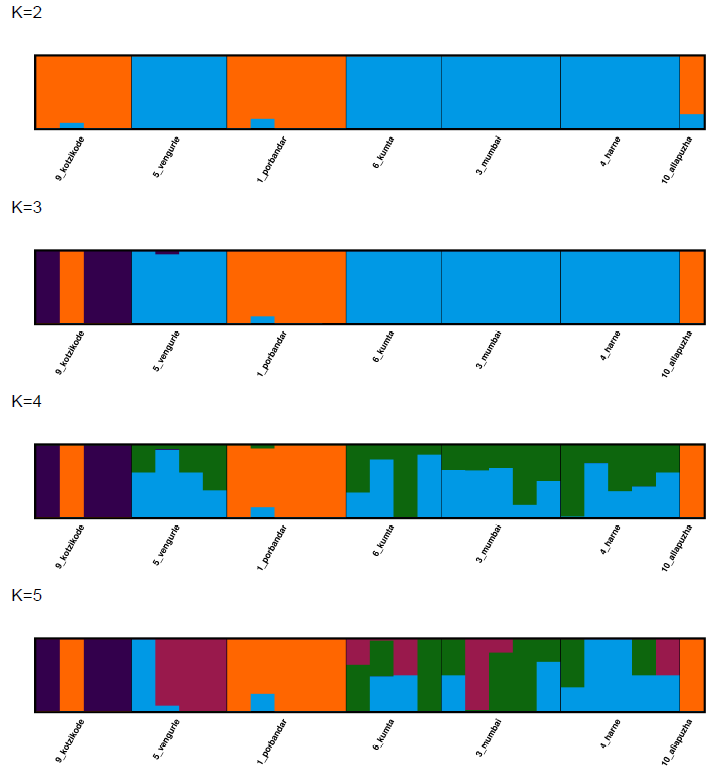 | | 0.5626  0.6303  0.6069  0.7549 | 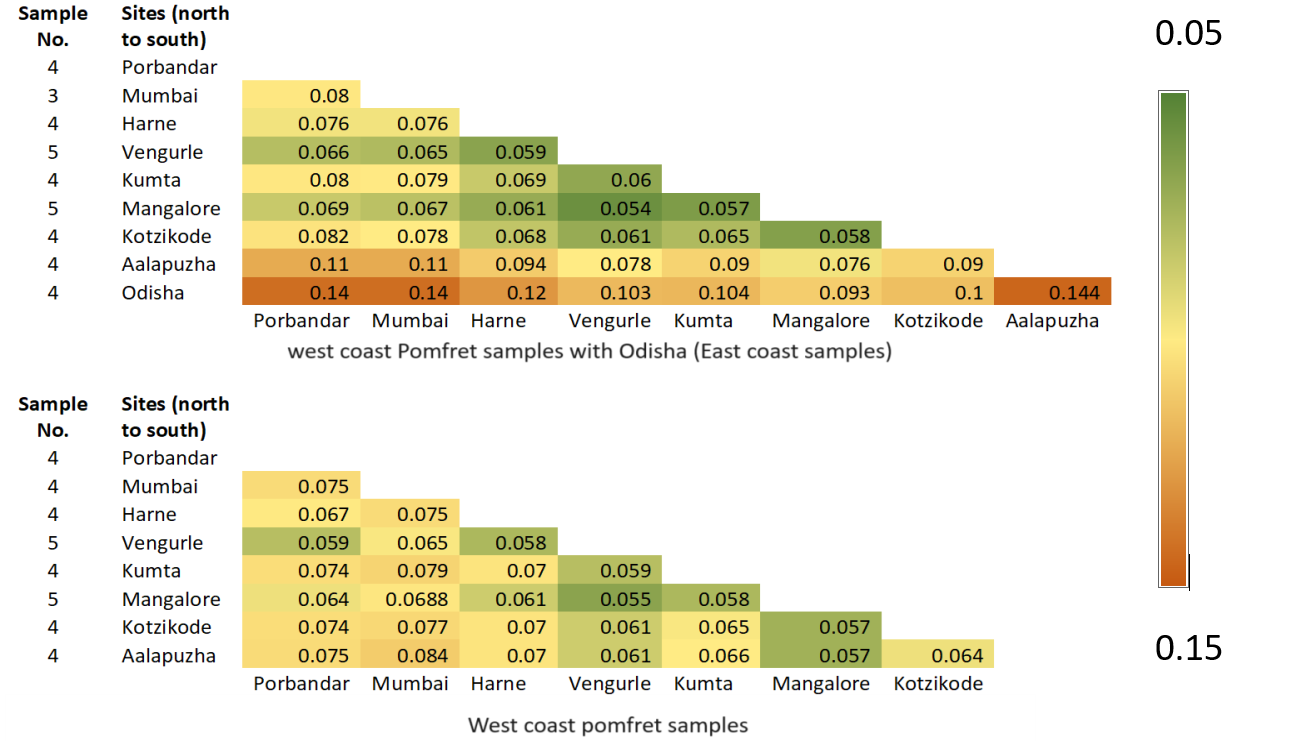  0.0  0.1  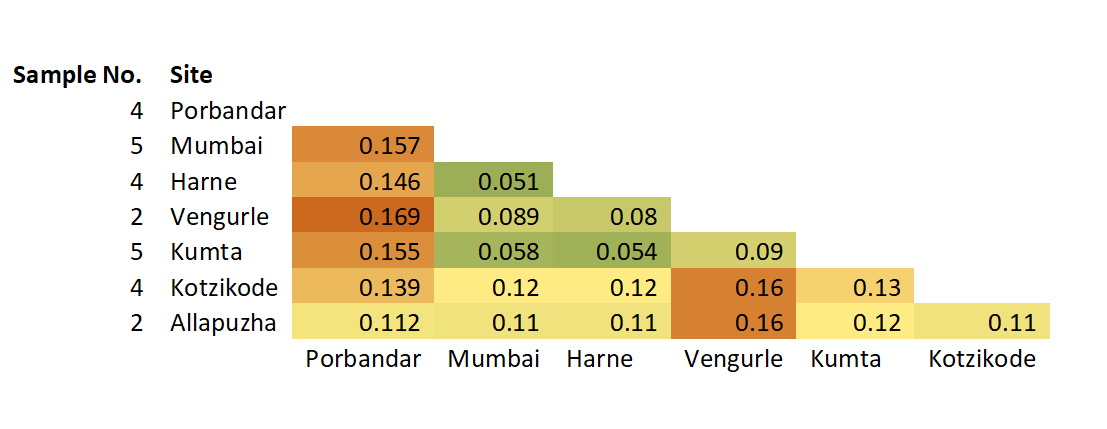  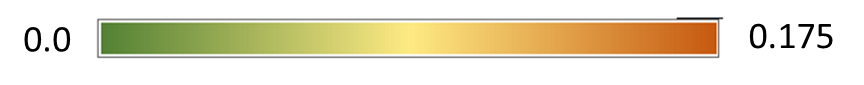 |

**9.Ladyfish / Silver Sillago (*Sillago sihama*)**

Silver silago results are the most unusual of our analysis. Our results indicate that Silver silago exhibit very strong structure and genetic differentiation between samples from Kumta, as compared with the rest of the samples from Maharashtra and Karnataka. The results from the PCA (Fig S13A) Admixture (best K is K =2) (Fig.S13B) and FST (Fig S13C) all shows that the Kumta population clearly separatees from rest of the west coast.

**Figure S13. Genetic population structure results of Silver Silago / Ladyfish**

| 9*. Sillago sihama (Silver Silago)*  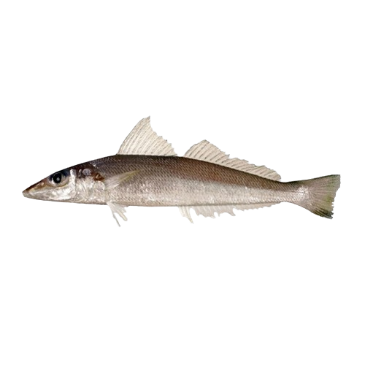 | | | |
| --- | --- | --- | --- |
| **Sampling locations:** Map of sample points and Legend of PCA plots  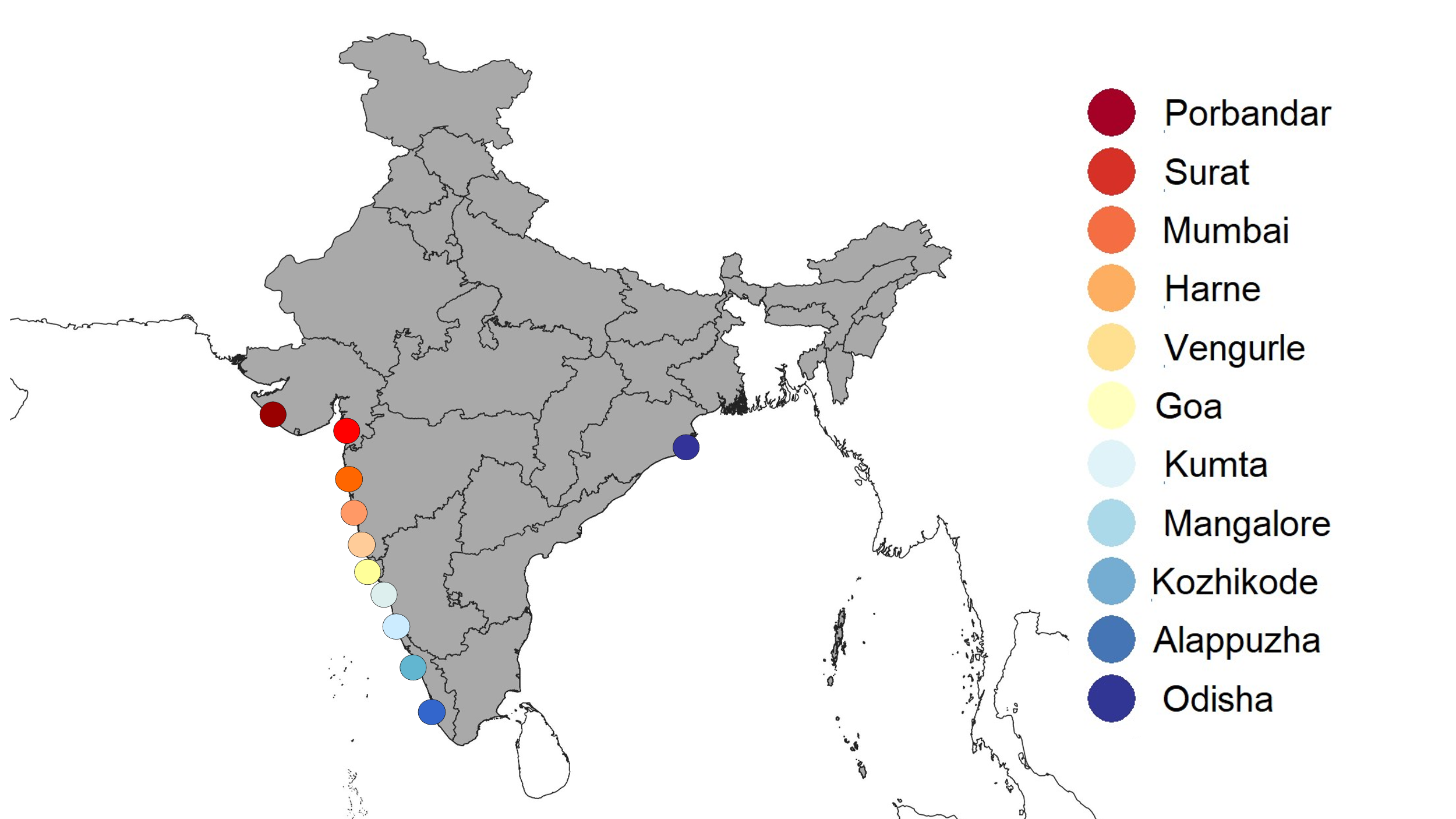 | **A:** PCA plot - West coast Silver silago samples, Number of SNPs – 2755  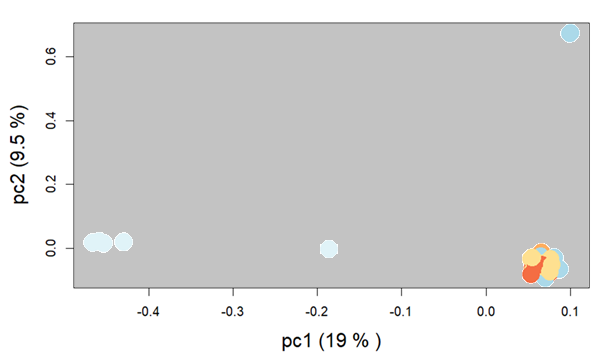 | | |
| **B:** Admixture plots – west coast Silver silago samples | | CV errors | **C:** Pairwise fixation index (FST) matrix for samples from each site |
| 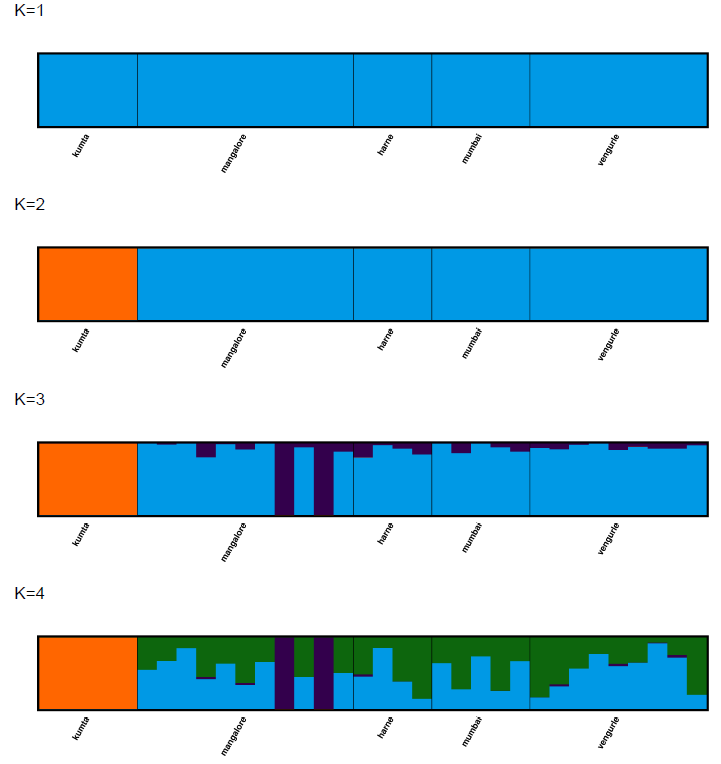 | | 0.5167  0.4464  0.5011  0.6075 | 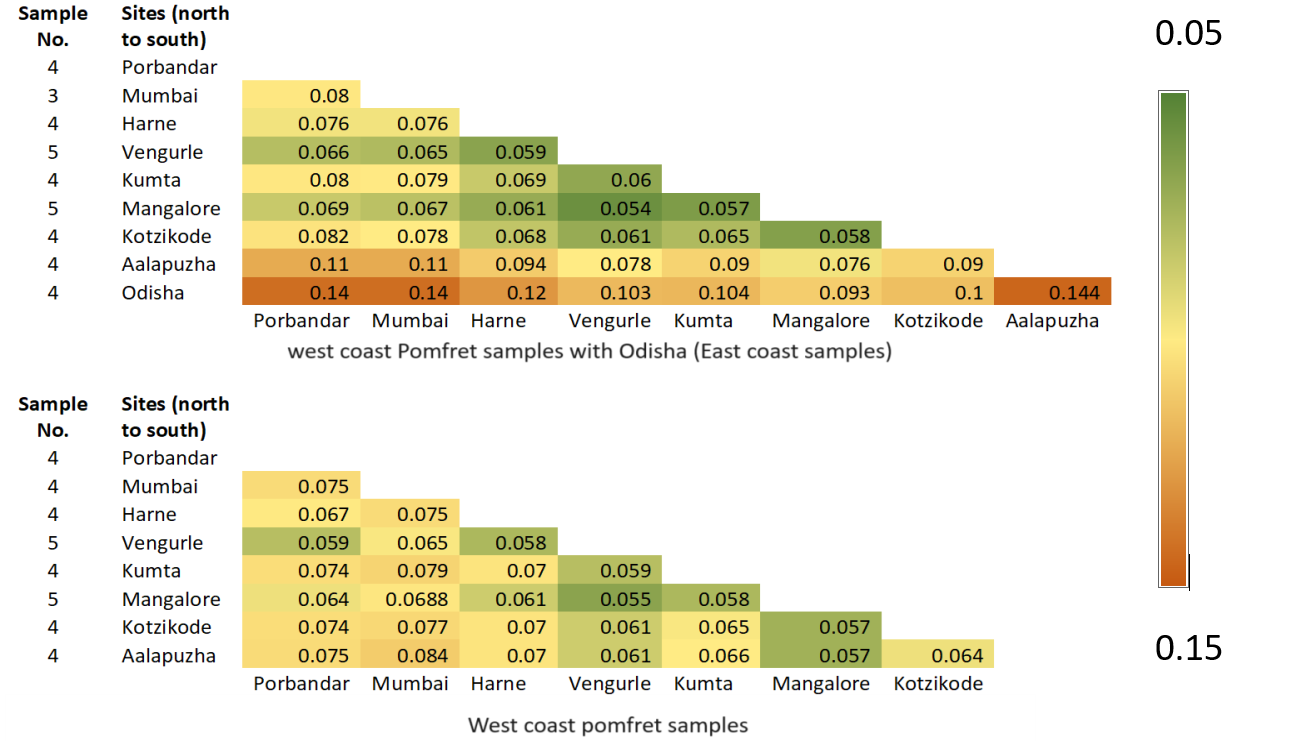  0.0  0.1  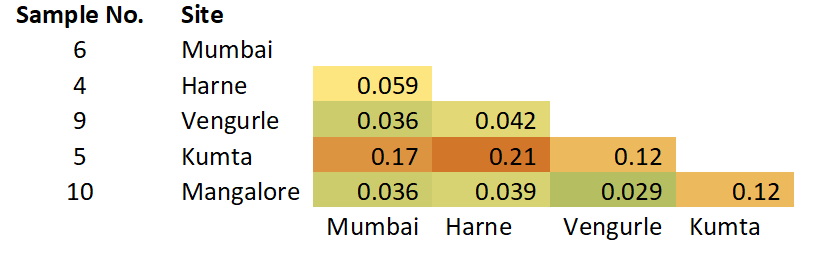  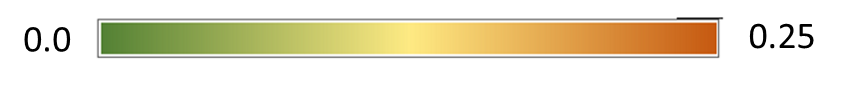 |

Table S8. Likelihood distributions for K values for all the species

| Species | Spatial extent | K1 | K2 | K3 | K4 | K5 | K6 | K7 | K8 |
| --- | --- | --- | --- | --- | --- | --- | --- | --- | --- |
| *Rastrelliger kanagurta* | West coast and odisha (east coast) |  | 4.029 | 3.271 | 0.197 | 2.326 | 0.302 | 0.739 | 1.111 |
| *Rastrelliger kanagurta* | West coast only |  | 3.165 | 2.206 | 0.052 | 2.624 | 1.349 | 0.134 | - |
| *Chirocentrus nudus* | West coast only |  | 0.140 | 4.023 | 0.623 | 1.349 | 0.505 | 0.789 | 0.912 |
| *Epinephelus diacanthus* | West coast only |  | 0.586 | 0.307 | 0.589 | 3.284 | 0.304 | 1.043 | 0.011 |
| *Trichiurus lepturus* | West coast and odisha (east coast) |  | 9.4 * 10^8 | 2.810 | 3.996 | 0.067 | - | - | - |
| *Trichiurus lepturus* | West coast only |  | 1.954 | 0.311 | 2.344 | - | - | - | - |
| *Nemipterus japonicus* | West coast and odisha (east coast) |  | 2.7 * 10^11 | 2.517 | 2.458 | 0.582 | 1.977 | - | - |
| *Nemipterus japonicus* | West coast only |  | 2.846 | 1.129 | 1.651 | 0.783 | 0.018 | 0.370 | - |
| *Pampus argenteus* | West coast and odisha (east coast) |  | 3.1 * 10^9 | 0.531 | 2.989 | 1.091 | 1.915 | 0.111 | 0.143 |
| *Pampus argenteus* | West coast only |  | 190.5 | 0.488 | 1.705 | 1.166 | 0.193 | 0.655 | - |
| *Sardinella longiceps* | West coast only |  | 1.182 | 0.602 | 1.531 | - | - | - | - |
| *Scoliodon laticaudus* | West coast only |  | 2.806 | 521.3 | 0.497 | 14.423 | 0.556 | 1.050 | - |
| *Sillago sihama* | West coast only |  | 1.5 * 10^9 | 3.474 | 9.690 | - | - | - | - |

**Supplementary material – section 4**

**Scripts of all the codes used in genetic analysis**

We have conduced same set of analysis for nine different species. In this section we have provided script of all the steps in pipeline using representative samples/files. Based on representative samples/files one can extrapolate code for all nine species by simply changing file and folder names

Please note that all the instructions are written in the black font whereas the script to type in consol is typed in italics and blue font.

Subsection 1: Processing ddRAD data for SNP calling

A) When reference genome is available

**Step 0)** how the data looks -

File for each sample which have come after sequencing and demultiplexing would look something like this

pd1_S88_R1_001.fastq.gz

pd1_S88_R2_001.fastq.gz

**Step 1)** trimming - This is done for getting rid of any adapter or index sequences attached to the reads even after demultiplexing

Required files and directories

| File or directory name | Their function | Code to generate those files |
| --- | --- | --- |
| Trimmed_reads folder | This folder will store the trimmed reads that you will generate | *mkdir Trimmed_reads* |
| SampleNames file | In this text file the names of all the samples are stored  They should be listed in this format  pd1_S88  pd2_S89  I usually make this file in notepad and then copy paste the content in newly created file | *vim SampleNames*  vim is a command use to create and edit text files |

After making Trimmed_reads folder and SampleNames file we will make text files with codes which will talk to each other

1) TrimmingFiles

*vim TrimmingFile*

Then use following code

*#$ -pe orte 16*

*java -jar /softwares/trimmomatic/bin/trimmomatic.jar PE -threads 16 -phred33 ~/pomfretwork/pomfret/Sample_R1_001.fastq.gz ~/pomfretwork/pomfret/Sample_R2_001.fastq.gz ~/pomfretwork/pomfret/Trimmed_reads/Sample_Trimmed_1P.fastq.gz ~/pomfretwork/pomfret/Trimmed_reads/Sample_Trimmed_1U.fastq.gz ~/pomfretwork/pomfret/Trimmed_reads/Sample_Trimmed_2P.fastq.gz ~/pomfretwork/pomfret/Trimmed_reads/Sample_Trimmed_2U.fastq.gz ILLUMINACLIP:/softwares/trimmomatic/adapters/TruSeq3-PE.fa:2:30:10:2:keepBothReads AVGQUAL:30 LEADING:3 TRAILING:3 MINLEN:30*

2) Submitjob

*vim Submitjob*

Then use following code

*while read Sample*

*do*

*cp TrimmingFile TrimmingFile_${Sample}*

*sed -i "s/Sample/${Sample}/g" TrimmingFile_${Sample}*

*qsub TrimmingFile_${Sample}*

*done < SampleNames*

3) To activate Submitjob code we have to execute follwoing command

*chmod +x Submitjob*

4) Running the Submitjob (step 2, Trimming the files)

After chmod +x now Submitjob code is ready to run

Run it using following command

*./Submitjob*

**Step 2)** Indexing reference genome

*/softwares/bwa-0.7.17/bwa index E.moara_RefGenome*

**Step 3)** alignment - in this step we will align trimmed reads which we made in step one with the reference genome and creating bam files for each samples

1) First make a folder where aligned files will sit. Make sure this folder is in the same folder where the SampleNames file is saved for the trimming procedure. Since bothe procedures use same file it will be easy to use it

*mkdir alignment*

After making alignment folder make a text file for alignment script

*vim Alignment*

populate the alignment script text file with this code

*#$ -pe orte 24*

*/softwares/bwa-0.7.17/bwa mem -M -t 24 -R "@RG\tID:Sample\tPL:ILLUMINA\tSM:Sample" ~/grouper_trim/grouperwork/RefGenome/E.moara_RefGenome ~/grouper_trim/grouperwork/grouper/Trimmed_reads/Sample_Trimmed_1P.fastq.gz ~/grouper_trim/grouperwork/grouper/Trimmed_reads/Sample_Trimmed_2P.fastq.gz| /softwares/samtools-1.9/samtools view -Sb -F4 - | /softwares/samtools-1.9/samtools sort -o ~/grouper_trim/grouperwork/grouper/alignment/Sample_sorted.bam*

*/softwares/samtools-1.9/samtools index ~/grouper_trim/grouperwork/grouper/alignment/Sample_sorted.bam*

2) Create a submitjob-alignment script

*vim submitjob-alignment*

use the following code to populate that script

*while read Sample*

*do*

*cp Alignment Alignment_${Sample}*

*sed -i "s/Sample/${Sample}/g" Alignment_${Sample}*

*qsub Alignment_${Sample}*

*done < SampleNames*

4). To activate submitjob-alignment code we have to execute follwoing command

*chmod +x submitjob-alignment*

5) Running the submitjob-alignment (step 2, Trimming the files)

After chmod +x now Submitjob code is ready to run

Run it using following command

*./Submitjob*

**Step 4)** indexting before variant calling - it is done because for variant calling you need a .fai file

To unzip the .gz code we can use command gunzip

*gunzip GCA_000697985.1_PamArg1.0_genomic.fna*

Then we will proceed with indexing

*/softwares/samtools-1.9/bin/samtools faidx GCA_000697985.1_PamArg1.0_genomic.fna*

After this step there are two ways by which one can do variant calling

A) using freebays
B) using STACKS

**Step 5A)** variant calling – using freebays

1) Making a text file with all the list of bam files –

*vim list_bam_mackerel*

2) Creating a folder for freebayes_output

*mkdir freebayes_output_withodisha*

3) Make a text file for following variant calling code and run it using qsub

*#$ -S /bin/bash*

*#$ -cwd*

*#$ -V*

*#$ -pe orte 64*

*#$ -q all.q*

*/softwares/freebayes1.1/bin/freebayes -f ~/mackerelwork/mackerel/mackerel_refgenome/Mackerel_GCA_024605255.1_ASM2460525v1_genomic.fna --genotype-qualities -L ~/mackerelwork/mackerel/Mackerel/list_bam_mackerel > ~/mackerelwork/mackerel/Mackerel/freebayes_output_withodisha/mackerel_freebayse_all.vcf*

**Step 5B)** variant calling – using STACKS

1) Making a population map file –

*vim sampleMap95_bigeye*

make sure that the samples and population identity should be written in following way

obig3_S213_L001_sorted odisha_big

obig4_S206_L001_sorted odisha_big

AM7_S74_sorted allapuzha

KM1_S65_sorted Kotzhikode

KM2_S66_sorted Kotzhikode

2) Creating a folder for freebayes_output

*mkdir ref_map_pl_output_odisha_big*

1) Make a text file for following variant calling code and run it using qsub

*$ -S /bin/bash*

*#$ -cwd*

*#$ -V*

*#$ -pe orte 64*

*#$ -q all.q*

*/softwares/stacks2.3/bin/ref_map.pl -T 8 -o /home/uramakri/msgangal/mackerelwork/mackerel/Mackerel/ref_map_pl_output_odisha_big/ --popmap /home/uramakri/msgangal/mackerelwork/mackerel/Mackerel/sampleMap95_bigeye --samples /home/uramakri/msgangal/mackerelwork/mackerel/Mackerel/alignment/ -X "populations:--fstats"*

Subsection 1: Processing ddRAD data for SNP calling

B) When reference genome is NOT available

**Step 0)** how the data looks -

File for each sample which have come after sequencing and demultiplexing would look something like this

pd1_S88_R1_001.fastq.gz

pd1_S88_R2_001.fastq.gz

**Step 1)** concatenate files using process radtags tool in stacks

First make a output directory

mkdir *concat-wolf_june2023*

Make a text file for following process_radtags code and run it using qsub

*#$ -S /bin/bash*

*#$ -cwd*

*#$ -V*

*#$ -pe orte 64*

*#$ -q all.q*

*/softwares/stacks2.3/bin/process_radtags -P -p /home/uramakri/msgangal/wolfwork/wolf_june2023/ -o /home/uramakri/msgangal/wolfwork/concat-wolf_june2023/ -q -c -r -t 95 --renz_1 sphI --renz_2 mluCI --disable_rad_check*

**Step 2) Copy files without rem files**

*mkdir concat-grouper-norem*

*cp /home/uramakri/msgangal/concat-grouper/{AG1_S36_R1_001.1.fq.gz,AG1_S36_R2_001.2.fq.gz, ….. (all file names)} /home/uramakri/msgangal/concat-grouper-norem*

**Step 3) change the file names**

as denovo_map.pl command only understand file r1 files as .1.fq.gz format. So we have to remove other suffixes in file names for eg. In the file name written after this statement the highlighted part should not exist AG_D20_R1_001.1.fq.gz

change in file name code

*mv AG_D20_R1_001.1.fq.gz AG_D20.1.fq.gz*

Similarly change names of all the fq.gz files

**step 4) Running denovo_map.pl**

First make a directory to store denovo bam files

mkdir *denovo_bam_wolf_nudus_westcoast*

Then make list of samples

vim *sampleMap_nudus_westcoast*

Make sure in sample files samples are arranged like this

WM1E2_S129 mumbai

H2W7_S227 kumta

PW2_S218 porbandar

Make a text file for following denovo_map.pl code and run it using qsub

*$ -S /bin/bash*

*#$ -cwd*

*#$ -V*

*#$ -pe orte 64*

*#$ -q all.q*

*/softwares/stacks2.3/bin/denovo_map.pl -T 8 -M 5 -n 5 -o /home/uramakri/msgangal/wolfwork/denovo_bam_wolf_nudus_westcoast --samples /home/uramakri/msgangal/wolfwork/norem --popmap /home/uramakri/msgangal/wolfwork/sampleMap_nudus_westcoast –paired*

**Step 5) creating a vcf file using tools populations from STACKS**

First make a directory to store vcf files

mkdir *vcf_nudus_jan2024*

Then create a text file of population map

*vim sampleMap_nudes*

Make sure that in map file populations are arranged like this samples are arranged like this

S2W5_S153	mangalore

WD3_S136	harne

WM1E2_S129	mumbai

Make a text file for following populations code and run it using qsub

*$ -S /bin/bash*

*$ -cwd*

*#$ -V*

*#$ -pe orte 64*

*#$ -q all.q*

*/softwares/stacks2.3/bin/populations -P /home/uramakri/msgangal/wolfwork/denovo_bam_wolf_nudus_westcoast --popmap /home/uramakri/msgangal/wolfwork/sampleMap_nudus --vcf -t 8 -O /home/uramakri/msgangal/wolfwork/vcf_nudus_jan2024/*

Subsection 2: Filtering varients using VCFtools for SNP calling

Once we have vcf files with us we did filtering using VCF tools. The example of the filtering is shown with the help of filtering file for removing indels

/softwares/vcftoolsV13/bin/vcftools --vcf mackerel_freebayse_all.vcf --remove-indels --out mackerel_freebayse_all_NoIndel --recode

For filtrations we took help of vcftools manual <https://vcftools.sourceforge.net/man_latest.html>

The filtration sequence of variants to identify Single Nucleotide Polymorphisms (SNPs)n is explained in supplementary material section 3 table 7

For checking missing data at any step, we first used this command

/softwares/vcftoolsV13/bin/vcftools --vcf mackerel_freebayse_all_NoIndel_minQ30.recode.vcf --missing-indv --out mackerel_freebayse_all_NoIndel_minQ30_missingIndv

This creates a .imiss file which can be checked to see percent missing data in each sample to filter out samples with high missing data

*less mackerel_freebayse_all_NoIndel_minQ30_missingIndv.imiss*

Subsection 3: Codes for Analysis of SNP data

A) Principal Component Analysis (PCA)

We use software PLINK for principle component analysis

Once the filtered vcf file is ready for analysis we make .ped and .map files using this command

*/softwares/vcftoolsV13/bin/vcftools --vcf nudus_westcoast_noindels_minGQ3 0_minDP10_minmax2_noBad_hwe005_mac3_MM90.recode.vcf –plink –out nudus*

Then using. ped and .map files we will make PCA plot

*/softwares/plink/plink --ped nudus.ped --map nudus.map --pca --allow-extra-ch r --out nudus_westcoast --autosome-num 95*

After this command we will get two files. First is file with eigenvalues which can be accessed using following command

*less nudus_westcoast.eigenval*

We calculate proportion of variation explained by first principal component by taking ratio of its eigenvalue to sum of all eigenvalues. For subsequent principle components we calculated variation explained by them using similar procedure

Another file is of corresponding eigenvectors which can be accessed by following command

*less nudus_westcoast.eigenvec*

We copied the eigen vector file into excel. Named the columns representing individual principal components. We also named rows with appropriate sample name and population name. Then we exported that data in R. Plotting of principle component plots was done in R

Plotting PCA in R

First to get colours we loaded a library RColorBrewer

*> library(RColorBrewer)*

To depict north south gradient we created a colour map using colours in R. With dark red colours for the northernmost side and converting it slowly to Dark blue colour at the southern and even darker blue to east cost site

*> color_mapping <- c(a_Porbandar = "#A50026", b_Surat = "#D73027", c_Mumbai = "#F46D43", d_Harne = "#FDAE61", e_Vengurle = "#FEE090", f_Goa = "#FFFFBF", g_Kumta = "#E0F3F8", h_Mangalore = "#ABD9E9", i_Kozhikode = "#74ADD1", j_Alappuzha = "#4575B4", k_Odisha = "#313695")*

Then we brought the eigenvector data in R console by copying the eigenvector metrix created in previous step on clipboard

*> wolf_westcoast <- read.delim("clipboard")*

We assigned locations on the wolf_westcoast matrix with the corresponding colour template created in previous step

*> point_colors <- color_mapping[wolf_westcoast$site]*

Then we plotted the first and second principle components using the colour template to indicate samples coming from individual locations. We also mentioned the individual variation explained by each principle component in the script

*> plot(x2~x1, data=wolf_westcoast, pch = 16, cex = 3, col=point_colors, cex.lab = 2, xlab = "pc1 (6.8 % )", ylab = "pc2 (5.8 %)")*

B) Admixture analysis

For admixture analysis we need to convert vcf file into .bed file. We can use .map and .ped files made with help of plink for admixture analysis. to make .ped file we use following code

*/softwares/plink-1.07-x86_64/plink --file out --make-bed --out nudus_westcoast*

After making .ped file, we make a directory to store all the admixture files (.Q and .out files)

*mkdir admixure_loops*

Now we are ready to run admixture analysis using software Admixture. We will first create a script file (text file) to house admixture code. In the script file we use following code. In this particular code we have taken 6 different values of K and we have ran the code 10 times (r=10) for each value of K (number of pre-defined populations)

*#!/bin/bash -l*

*#$ -S /bin/bash*

*#$ -cwd ##Move to current working directory*

*#$ -V*

*#$ -q all.q*

*#loops for running admixture at each K for 10 runs per k*

*#First, defining the file name as prefix so that I will not have to type the file name each time*

*prefix=nudus_westcoast*

*#first loop to run admixture r number of times for each value of K*

*for r in {1..10}*

*do*

*#second loop to run admixture for different values of K*

*for K in {1..6}*

*do*

*# --cv option will do cross-validation to find the best K value from the output log file*

*# tee will read the standard output of the run and write it in the logfile*

*/home/uramakri/msgangal/Softwares/dist/admixture_linux-1.3.0/admixture --cv -s ${RANDOM} ~/wolfwork/vcf_nudus_jan2024/${prefix}.bed ${K} | tee ~/wolfwork/vcf_nudus_jan2024/admixure_loops/log${prefix}K${K}r${r}.out*

*#As the output file name is chosen by default, after each run for a K and r value, the output file needs to be renamed*

*mv ~/wolfwork/vcf_nudus_jan2024/${prefix}.${K}.Q ~/wolfwork/vcf_nudus_jan2024/admixure_loops/${prefix}.K${K}r${r}.Q*

*done*

*done*

Here each .Q file contains probability of each sample to be associated with different populations which are assigned according to value of K.

We will compile all 10 replicates for each value of K by first making directory for certain value of K

*mkdir nudus_k1*

And then copy all the replicates to that directory

*scp nudus_westcoast.K1* /home/uramakri/msgangal/wolfwork/vcf_nudus_jan202 4/admixure_loops/nudus_k1*

And then making a zip file of it

*zip -r nudus_k1 nudus_k1*

once we did this for all the values of K then we will combine all these zip files into one larger directory which we will then zip. We will use this file in online tool CLUMPAK to run admixture visualisation

*mkdir nudus_zip*

*scp *zip ./nudus_zip*

Apart from zip file we also require text file which identifies the location for each sample. In this file we have to keep the same sequence of samples as it was present in vcf file. The file should comprise of just list of locations and it should look something like this

Mangalore

Mangalore

Harne

Mumbai

Kumta

Porbandar

Porbandar

Now the zip file and population file is used in CLUMPAK to give a visualisation of probabilities of individual samples in each location belonging to a population. In CLUMPAK output one colour represents one population of genetically similar individuals.

Calculating cross validation error

For calculating cross validation error for each value of K we use software admixture and a single code. In the given example we have written a script to calculate cross validation error for values of K ranging from 1 to 6

*for K in 1 2 3 4 5 6; do /home/uramakri/msgangal/Softwares/dist/admixture _linux-1.3.0/admixture --cv nudus_westcoast.bed $K | tee log${K}.out; done*

C). fixation index (FST)

We calculated fixation index with the help of software STACKS using tool populations

For that first we have to create a population map file

*vim nudus_westcoast_popmap*

The samples in that file should be arranged in following way

S2W4_S152	Mangalore

S2W5_S153	Mangalore

WD3_S136	Harne

WM1E2_S129	Mumbai

We also need a director to host output files

*mkdir fst_nudus_westcoast*

We can now run the FST script

*/softwares/stacks2.3/bin/populations -V nudus_noindel_minGQ30_minDP10_minmax2_NoOdisha_hwe005_mac3_MM90.recode.vcf -M nudus_westcoast_popmap -O fst_nudus_westcoast --fstats*

The summery of the output shows pairwise fst between each population

1. L50 - length at which a fish has a 50% probability of being retained by a specific fishing gear
2. Lm - length at which 50% of the individuals in a fish population reach sexual maturity
